# Understanding genetic and environmental complexity in phenotypic resemblance across relatedness in population-scale data

**DOI:** 10.64898/2026.10.18.759872

**Authors:** Mariadaria K. Ianni-Ravn, Andy W. Dahl, John Novembre

**Affiliations:** Department of Human Genetics, University of Chicago, Chicago, IL 60637; Division of Genetic Medicine, University of Chicago, Chicago, IL 60637; Department of Ecology and Evolution, University of Chicago, Chicago, IL 60637

## Abstract

A fundamental feature of a trait’s variation in a population is how its phenotypic similarity scales with genetic relatedness. Under a strictly additive model of genetic architecture, phenotypic similarity increases linearly with genetic relatedness, with a slope equal to the heritability; this underpins approaches to heritability estimation from sibling IBD regression to Haseman–Elston regression and GREML. Empirically, however, the relationship is not linear for many traits (Kemper et al., 2021). Biobank-scale measurements make fine-grain observations of the phenotypic covariance versus genetic relatedness relationship increasingly feasible, motivating new theory to understand these new empirical observations. Here we evaluate how three processes might introduce non-linearities to the relationship: imperfect tagging of causal variants, assortative mating, and multi-generational shared environments. Through theoretical modeling and simulation, we find each process acts within pedigree classes differently, introducing non-linearities through impacts on the local slope and intercept of phenotypic covariance on genetic relatedness: imperfect tagging lowers the slope among distant relatives, shared environments elevates the intercepts for closer relatives, and assortative mating changes both the slope and intercept. Within these models, we explore a variety of scenarios, finding cases where the local slope of covariance on relatedness values among unrelated pairs versus among siblings can reach nearly three times the raw heritability, and building insight to where heritability estimates are likely to be biased. We then examine these patterns across a range of traits in the UK Biobank, with a supplementary analysis in All of Us. The shape of the relationship varies widely across traits, with profiles exhibiting features found under each of the theoretical mechanisms we explored. Overall, we conclude disentangling these multiple overlaid processes will be necessary to fully understand how phenotypic similarity increases with genetic relatedness in large cohorts.

## 1 Introduction

For a quantitative trait with a genetic component, closer relatives are expected to resemble each other more than distantly related individuals. This is an intuitive result that is visible in family resemblances across species, and has underpinned the concept of heritability for over a century (Rice, 2008). Yet the full shape of this relationship across the relatedness spectrum is a richer object than a summary such as heritability, as it carries the joint imprint of all the forces involved in trait variation.

A fundamental result predicts more specifically what we should expect: for a trait subject only to additive polygenic variation, phenotypic covariance between a pair of individuals increases linearly with their genetic relatedness at causal variants (Fisher, 1919). When the phenotype is standardized to unit variance, the slope of this linear relationship equals the narrow-sense heritability, *h*^2^. The empirical slope of the regression of phenotypic covariance on genetic relatedness is known as the Haseman–Elston (HE) estimator of heritability (Haseman and Elston, 1972). Beyond HE regression, many contemporary approaches including genomic restricted maximum likelihood (GREML) and related variance-component methods similarly assume a linear relationship between genetic and phenotypic covariance, and leverage it for inference.

Despite how fundamental the relationship between phenotypic covariance and genetic relatedness is, relatively few studies have examined its shape empirically across the full range of genetic relatedness. The most comprehensive analysis to date is Kemper et al. (2021), who used UK Biobank (UKB) data to examine phenotypic covariance across many quantitative traits and a broad range of genetic relatedness, from twins to unrelated individuals, which was measured with a GRM at common SNPs. Strikingly, the relationship between phenotypic covariance and genetic relatedness departed from linearity for a range of quantitative traits, three of which are shown in Figure 1. Specifically, the data revealed two striking patterns: firstly, the slope of phenotypic covariance on genetic relatedness was substantially steeper among close relatives (genetic relatedness above about 0.05) than among nominally unrelated pairs (below about 0.02); secondly, among distant relatives, at intermediate genetic relatedness (0.02–0.05), phenotypic covariance showed an abrupt increase inconsistent with a simple linear relationship predicted under an additive trait architecture.

**Figure 1:**
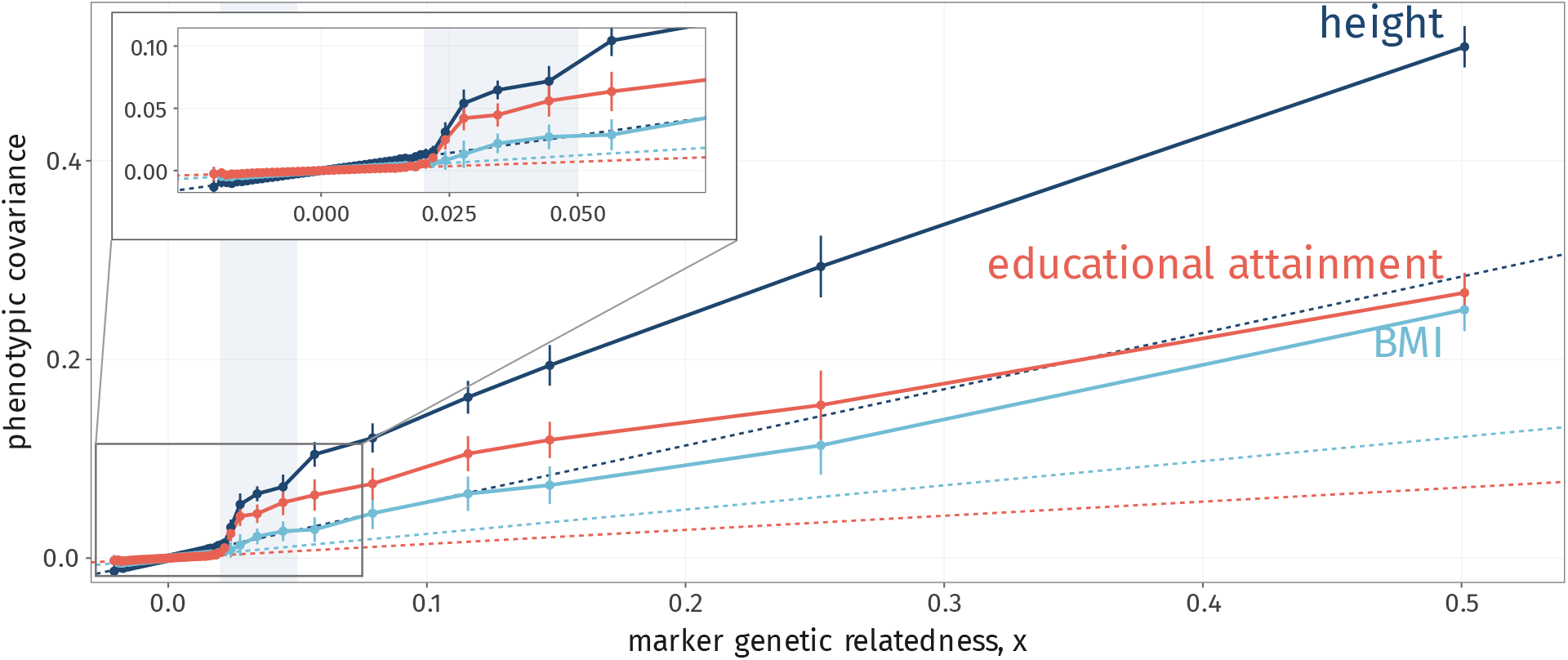
Phenotypic covariance against genetic relatedness (GRM) in UK Biobank for height, educational attainment and BMI, redrawn from Kemper et al. (2021). Intervals, 95%; dashed lines, slope equal to the SNP heritability. Inset, the low-relatedness region.

These patterns relate to the long-standing phenomenon of missing heritability: family-based heritability estimates are typically higher than those from genome-wide SNP data in unrelated individuals (Manolio et al., 2009). The Kemper et al. observations are consistent with this, with a steeper slope of phenotypic covariance on genetic relatedness among close relatives than among unrelated pairs, suggesting systematic differences in heritability captured by common SNPs across the relatedness spectrum.

Kemper and colleagues investigated whether the empirical patterns they observed could be explained by imperfect tagging of causal variants, also called imperfect linkage disequilibrium (LD). This describes the failure of observed marker variants to perfectly predict causal variation in unrelated samples. They simulated quantitative phenotypes under varying assumptions about the allele-frequency distribution of causal variants and found that when causal variants were rare and marker variants common, simulations reproduced the accelerated increase in phenotypic covariance among distant relatives (genetic relatedness 0.02–0.05) observed in their data. This arises because the predictor in HE regression is ideally the covariance at causal variants, but in practice this is approximated using observed genetic variants.

The quality of this approximation varies across the relatedness spectrum: for unrelated pairs, a GRM captures covariance only at observed SNPs and variants in LD with them; when causal variants are rare, population-level LD provides poor information about rare variants. At high genetic relatedness, by contrast, segments shared identical-by-descent (IBD) may increase tagging of rare variants. This suggests the efficiency with which markers tag causal variants should vary systematically across relatedness (de los Campos et al., 2013; Eynard et al., 2015). Nevertheless, we currently lack a theoretical framework for understanding how tagging efficiency varies across the relatedness spectrum in relation to genetic architecture and marker properties.

Another potential influence on the phenotypic–genotypic covariance relationship is shared environment between relatives. Environmental correlations may extend beyond nuclear families into nominally unrelated pairs through several mechanisms, such as through cultural transmission of behaviors and preferences (e.g., dietary practices, physical activity), intergenerational wealth transfer affecting a broader suite of outcomes, such as education and health, and geographic clustering exposing relatives to similar air quality, green space, and other spatially varying conditions. We label these factors multi-generational shared environments (MGSEs) to emphasize their persistence across extended genealogical distances.

Early practitioners of genetic studies recognized that such environmental confounding could inflate heritability estimates. This concern, particularly following reports of high familial IQ heritability, motivated formal models of cultural inheritance in the 1970s and 1980s (Cavalli-Sforza and Feldman, 1973, 1981; Cloninger et al., 1979), which showed that Fisherian patterns of resemblance could arise from cultural factors alone. In extended pedigrees, environmental correlations are sometimes modeled explicitly via family-level effects, yet it remains unclear how best to represent these correlations given their likely variation across exposures and phenotypes. Notably, Zaitlen and colleagues found that heritability estimates vary systematically across pedigree relationship classes, which is a pattern consistent with unmodeled shared environmental structure (Zaitlen et al., 2013).

Assortative mating (AM) — the tendency of individuals to select partners who are phenotypically similar to them in some way — is thought to be another important force affecting the relationship between phenotypic covariance and genetic relatedness for many traits. Phenotypic correlations between mates have been documented across diverse traits, ranging from ~0.2 for anthropometric measures (height, body mass index (BMI)) (Robinson et al., 2017) to ~0.5 for educational attainment (Horwitz et al., 2023). Assortative mating’s impact on heritability estimates varies across the relatedness spectrum (Yengo and Visscher, 2018; Kemper et al., 2021), complicating the interpretation of this relationship. Furthermore, even mild assortative mating, acting together with cultural transmission from parents to offspring, can strongly inflate heritability estimates (Border et al., 2026). A further consequence is the induction of gene–environment correlations: when environmental influences are also transmitted from parents to offspring, assortment correlates trait-increasing alleles with trait-increasing environments, coupling components that would otherwise be independent (Rice et al., 1978; Cloninger et al., 1979; Border et al., 2026).

Phenotypic similarity across the genetic relatedness spectrum is therefore a fundamental emergent pattern of interest. However, several questions remain unanswered: how do quantitative genetic processes shape the relationship between phenotypic covariance and genetic relatedness across trait architectures? What can we learn about individual traits by examining these covariances at fine-grained resolution, and how does the relationship vary across traits and datasets empirically? Addressing these questions requires a more refined understanding of this relationship and the mechanisms operating across scales of relatedness.

Beyond querying fundamental processes shaping human phenotypic variation, the answers to these questions have practical implications, since they affect statistical genetic summaries of this relationship, such as the Haseman–Elston estimates of heritability. In particular, we must understand these patterns in order to interpret what this statistic recovers in practice, where it is most vulnerable to confounding, and over which ranges of relatedness it can be meaningfully read as a heritability in the traditional sense. This relationship has implications for phenotypic prediction, as the excess resemblance between close relatives could surmisably be leveraged in genomic prediction models by upweighting.

In this work, we develop a unified theoretical framework to address these questions. We combine analytical modeling with simulations to characterize how imperfect tagging of causal variants, multi-generational shared environments, and assortative mating shape phenotypic covariance across the full relatedness spectrum. By explicitly linking these three mechanisms to pedigree structure and the empirical distribution of relatedness in finite samples, we show how non-linear patterns arise and how their signatures differ. This provides a principled basis for interpreting deviations from linearity and understanding their consequences for heritability estimation across the relatedness spectrum.

We then see how these predictions play out empirically using fine-scale binning of relatedness across complex traits and exposures in the UK Biobank. We observe substantial heterogeneity in covariance patterns across phenotypes, some of which resemble the signatures the theory predicts, showcasing the broad variety of trait architectures visible in the Biobank.

Our findings support that, for a wide range of traits, a simple linear model does not describe the data well. Such patterns might confound naive slope-based heritability estimates. Overall, these covariance spectra point to the power of fine-grained relatedness-based analyses to dissect sources of familial resemblance.

## 2 Results

### 2.1 Phenotypic covariance as an average over pedigree classes

At any observed level of genetic relatedness, a sample contains a mixture of different pedigree relationship classes. The phenotypic covariance is therefore an average over this mixture:

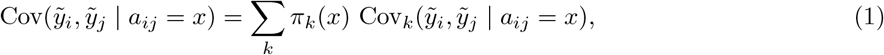

where 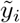 and 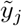 denote the standardized phenotypes of individuals *i* and *j, a*_*ij*_ is their genetic relatedness, with value *x* (Kemper et al. 2021 *write π* for it; we use *π* for class proportions), *k* indexes pedigree relationship class, and *π*_*k*_(*x*) = P(*k* | *a*_*ij*_ = *x*) is the probability that a pair with genetic relatedness *x* belongs to class *k*. Cov_*k*_ denotes a covariance among pairs of class *k*. Since the phenotypes are standardized to mean zero, each covariance equals the expected cross-product 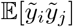 under the same conditioning.

The sum on the right-hand side has two components: firstly, *π*_*k*_(*x*) weights the contribution of each pedigree class in the cohort. We will begin by focusing on these weights, which describe the shifting mixture of relatives across genetic relatedness values. The second term is the expected phenotypic covariance, conditioned on pedigree class *k* and genetic relatedness value *x* within that class. In the sections that follow, we characterize how imperfect tagging, multi-generational shared environments, and assortative mating each affect this expectation, and how these signatures interact with the shifting mixture of relatives to shape empirical patterns of covariance.

### 2.2 The shifting mixture of pedigree relatives across genetic relatedness

Relatives of different degrees have characteristic but overlapping distributions of realized genome sharing. These distributions are centered on expected values (e.g., 0.5 for full siblings, 0.125 for first cousins) determined by the number of meioses connecting them, but vary due to the random processes of Mendelian segregation and recombination. As a consequence, relatives of increasingly distant classes have larger coefficients of variation and skewness in their distributions, resulting in substantial overlap at low levels of genetic relatedness (Hill and Weir, 2011), as seen in Figure 2a. This overlap means that at any observed value of genome-wide genetic relatedness, a sample may contain a mixture of many different pedigree classes.

**Figure 2:**
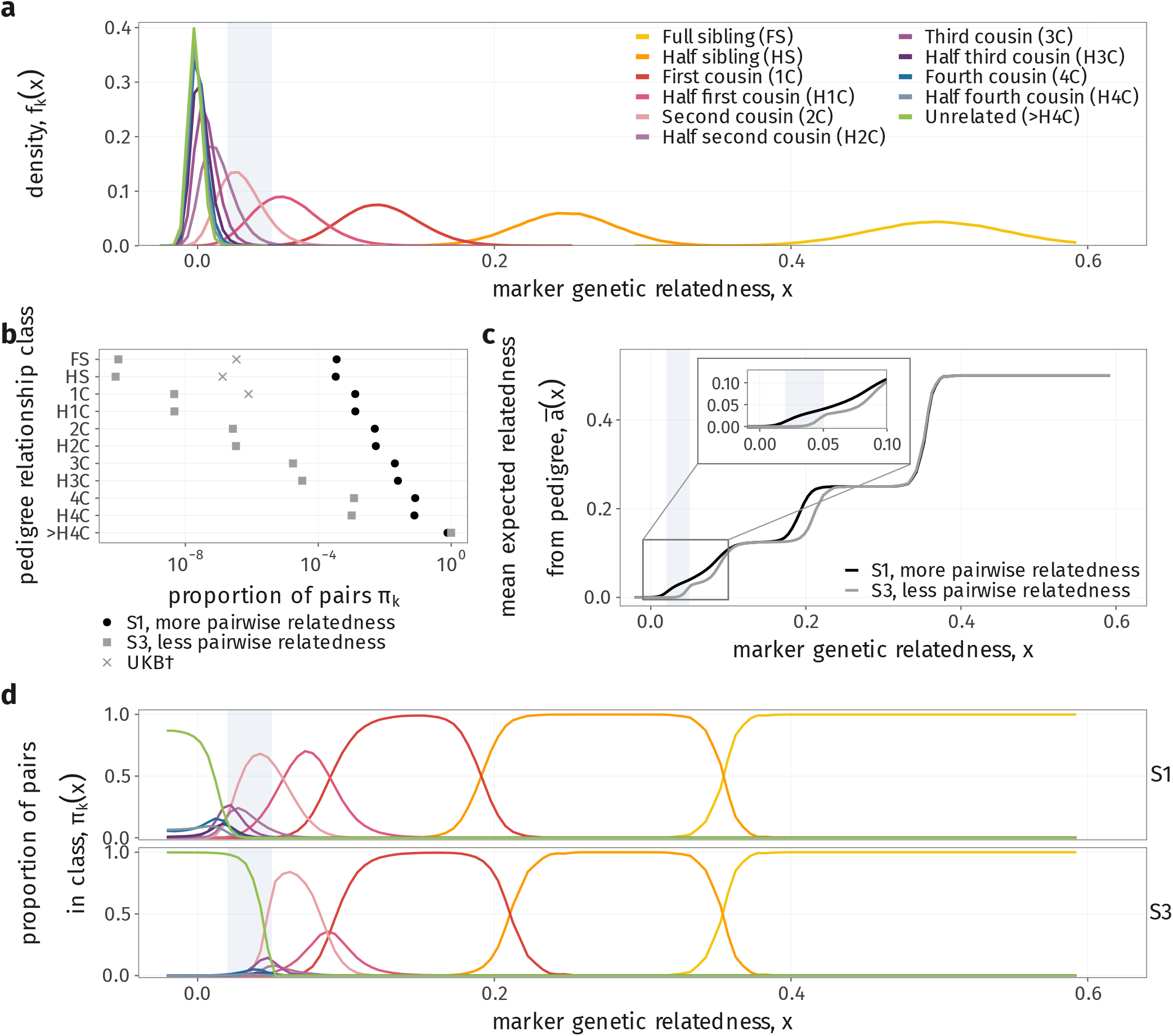
Pairs at the same genetic relatedness are a mixture of pedigree classes, and the mixture depends on how the sample is drawn. (a) Distribution *f*_*k*_(*x*) of genetic relatedness *x* in each pedigree class, from full siblings (FS) to half fourth cousins (H4C), and among unrelated pairs (*>*H4C); 50 replicate forward-in-time simulations (*N* = 5,000, 20 generations). (b) Proportion of pairs *π*_*k*_ in each class under S1 (the simulated population) and S3 (S1 proportions cubed, renormalized). Crosses, first-to third-degree proportions in UK Biobank^*†*^. (c) Mean expected pedigree relatedness *ā* (*x*) = ∑_*k*_ *π*_*k*_(*x*) *ā*_*k*_ under S1 and S3. Inset, the low-relatedness region. (d) Class composition *π*_*k*_(*x*) = Pr(*k* | *x*) under S1 (top) and S3 (bottom); colors as in (a). Gray band (0.02 ≤ *x* ≤ 0.05): composition shifts from unrelated pairs and distant cousins to second cousins. ^*†*^Inferred by KING in UK Biobank (Manichaikul et al., 2010; Zhang et al., 2025).

In principle, a population contains an enormous space of possible pedigree relationships. For our purposes, it is enough to concentrate on a narrower set of these relationships to capture the covariance structures relevant to our study—specifically, pairs of individuals in the same generation who share common ancestors via one or two lineages. Throughout this work, we index these pedigree classes by *k* and represent relationships up to the level of half fourth cousins (with expected relatedness 2^−10^), treating individuals who are more distantly related as effectively “unrelated”. For details on how we treat these classes in the simulations described later, see Supplementary Section S8.1.

A key question then becomes: at a given level of measured genetic relatedness, what is the underlying composition of pedigree relationships? We characterize this mixture through *π*_*k*_(*x*), the probability that a pair of individuals with observed genetic relatedness *x* belongs to pedigree class *k*:

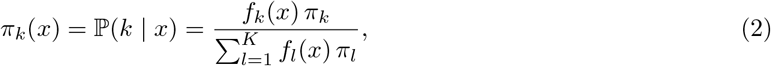

where *f*_*k*_(*x*) is the within-class distribution of genetic relatedness and *π*_*k*_ is the mixture proportion of pedigree class *k* in the sample, for *K* pedigree classes in total.

The *π*_*k*_ proportions are determined by sample composition. Population-based cohorts like the UK Biobank contain far more distantly related pairs than close relatives, but this distribution may be skewed by recruitment patterns, geographic clustering, or demographic factors. For instance, in expanding populations we expect to sample fewer close relatives, while regional sampling biases may over-represent nearby kin. Estimating these proportions empirically is challenging, as kinship inference methods typically can only reliably detect relatives up to the fourth or fifth degree (Manichaikul et al., 2010).

To explore how sample composition affects our results, we compared two sampling schemes, which differ in the proportion *π*_*k*_ of pairs in each pedigree class. The first, S1, is the composition of our forward-in-time simulated population (*N* = 5,000). The second, S3, raises the S1 proportions to the power of three and renormalizes them (each scheme is named after its power). This shifts the mixture toward more distant relationships, as expected in large biobank samples. Since S1 follows the simulations by construction, we compare theory with simulation under S1 throughout, and show S3 as a theoretical prediction only.

These are illustrated in Figure 2b. Pair counts for close relatives in the UK Biobank, inferred by KING (Manichaikul et al., 2010; Zhang et al., 2025), fall between our S1 and S3 distributions, suggesting these schemes reasonably bracket real biobank composition.

We then examined *π*_*k*_(*x*) under both sampling schemes to understand how sample composition influences the observed mixture across relatedness. Figure 2d shows *π*_*k*_(*x*) under S1 and S3.

A first key result is that, at low genetic relatedness (*x <* 0.05), both schemes reveal a heterogeneous mixture of many distant pedigree classes. At higher genetic relatedness (*x >* 0.1), relationship classes become increasingly well-separated. Strikingly, in samples dominated by distantly related individuals, mean pedigree relatedness rises sharply between *x* = 0.02 and 0.05, below which the sample consists almost entirely of effectively unrelated pairs (Figure 2c–d).

### 2.3 Genetic architecture and imperfect tagging of causal variants

First, we investigate how imperfectly tagged causal variants and genetic architecture shape the relationship between phenotypic covariance and genetic relatedness within pedigree classes. In the absence of other forces, imperfect tagging attenuates the within-class slope by a factor *υ*_*k*_, and the line passes, to a good approximation, through the origin (Supplementary Section S3.4):

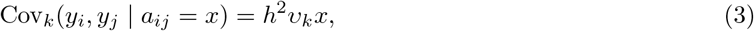

where *h*^2^ is the narrow-sense heritability.

A key quantity is therefore

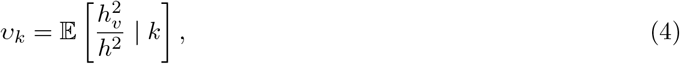

the proportion of heritability captured by observed variants in pairs of pedigree class *k*, where 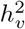is the heritability captured by the observed variants. This quantity determines the slope of phenotypic covariance on genetic relatedness within a class under HE regression, subject to imperfect tagging. Kemper and colleagues previously demonstrated that imperfect tagging can produce patterns similar to empirically observed spectra of covariance across the relatedness spectrum (Kemper et al., 2021), motivating our theoretical characterization of this mechanism.

Let **A** and **G** denote genomic relatedness matrices (GRMs) constructed from the set M of *M* observed marker variants and the set Q of *M*_*q*_ (unobserved) causal variants, respectively:

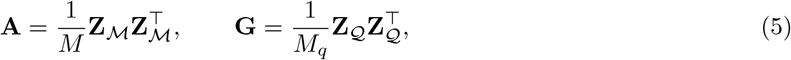

with standardized genotypes 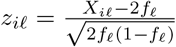, where *f*_*l*_ is the allele frequency of variant *l*; we write *a*_*ij*_ and *g*_*ij*_ for the (*i, j*)-th entries of **A** and **G** respectively. Notice how with this choice of scaling, these matrices are the correlations of the rows (individuals) of the genotype matrix. Under an additive model in which every standardized causal variant has the same effect-size variance, *υ*_*k*_ is the regression slope

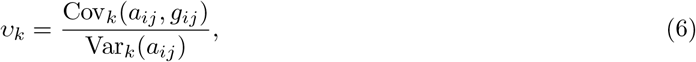

where, as in Eq. (1), the subscript *k* denotes pairs of class *k*.

Among unrelated individuals, *υ*_*∞*_ depends on population-level LD. Yang et al. (2015, Supplementary Note) showed that, among unrelated individuals, this attenuation is the ratio of average LD between markers and causal variants to that among markers; in our notation, 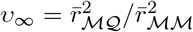(Supplementary Eq. (S10)). When causal variants are rare and markers common, as expected under forces such as stabilizing selection, the maximal achievable LD between these sets of variants is small (Wray, 2005; VanLiere and Rosenberg, 2008), leading to substantial downward bias in heritability estimates.

For related individuals, shorter coalescent times and in turn larger IBD segments induce additional correlations between loci, increasing the extent to which marker variants predict causal variants. The strength of these correlations does not depend on allele frequencies, as the correlations arising from population LD do. As a result, *υ*_*k*_ changes across pedigree relationship classes, with varying contributions from population LD and IBD sharing among close relatives (Figure 3a).

**Figure 3:**
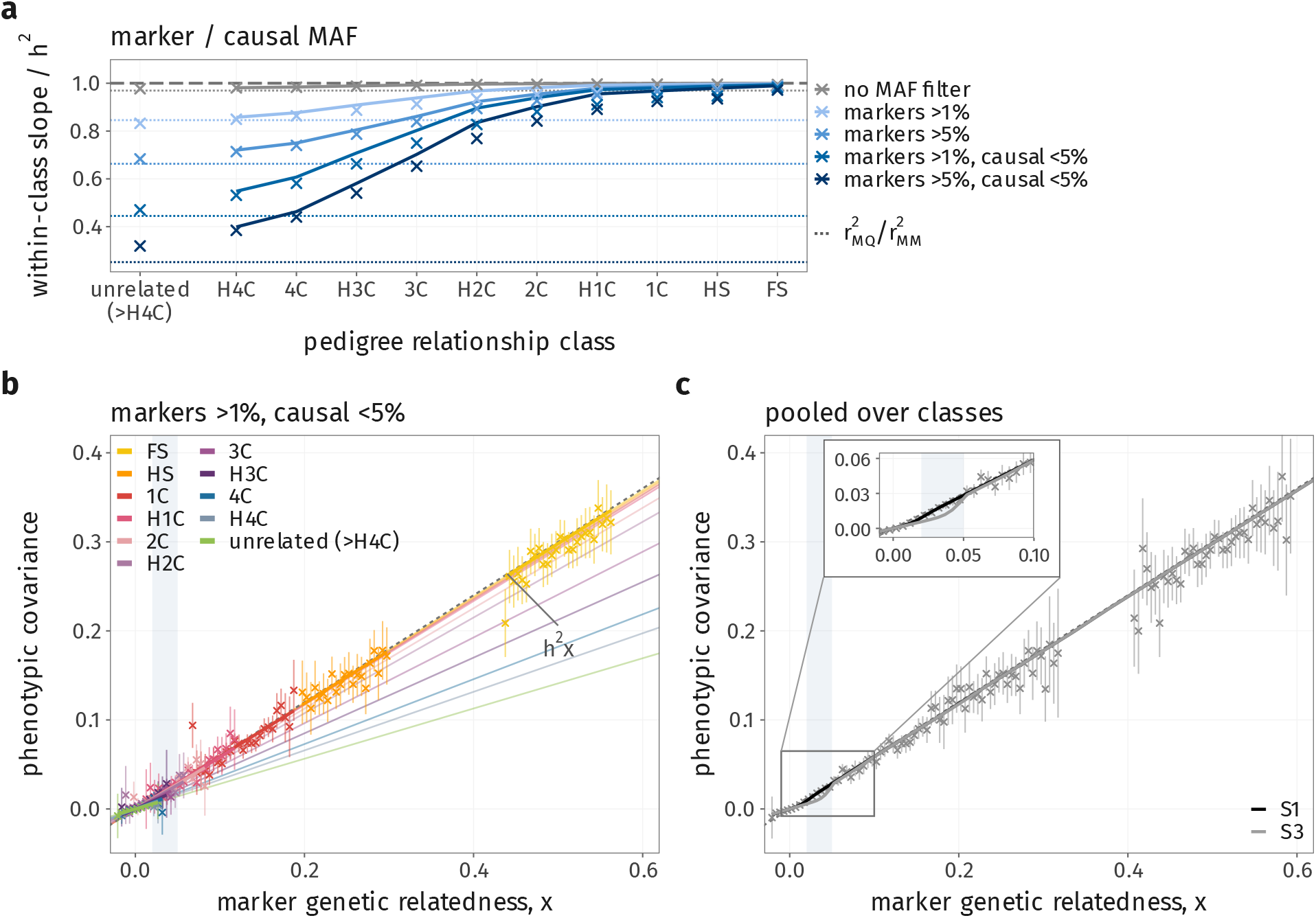
Imperfect tagging of causal variants attenuates the heritability captured in distant relatives. (a) Within-class slope of phenotypic covariance on genetic relatedness, relative to *h*^2^, by pedigree class for five marker and causal-variant MAF scenarios (perfect: every variant is both marker and causal). This ratio is the tagging *υ*_*k*_. Lines, Eq. (8), with *ν*_*k*_ (Eq. (7)) measured on each marker GRM; dotted, its limit among unrelated pairs, 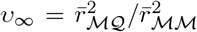; dashed, 1. (b) Mean standardized cross-product 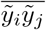 against *x* in each class (markers MAF *>* 1%, causal variants MAF *<* 5%, *h* = 0.6). Lines, Eq. (3); dashed, the additive expectation *h*^2^*x*. (c) As (b), pooled over classes under S1 and S3. Inset, the low-relatedness region. Lines, theory; crosses, simulation under S1 (50 replicates; 95% intervals). Gray band as in Figure 2. Frequency-dependent effect sizes are shown in Supplementary Figure S2. the least-related sampling scheme (S3). This increase matches the empirical patterns observed in Kemper et al., supporting imperfect tagging as a candidate explanation for their findings.

The two sources of tagging can be separated by splitting marker relatedness into two parts (Supplementary Section S2): *x* = *R* + *η*, where *R* is the relatedness implied by the segments the pair shares IBD from the common ancestors that define its pedigree class, and *η* is an independent deviation due to identity by state at the markers outside these segments. We call *η* background LD (BLD): its variance arises from stochastic variation in coalescence across the genome, rather than from segments shared through the recent pedigree. Within class *k*, Var_*k*_(*x*) ≈ Var_*k*_(*R*) + Var(*η*), where 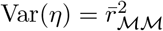 is the same in every class. Let

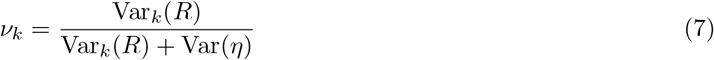

be the proportion of the variance in relatedness within class *k* that is due to IBD. *ν*_*k*_ is close to one in close relatives and goes to zero in unrelated pairs, who by this definition share no segments IBD. An IBD segment carries every causal variant on it, whatever its frequency, so the IBD part tags the causal variants perfectly, while the BLD part tags them only as well as population LD allows. *υ*_*k*_ is therefore the *ν*_*k*_-weighted average of the two (Supplementary Section S3),

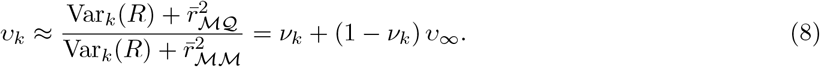

Var_*k*_(*R*), and hence *ν*_*k*_, is difficult to observe empirically, but can be derived analytically as an integral over the genetic map (Supplementary Eq. (S17) and Supplementary Figure S1).

Before we assess this approximation, it is important to note that bias might also arise from misspecified assumptions about genetic architecture. The standard GCTA GRM (Eq. (5)) variance-standardizes all variants and gives them the same weight, which implicitly assumes that every standardized variant has the same effect-size variance, so that rarer alleles have proportionally larger per-allele effects. This is the case *α* = −1 of the more general parametrization of Speed et al. (2017), in which the effect *u*_*l*_ of standardized variant *l* has variance 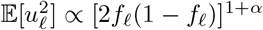. When the true effect-size–frequency relationship differs, *υ*_*k*_ is biased even if all causal variants are observed. The magnitude and direction of this bias can depend on relatedness. We discuss this further in Supplementary Section S3.3 and Supplementary Figure S2.

We then measured *υ*_*k*_ in simulations by constructing GRMs under different genetic architectures and computing the regression slope of the causal-variant GRM on the marker-variant GRM, separately for each pedigree class. The GRM is built from every SNP with minor allele frequency (MAF) of at least 1% in the final generation (about 1,000,000 per replicate — which is close to standard practices).

Figure 3a shows the results: when markers and causals share the same allele frequency spectrum, *υ*_*k*_ ≈ 1 across all classes. However, when we restrict markers to MAF *>* 1% while causals span all frequencies, substantial downward bias emerges. This bias is even more severe when causals are rare (MAF *<* 5%). In this case, about half of the heritability is lost in distant relatives. A very similar pattern emerges when we compute the GRM with the causal SNPs themselves, but simulate a trait with genetic architecture where rare SNPs have larger effect sizes (Supplementary Figure S2).

To predict the phenotypic covariance patterns that imperfect tagging of causal variants would produce, we incorporated the *υ*_*k*_ values obtained from simulations into Eq. (1), varying across sampling schemes. Figure 3b shows the resulting within-class lines for rare causal variants, and Figure 3c their average under S1 and S3. Notably, the predictions show a sharp increase in covariance between *x* = 0.02 and 0.05, where unrelated pairs and distant cousins give way to second cousins — an increase that is most pronounced in

### 2.4 Environmental trait architecture: multi-generational shared environments

Environmental correlations between relatives may also affect the relationship between phenotypic covariance and genetic relatedness. Unlike genetic effects, which depend on realized genome sharing, environmental similarities can persist across arbitrary genealogical distances through mechanisms such as cultural transmission, wealth transfer, and geographic proximity. When present, MGSEs introduce non-genetic phenotypic covariance that varies across pedigree classes, potentially inflating heritability estimates.

We model a phenotype *y*_*i*_ as the sum of additive genetic (*g*_*i*_), shared environmental (*c*_*i*_) and residual environmental (*ϵ*_*i*_) components,

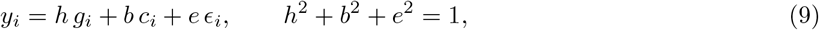

where *h, b* and *e* are scalar coefficients, and *g*_*i*_, *c*_*i*_ and *ϵ*_*i*_ have mean zero and, under random mating, unit variance; the residual *ϵ*_*i*_ is independent of the other components and between individuals. The expected cross-product of standardized phenotypes 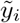 between individuals *i* and *j* (which, under random mating, equals that of the raw phenotypes, since Var(*y*_*i*_) = 1) then decomposes as

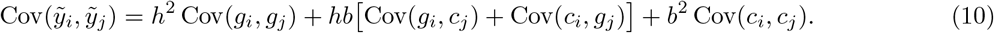

We define Cov_*k*_(*c*_*i*_, *c*_*j*_) = *C*_*k*_, such that *C*_*k*_ captures environmental correlations that extend across arbitrary pedigree classes. These correlations depend on pedigree relationship rather than realized genetic relatedness. We assume that effectively unrelated pairs (class ∞) share no environment, *C*_*∞*_ = 0. This is not restrictive: phenotypes are standardized in the sample, so an environment shared by all pairs shifts every class by the same amount, and *C*_*k*_ can be read as the covariance in excess of that among unrelated pairs. We momentarily assume that Cov(*g*_*i*_, *c*_*j*_) = 0, i.e., that there is no gene–environment covariance; we return to this in Section 2.6, where assortative mating induces one.

The phenotypic covariance is then

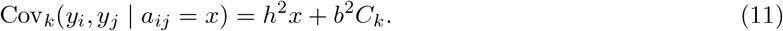

To make these ideas concrete, we adopt a parental-mean transmission model as an example environmental architecture, and mark results specific to it with a superscript PM. Under this model, offspring inherit the mean parental environmental value:

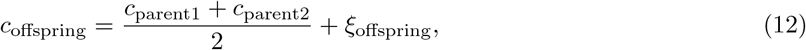

where *ξ*_*i*_ represents independent environmental variation. Let *b*^2^ be the share of phenotypic variance from MGSE under random mating, with Var(*c*) = 1, so that Var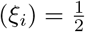. We will index the pedigree relationship class with *k* (*k* = 2 for full siblings, *k* = 3 for half siblings, *k* = 4 for first cousins, and so on), so that the expected relatedness of class *k* is *ā*_*k*_ = 2^−(*k*−1)^. Under parental-mean transmission, the shared environment is passed down the pedigree in the same way as the genome, so its covariance decays geometrically with pedigree distance, at the same rate as expected relatedness:

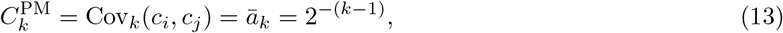

as illustrated in Figure 4a. This equality is specific to the parental-mean model; other forms of environmental sharing, such as a household effect shared only by siblings, give a different *C*_*k*_ (Supplementary Section S4.2). Under this model and random mating, Cov(*g*_*i*_, *c*_*j*_) = 0, so by Eq. (11) the relationship within each class has slope *h*^2^ and intercept 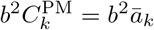.

**Figure 4:**
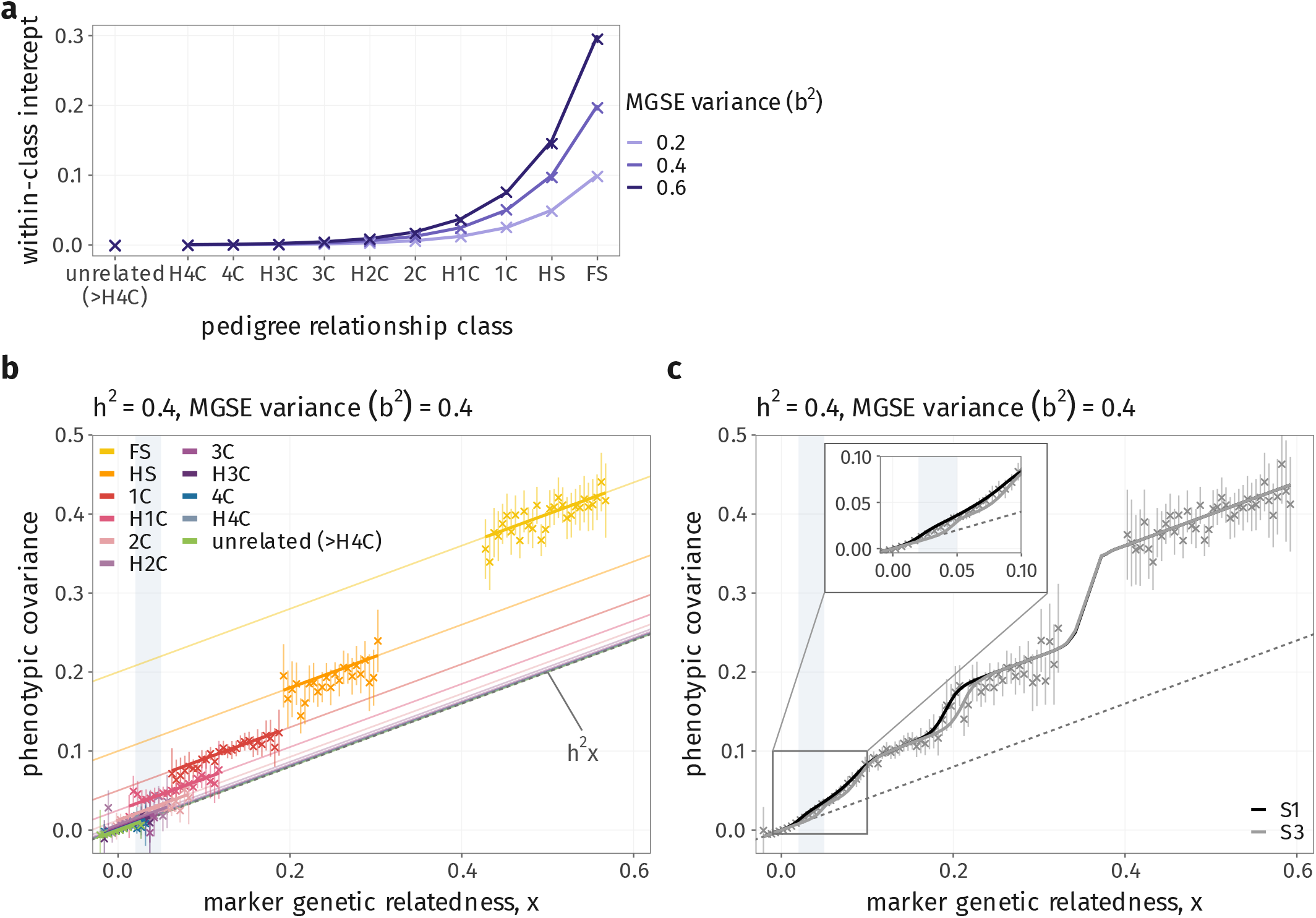
A shared environment transmitted across generations shifts the intercept of each pedigree class; pooled over classes, the shifts become steps. (a) Within-class intercept by pedigree class for *b*^2^ = 0.2, 0.4 and 0.6. Under the parental-mean model it is 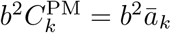, halving with each degree; it is zero among unrelated pairs (*>*H4C), as assumed (Section 2.4). (b) 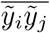 against *x* within each class for *h*^2^ = *b*^2^ = 0.4: parallel lines of slope *h*^2^, offset by *b*^2^*ā*_*k*_; dashed, *h*^2^*x*. (c) As (b), pooled over classes under S1 and S3. Inset, the low-relatedness region. Lines, theory; crosses, simulation under S1 (50 replicates; 95% intervals). Gray band as in Figure 2.

The within-class predictions and the average phenotypic covariance under different sampling schemes are shown in Figure 4b–c. Under this model, we observe sharp increases in covariance in areas where pedigree relatedness increases — such as at the transition between half-siblings and full-siblings, and between *x* = 0.02 and 0.05. Notably, in the low relatedness zone, this pattern is similar to that produced with imperfect tagging. This suggests that in this low relatedness realm, the two forces may not be easily identifiable.

Although we focus here on the parental-mean model, the framework is general: substituting alternative forms of *C*_*k*_ yields corresponding predictions for Cov(*y*_*i*_, *y*_*j*_ | *a*_*ij*_ = *x*) and its local slope (Section 2.7) under different environmental structures.

### 2.5 Assortative mating

When partners are matched on phenotype, assortative mating induces positive correlations between trait-increasing alleles at different, potentially distant, causal loci across the genome. These correlations increase the phenotypic covariance between relatives and inflate the additive genetic variance above its random-mating value (Fisher, 1919). Heritability estimates that assume random mating are therefore often biased. In twin designs, the expected estimate of heritability from an ACE model is 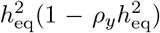, where 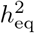is the heritability at equilibrium and *ρ*_*y*_ the phenotypic correlation between mates (Kemper et al., 2021), so assortment biases twin estimates of heritability downwards (Sunde et al., 2024). Regression on IBD sharing between full siblings (Visscher et al., 2006) estimates neither the random-mating nor the equilibrium heritability, but the random-mating genetic variance divided by the equilibrium phenotypic variance (Kemper et al., 2021), and therefore underestimates heritability. Kemper et al. (2021) further detected assortment for height and educational attainment, and inferred mate correlations of 0.24 and 0.60, respectively, by fitting to close relatives (genetic relatedness above 0.05) the expectation that assortment inflates the covariance of relatives by a factor that grows geometrically with the number of meioses separating them (Yengo and Visscher, 2018). Under this expectation the inflation is proportionally largest in distant relatives.

In samples of unrelated individuals, by contrast, assortment can bias heritability estimates upwards. Border et al. (2022) showed that assortative mating biases marker-based estimators: the long-range dependence it induces between causal variants violates the assumption, shared by these estimators, that causal variants are independent apart from local LD. HE regression overestimates heritability relative to both its random-mating and its equilibrium value (whereas GREML is inflated at typical sample sizes but converges to the random-mating value in very large samples). Here we extend these analyses across the full relatedness spectrum, and characterize how assortative mating interacts with MGSE and imperfect tagging. In this section, we will continue to refer to heritability as the random-mating heritability.

We consider a simple form of assortative mating in which partners choose each other on their phenotypes, with correlation *ρ*_*y*_, and analyze the population at equilibrium. In this section there is no shared environment (*b* = 0), so that *y*_*i*_ = *h g*_*i*_ + *e ϵ*_*i*_; in Section 2.6, we study assortative mating together with MGSE.

Assortment inflates the variance of the genetic component above its random-mating value of one (Supplementary Section S5). Mates are matched on their phenotypes, and their genetic values become correlated, with correlation 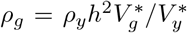, where 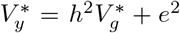 is the phenotypic variance at equilibrium. The genetic variance is inflated to 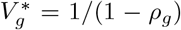 (Fisher, 1919; Crow and Felsenstein, 1968). A closed form for 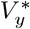 is given in Supplementary Eq. (S31). As in the data, we work with phenotypes standardized at equilibrium, so all covariances below are divided by 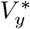.

The expected covariance of relatives follows by tracing paths through the pedigree (Supplementary Section S5). The genetic covariance between relatives of class *k* is *ā*_*k*_ times its within-person value, as under random mating, plus *θ*_*k*_ times its value between mates, where

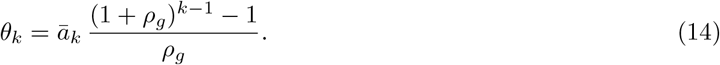

*θ*_*k*_ describes how the resemblance added by each assortative mating accumulates along the path connecting the pair. The class mean is therefore

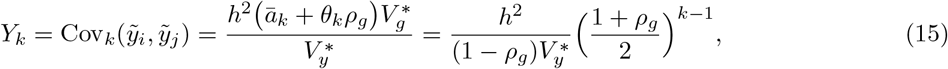

which is consistent with the classical result for the resemblance of relatives under assortment (Fisher, 1919; Nagylaki, 1978; Yengo and Visscher, 2018). Under random mating the class mean is *h*^2^*ā*_*k*_. Assortment multiplies this by 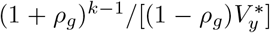, a factor that grows with pedigree distance, so the inflation is proportionally largest in distant relatives (Yengo and Visscher, 2018).

Assortment changes the within-class slope *β*_*k*_ of phenotypic covariance on genetic relatedness *x* by affecting both the phenotypic similarity carried by segments that two individuals share IBD, and that carried by background LD. The BLD deviation *η* covaries with each causal locus together with the long-range dependence that assortment builds up between causal loci (Border et al., 2022), and the slope of phenotypic covariance on *η* is inflated to

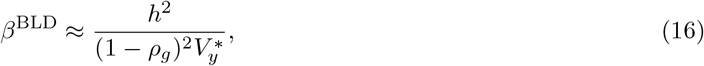

which is the limit that we expect for unrelated individuals. Under positive assortment this is greater than *h*^2^, because 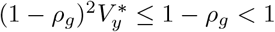 (Supplementary Section S5.6.2).

The slope on the IBD part *R*, in contrast, is

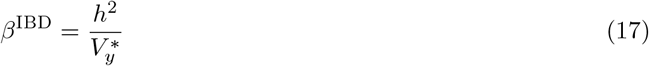

for any strength of assortment (Supplementary Eq. (S50)). Since variation in relatedness among full siblings arises almost entirely from variation in IBD sharing, this is the quantity estimated by IBD regression in siblings, consistent with Kemper et al. (2021). Note that this is lower than *h*^2^.

In more distant relatives, each in-law was chosen on the phenotype of the relative they married, and so covaries with the segments that relative shares IBD with the other side of the pedigree. For more distantly related individuals, this inflation is challenging to derive analytically, and we leave this for further work and use the sibling value *β*^IBD^ in all classes. This understates the within-class slopes of cousins and so overstates their intercepts. At *ρ*_*y*_ = 0.4 (with *h*^2^ = 0.6 and *b*^2^ = 0.2), the simulated slopes of half first to half fourth cousins exceed the prediction by 0.45 *h*^2^ on average (95% interval 0.36–0.54 *h*^2^), about 40% of the predicted slope (Supplementary Figure S7). For first cousins we derive the missing term, which closes most of this gap (Supplementary Section S5.6.1). Even so, the pooled predictions fit the simulations well: the class means are exact, and each within-class line passes through its class mean (Eq. (19)), so a slope error only tilts the line about that point.

The within-class slope moves between these two limits with *ν*_*k*_, the proportion of the variance in genetic relatedness due to IBD (Eq. (7)). It is their *ν*_*k*_-weighted average (Supplementary Eq. (S42)),

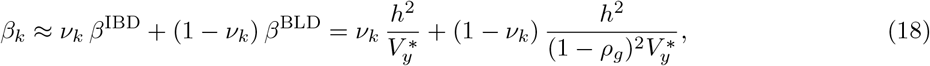

which moves from close to *β*^IBD^ in close relatives (*ν*_*k*_ ≈ 1) to *β*^BLD^ in unrelated pairs (*ν*_*k*_ = 0; Figure 5b). The within-class regression line passes through the class mean *Y*_*k*_ at *x* = *ā*_*k*_, so the phenotypic covariance between relatives of class *k* with observed genetic relatedness *x* is

**Figure 5:**
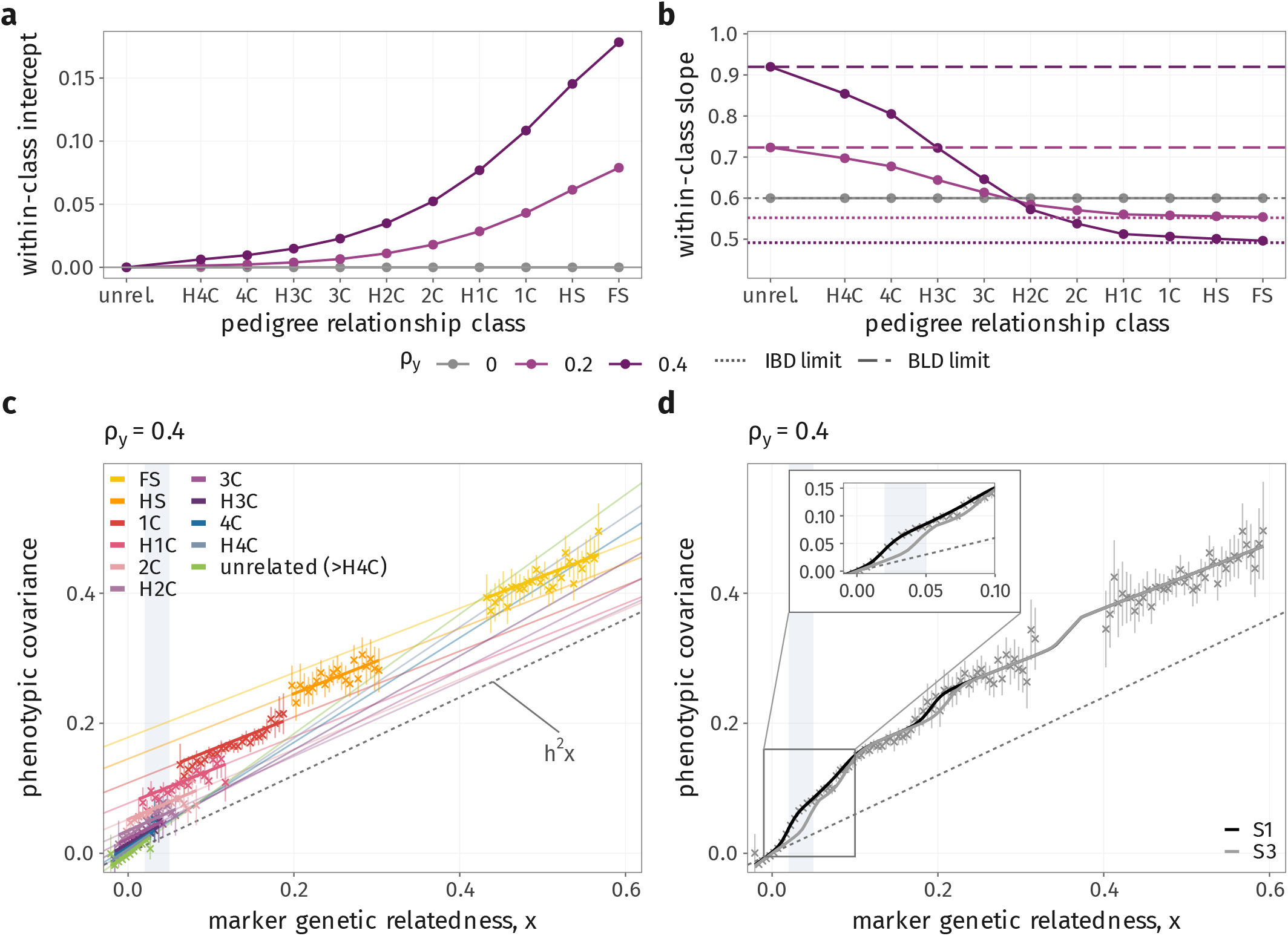
Assortative mating adds class-specific intercepts and moves the within-class slope between an IBD and a BLD limit. No MGSE (*b*^2^ = 0) and *h*^2^ = 0.6. (a) Within-class intercept *α*_*k*_ by pedigree class for *ρ*_*y*_ = 0 (gray), 0.2 and 0.4 (darker for stronger assortment). (b) Within-class slope *β*_*k*_, with the IBD limit 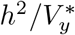 (dotted) and the BLD limit 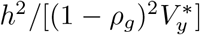 (dashed) for each *ρ*_*y*_. (c) 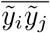against *x* within each class for *ρ*_*y*_ = 0.4; dashed, *h*^2^*x*. (d) As (c), pooled over classes under S1 and S3. Inset, the low-relatedness region. All covariances are divided by the equilibrium phenotypic variance 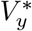. Lines, theory; crosses, simulation under S1 (50 replicates; 95% intervals). Gray band as in Figure 2. For cousins (1C–H4C), the theoretical slopes in (b) and intercepts in (a) are approximate (Supplementary Figure S7).

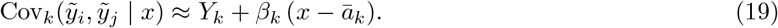

The class mean and the slope together determine the within-class intercept,

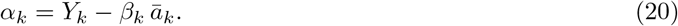

Without assortment, 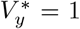, the class mean is *h*^2^*ā*_*k*_ and *β*_*k*_ = *h*^2^, so every intercept is zero and Eq. (19) reduces to the additive expectation *h*^2^*x*. With assortment the intercepts are positive even without a shared environment (Figure 5a). In close relatives, most of the inflation goes into the intercept: assortment packages trait-increasing alleles together, and close relatives inherit these packages whatever their realized IBD sharing. Among unrelated pairs, the inflation comes from small increases in positive LD between trait-increasing variants across the population, and so goes into the slope (Supplementary Section S5.6.5).

Figure 5c–d compares these predictions with simulations for *ρ*_*y*_ = 0.4, within each class and averaged over the pedigree mixture as in Eq. (1). The covariance lies above the additive expectation *h*^2^*x*, and the class-specific intercepts produce steps in the pooled covariance where the class composition changes. The sampling scheme matters mainly between *x* = 0.02 and 0.05 (Figure 5d).

### 2.6 Combining imperfect tagging, MGSE and assortative mating

In the sections above we describe how imperfect tagging of causal variants, MGSEs and assortative mating affect the within-class regression of phenotypic covariance on genetic relatedness in different ways. Imperfect tagging attenuates the slope, most strongly in distant relatives, where the BLD part of relatedness dominates. Under random mating, MGSE shifts only the class intercepts. Assortative mating inflates the class means, most strongly in distant relatives, and moves the slope between an IBD limit below *h*^2^ and a BLD limit above it.

#### 2.6.1 MGSE under assortative mating

Combined with MGSE, assortment affects the inheritance patterns in the model in two further ways (Supplementary Section S5). First, the shared-environment component is inflated too, to variance 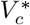. Second, *g* and *c*, independent under random mating, acquire a covariance *ω*^∗^ = Cov(*g*_*i*_, *c*_*i*_). This is because people with high genetic values tend to mate with partners with high shared-environment values, and their offspring inherit both.

The results in this section are specific to the parental-mean model described above, although several of their features hold for any form of MGSE. The genetic and shared-environment components of mates are correlated, with correlations *ρ*_*g*_ and *ρ*_*c*_ respectively. Many of the results from the case without MGSE carry over if we replace 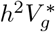 by 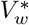 and *ρ*_*g*_ by *ρ*_*w*_, where *w* = *hg* + *bc* is the whole transmissible component of the phenotype, genetic and shared-environmental, with mate correlation 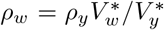. The phenotypic variance at equilibrium is then 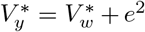. Summing the genetic, gene–environment and shared-environment components gives the class mean

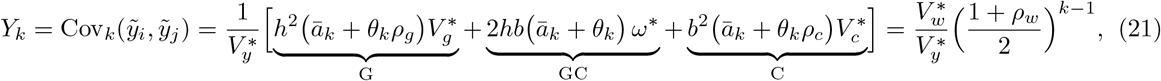

where *θ*_*k*_ is as in Eq. (14) with *ρ*_*w*_ in place of *ρ*_*g*_ (Supplementary Eq. (S38)). With *b* = 0, 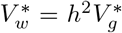 and *ρ*_*w*_ = *ρ*_*g*_, and Eq. (21) reduces to Eq. (15).

The IBD limit is unchanged, 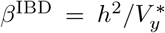, because neither the shared environment nor the gene– environment term varies with IBD sharing. The BLD limit becomes

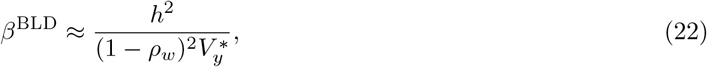

which is inflated further than without MGSE, since *ρ*_*w*_ *> ρ*_*g*_ when *b >* 0, and reduces to Eq. (16) when *b* = 0. The within-class slope and intercept keep the form of Eqs. (18) and (20),

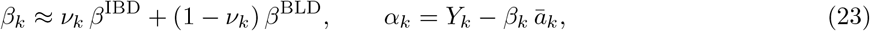

with *Y*_*k*_ now the class mean of Eq. (21).

An interesting consequence of the gene–environment coupling is that the shared-environment and gene–environment components now acquire BLD slopes, 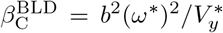 and 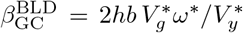 both of which are zero under random mating (Supplementary Table S3). Through *ω*^∗^, the shared environment covaries with the genotypes, so it carries part of the slope on BLD. The genetic and gene–environment components, in turn, contribute to the intercepts. Assortment and MGSE therefore act together to distort the relationship more than either alone. By contrast, with *h*^2^ = 0, *ω*^∗^ = 0: assortment raises the class intercepts but leaves the lines flat.

#### 2.6.2 The full model

Imperfect tagging acts only on the BLD part of the slope. The IBD part does not depend on which markers are used, while in the BLD part tagging scales the slope by *υ*_*∞*_, as without assortment. Within the BLD part the two mechanisms are separable: assortment scales the covariance of every causal locus with the markers by the same factor, whatever its frequency (Supplementary Section S6). With all three mechanisms, the within-class slope is therefore

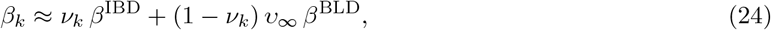

with *β*^IBD^ and *β*^BLD^ as in Section 2.6 and *υ*_*∞*_ as in Eq. (8). Without assortment, Eq. (24) reduces to *h*^2^*υ*_*k*_ (Eq. (8)). Among unrelated pairs the slope is *υ*_*∞*_*β*^BLD^, the product of two separate factors: tagging multiplies the slope by *υ*_*∞*_, which is below one for rare causal variants, and assortment multiplies it by 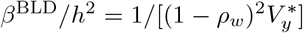, whatever the tagging. In proportion, assortment therefore biases the slope by the same amount whether or not the causal variants are well tagged. This agrees with Border et al. (2022), who found that the bias of marker-based heritability estimators under assortative mating was qualitatively similar when a large fraction of variants was not genotyped.

Among relatives, tagging removes a fraction (1 − *ν*_*k*_)(1 − *υ*_*∞*_) *β*^BLD^*/*[*ν*_*k*_*β*^IBD^ + (1 − *ν*_*k*_)*β*^BLD^] of the slope. This fraction grows with the ratio *β*^BLD^*/β*^IBD^, which assortment raises from one to 1*/*(1 − *ρ*_*w*_)^2^. Imperfect tagging therefore removes a larger fraction of the slope under assortative mating than under random mating (Supplementary Eqs. (S62)–(S63)), not because the two mechanisms interact within the BLD part, but because assortment shifts the slope towards the part that tagging attenuates.

Tagging does not change the intercepts, which stay at their values when the markers are the causal loci, M = Q (Eq. (23)). Each within-class line therefore pivots about its intercept, and the class mean falls to

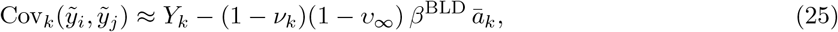

with *Y*_*k*_ as in Eq. (21). This reduction is small in every class: in close relatives *ν*_*k*_ ≈ 1, and in distant relatives *ā*_*k*_ is small. Figure 6 shows these effects within each class: MGSE raises the class covariances and intercepts, while tagging lowers the BLD limit of the slope and leaves the intercepts unchanged.

**Figure 6:**
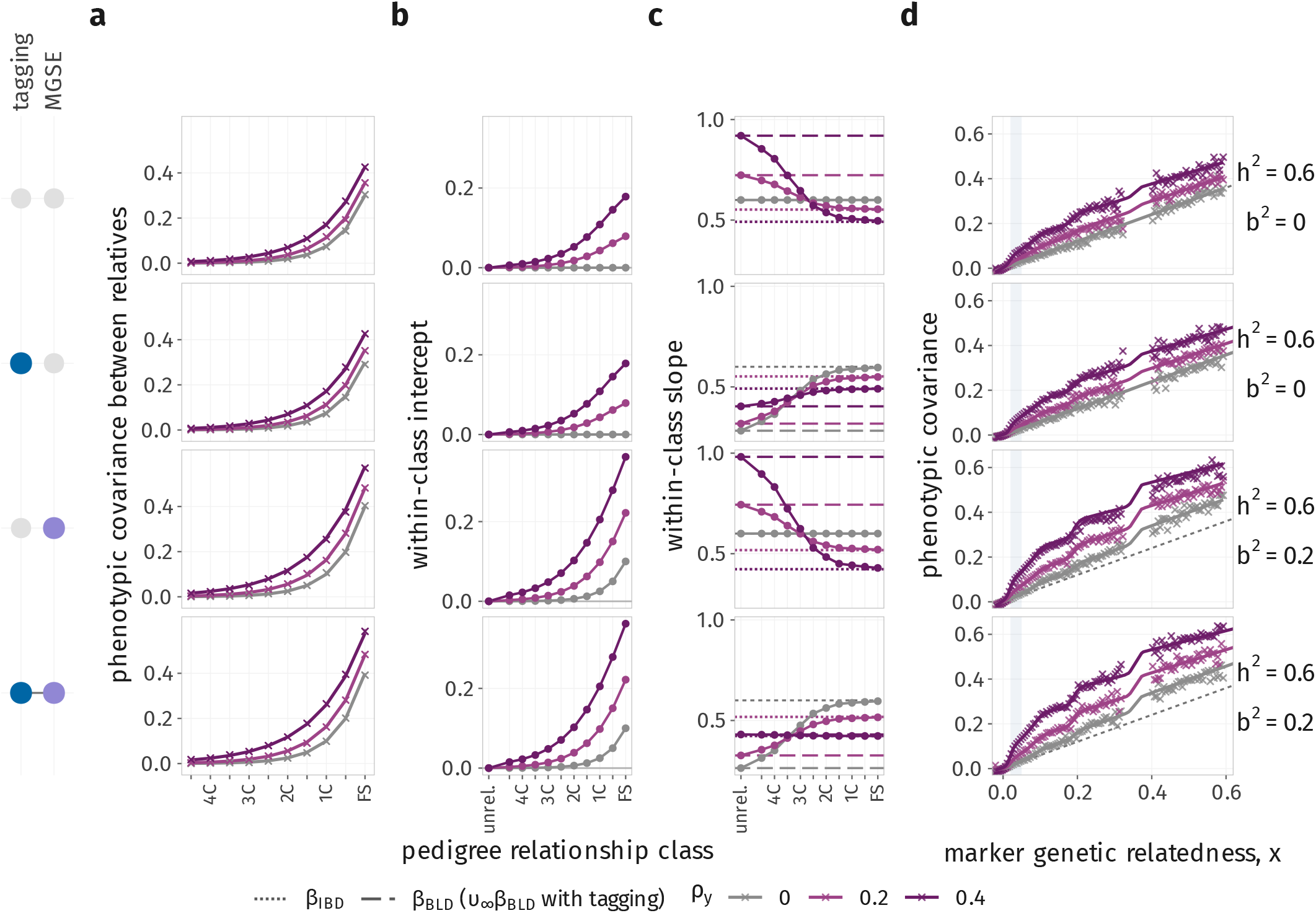
Under assortative mating, MGSE raises the class covariances and intercepts, while imperfect tagging lowers the BLD limit of the slope. Each row is one combination of mechanisms, shown by the dot matrix (left): assortment alone, with tagging, with MGSE, and with both. *h*^2^ = 0.6; tagging means markers with MAF *>* 1% and causal variants with MAF *<* 5%; MGSE means *b*^2^ = 0.2 with parental-mean transmission; *ρ*_*y*_ = 0 (gray), 0.2 and 0.4 (darker for stronger assortment). (a) Class covariance 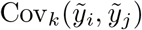 by pedigree class. (b) Within-class intercept *α*_*k*_. (c) Within-class slope *β*_*k*_, with the IBD limit *β*^IBD^ (dotted, *ρ*_*y*_ *>* 0) and the BLD limit *β*^BLD^ (dashed) for each *ρ*_*y*_; with tagging, the BLD limit is *υ*_*∞*_*β*^BLD^, drawn also for *ρ*_*y*_ = 0, where it is *υ*_*∞*_*h*^2^. (d) 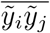 against *x*, pooled over classes under S1; dashed, *h*^2^*x*. All covariances are divided by 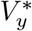 Lines, theory; crosses, simulation (50 replicates). Gray band as in Figure 2. For cousins (1C–H4C) under assortment, the theoretical slopes in (c) and intercepts in (b) are approximate (Supplementary Figure S7).

Figure 7a shows the pooled covariance for each mechanism alone and in combination. Assortment amplifies the steps that MGSE produces in the pooled covariance.

**Figure 7:**
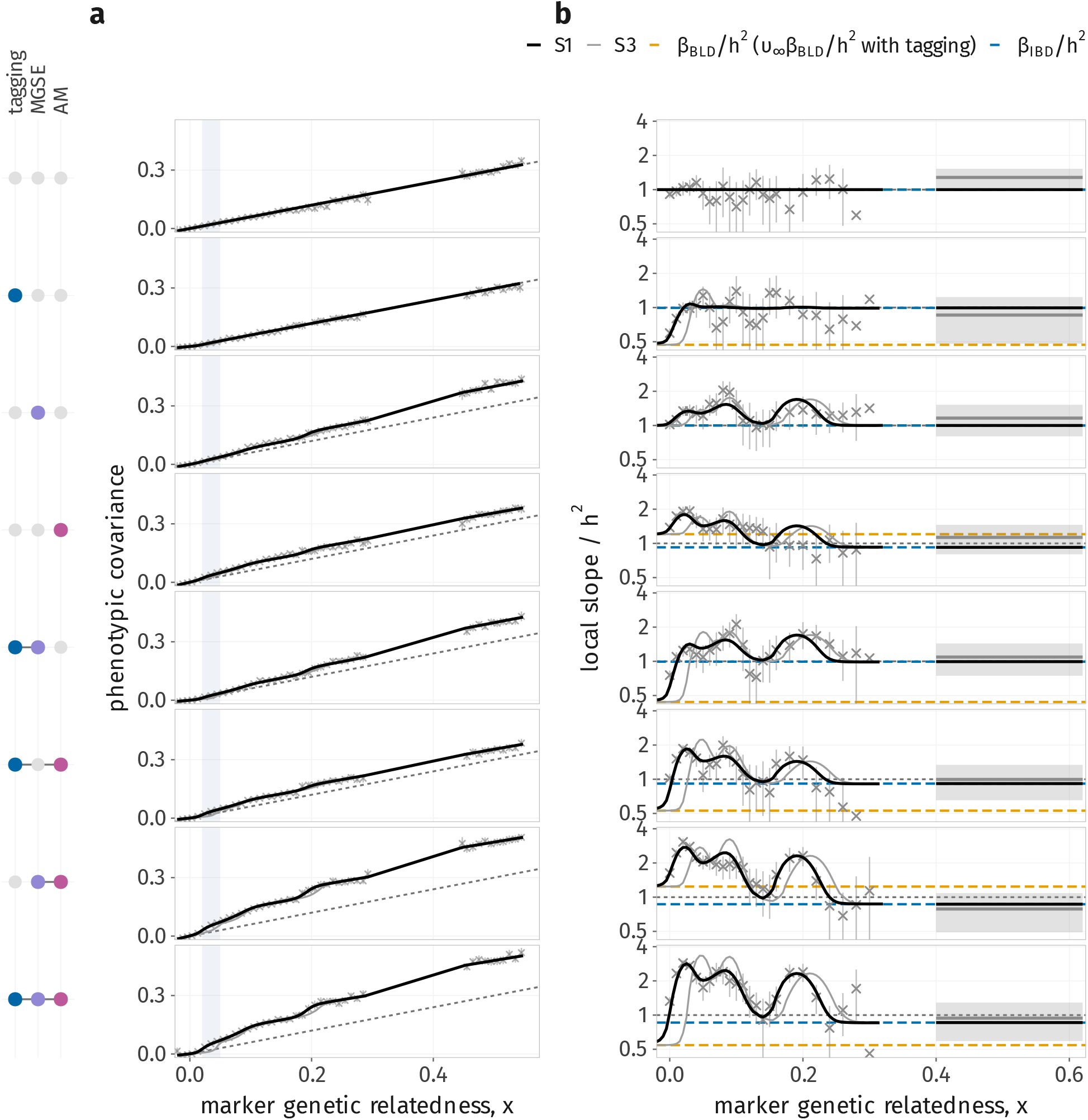
Imperfect tagging, MGSE and assortative mating leave distinct signatures in the local slope of phenotypic covariance on genetic relatedness. Each row is one combination of mechanisms, shown by the dot matrix (left). *h*^2^ = 0.6; tagging means markers with MAF *>* 1% and causal variants with MAF *<* 5%; MGSE means *b*^2^ = 0.2 with parental-mean transmission; AM means *ρ*_*y*_ = 0.2. (a) 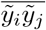 against *x* (divided by 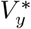 under AM); dashed, *h*^2^*x*. (b) Local slope of the pooled covariance relative to *h*^2^, log scale: a least-squares line through the bin means within ±0.02 of *x*, each bin counted once (the window widens to ±0.04 over the sparse half-sibling range), and a single line over 0.40 ≤ *x <* 0.62, almost all full siblings. Dashed lines, the BLD limit among unrelated pairs, *β*^BLD^*/h*^2^ (orange; *υ*_*∞*_*β*^BLD^*/h*^2^ with tagging), and the IBD limit within full siblings, *β*^IBD^*/h*^2^ (blue); dotted, *h*^2^. Lines, theory (S1 black, S3 gray); crosses, and gray line and band for full siblings, simulation under S1 (50 replicates; 95% intervals). Gray band as in Figure 2.

### 2.7 Local slopes bridge different heritability designs, but are not bounded by them

A quantity of interest is the local slope of the relationship between phenotypic covariance and genetic relatedness. For example, among unrelated pairs, where all of the variation in relatedness is due to BLD, it is the expectation of the Haseman–Elston estimate of SNP heritability. Instead, among full siblings it is that of sibling IBD regression. We would like to know how it moves between these two limits, and where it departs most from the underlying heritability. In real data, the true heritability in the population is unknown, so the level of the local slope is not informative on its own; how the local slope changes across relatedness, however, may be.

Regressing phenotypic covariance on genetic relatedness within a narrow window of *x* estimates the derivative of the pooled covariance, which by Eq. (1), with within-class lines *α*_*k*_ + *β*_*k*_*x*, is

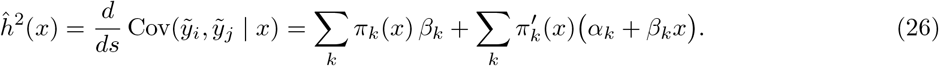

The first sum averages the within-class slopes, weighted by the class composition at *x*. The second is non-zero only where that composition changes with *x*, and there it adds the differences between the class lines: a mechanism that shifts the lines of different classes by different amounts moves the local slope away from *h*^2^ even when every within-class slope equals *h*^2^. Where one class dominates, the local slope is that class’s slope.

With imperfect tagging only, every line passes through the origin (Eq. (3)), so

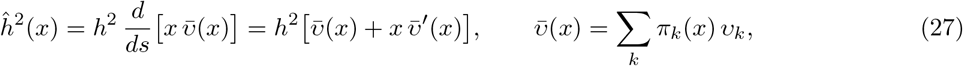

which rises from *h*^2^*υ*_*∞*_ among unrelated pairs to *h*^2^*υ*_2_ ≈ *h*^2^ among full siblings. With MGSE only (Eq. (11)),

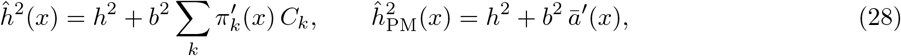

where *ā* (*x*) = ∑_*k*_ *π*_*k*_(*x*)*ā*_*k*_ is the mean pedigree relatedness of Figure 2c: the local slope exceeds *h*^2^ wherever *ā* (*x*) rises faster than *x*. With assortative mating only, Eq. (26) applies with *β*_*k*_ and *α*_*k*_ from Eqs. (18) and (20). Its two limits, 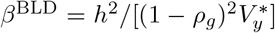 among unrelated pairs and about 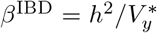 among full siblings, lie on either side of *h*^2^. When all three are present, it applies with *β*_*k*_ from Eq. (24) and the intercepts of Eq. (23), which tagging leaves unchanged. Among unrelated pairs, the limit is then 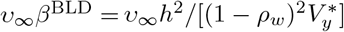.

Figure 7b shows how the local slope moves between these two limits. Rather than interpolating smoothly between them, it leaves the range spanned by these two slopes when the pedigree composition shifts, reaching about three times *h*^2^ near *x ≈* 0.02 under S1. This follows from Eq. (26), as the second term, which captures the shifting mix of relatives, can add to the slope purely because the mixture is shifting between classes of pedigree relatives.

The three mechanisms leave distinct signatures in this local slope. Imperfect tagging lowers it below *h*^2^ among unrelated pairs, and it recovers at intermediate relatedness, where IBD accounts for a growing share of the variance in relatedness (a higher *ν*_*k*_). MGSE leaves the slope at *h*^2^ among unrelated pairs and raises it wherever the class composition changes. Assortative mating raises it slightly above *h*^2^ among unrelated pairs and further between *x* = 0.02 and 0.05.

### 2.8 Empirical investigation

Here we show the results of a preliminary investigation in the UK Biobank (UKB, Bycroft et al. (2018)). Our aim was to examine the relationship between phenotypic covariance and genetic relatedness across the full range of relationships available in the UKB. We used fine bins across the whole spectrum (Figure 8) to distinguish mechanism-specific signatures, predicted in Figure 7, that coarse binning obscures.

**Figure 8:**
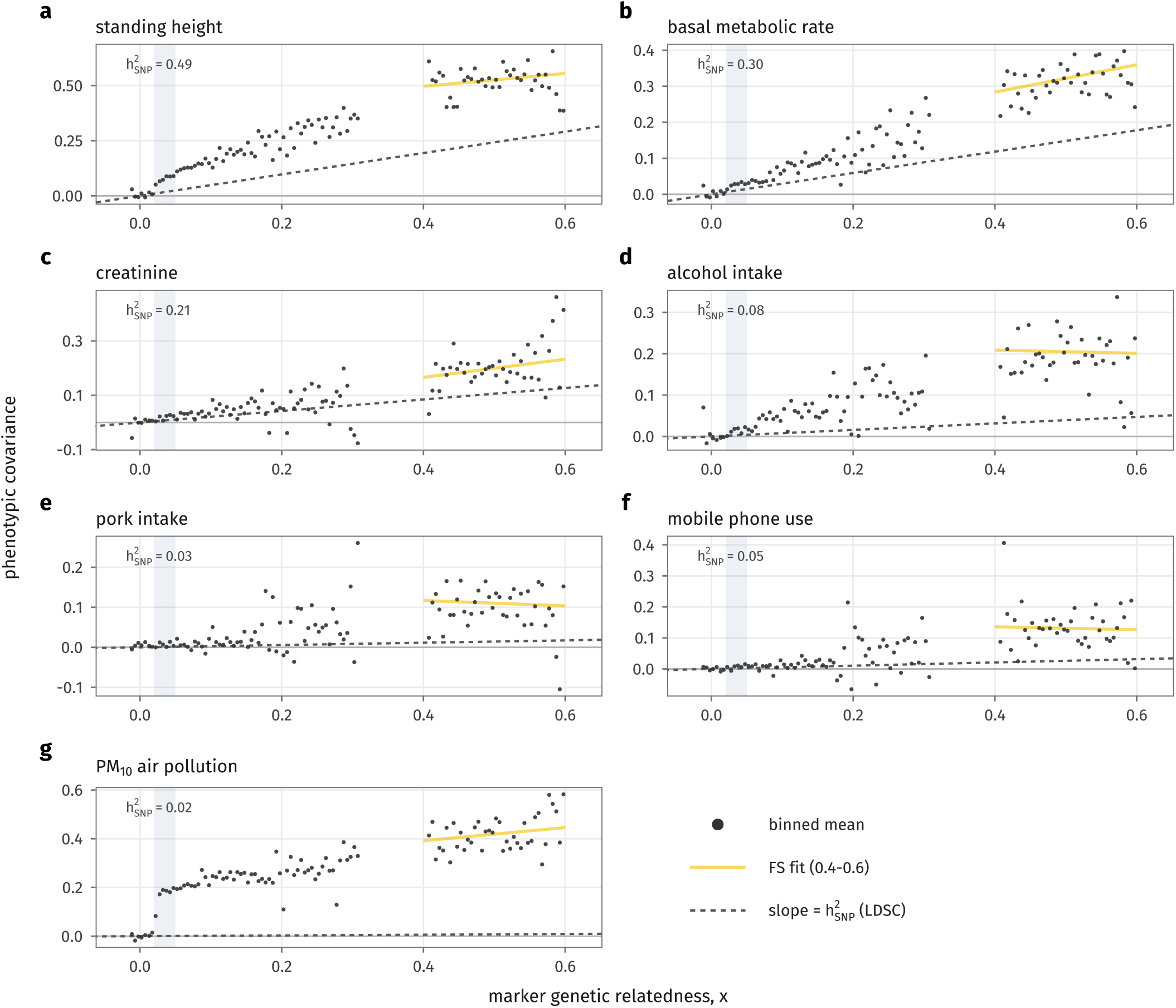
Phenotypic covariance across genetic relatedness for seven UK Biobank traits. Phenotypic covariance, estimated as the mean cross-product of standardized residuals, 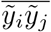 in bins of genetic relatedness *x* (dots; width 0.005, bins with more than 50 pairs), for anthropometric and metabolic traits (a–c) and lifestyle and environmental traits (d–g). Gold line, least-squares regression with an intercept within the full-sibling range (0.4 ≤ *x* ≤ 0.6), fitted to individual pairs. Dashed line, slope equal to the SNP heritability estimated by LD score regression in unrelated individuals, 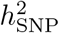 (Neale Lab; value in each panel). Shaded band, 0.02 ≤ *x* ≤ 0.05. Residuals adjusted for sex, age, year of birth and genotyping batch.

Phenotypic covariance between pairs of individuals was estimated as the cross-product of standardized residuals after regressing each trait on sex, age at recruitment, year of birth and genotyping batch, following Kemper et al. (2021). We focused on approximately 430,000 CEU/GBR-like individuals, selected by similarity to 1000 Genomes individuals. Relatedness was estimated from HapMap3 SNPs with frequency above 1%. For computational efficiency, we did not compute relatedness for all pairs: we screened for pairs with relatedness above 0.03 on a GRM built from 50,000 SNPs, added 100,000 random pairs to represent the unrelated end of the spectrum, and recomputed relatedness for all of these pairs on the full SNP set. Covariance is shown in bins of width 0.005. We also show an ordinary least-squares regression of the pair-level cross-products on relatedness within the full-sibling range (≤0.4 *x ≤*0.6) (Supplementary Section S9).

As a complication, we faced several limitations to exhaustively exploring the UK Biobank results due to recent curtailments in access. In particular, we only show results that did not include genomic principal components, so fine-scale population structure may contribute to the covariances, and we show no uncertainty intervals on phenotypic covariances.Finally, the reference slope 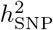 we use for visual baseline comes from the Neale Lab’s *h*^2^ LD score regression on rank-inverse-normal transformed traits (on the UKB ‘White British’ samples).

We also investigated these patterns in the All of Us (AoU) cohort (The All of Us Research Program Genomics Investigators, 2024). Since the ancestry filtering left around 228,000 individuals, the power to resolve finer bins was substantially lower than in the UK, so we focus on the preliminary UKB results here and report the All of Us results in Supplementary Section S11. The All of Us analysis uses all pairs, genomic principal components, parent–offspring separation and a delete-block jackknife over individuals (Supplementary Section S11).

Our investigation reveals foremost that there is wide variation in the covariance spectra across traits (Figure 8; all other traits in Supplementary Section S10). Most phenotypes’ phenotypic covariance rises above that predicted by the SNP heritability.Anthropometric and metabolic traits (height, basal metabolic rate and creatinine; Figure 8a–c) show steadily increasing covariance. Lifestyle, dietary, and environmental exposure traits (Figure 8d–g) are more consistent with shifts in intercept near pedigree-class transitions, which, though not definitive, resemble effects expected in the presence of multi-generational shared environments (Figure 4).

A striking example is exposure to particulate matter air pollution (Figure 8g, PM10). Its covariance is close to zero among pairs with *x <* 0.02 but rises to about 0.19 by *x* ≈ 0.03–0.05, and the covariance between full siblings is around 0.4, whereas its SNP heritability 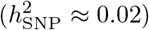 would imply a covariance of only about 0.01. One explanation is an MGSE that decays more irregularly as one extends deeper into the pedigree than the simplified parental-mean model considered in the theoretical modeling above; for example, MGSE arising from multi-generational patterns of shared residency in locales that vary by pollution status. Further assortment between mates on location or environment could exacerbate this pattern. Adjusting for Local Authority of birth had little effect on these patterns (data not shown), suggesting that the shared component is not explained by broad birthplace, orreflects finer-scale patterns not captured by the Local Authority variable.

As another focal example, we consider height, which is estimated to have a phenotypic correlation between mates of 0.27 (Horwitz et al., 2023). For height, the empirical pattern shows an intercept shift near *x* 0.3– 0.4, roughly the half-sibling to full-sibling transition, which resembles our predictions under assortative mating of these magnitudes (Figure 5), although, as noted above, such steps are not specific to assortment. The slope within the full-sibling range is harder to read in the UK Biobank, because there it also contains parent–offspring pairs, which we did not separate (Supplementary Section S9) and whose covariance and genome sharing behave differently from those of full siblings under assortment. In All of Us, where parent– offspring pairs were identified with KING and separated from the full-siblings, the full-sibling slope for height is 0.69 (Supplementary Figure S17), close to the full-sibling IBD estimate of 0.66 reported by Kemper et al.(2021) and to its expectation under assortment, 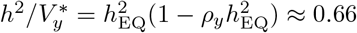. Presumably since height is measured precisely and in a large sample, it also shows a remarkably clear pattern, with a clear concave shape from *x ≈* 0.02 upwards similar to that predicted under assortative mating alone (Figure 6d, top row, with *h*^2^ = 0.6 and *ρ*_*y*_ up to 0.4).

## 3 Discussion

In summary, we use a framework that conditions on realized genome sharing to model phenotypic covariance across relatedness, under three processes known to affect complex traits: imperfect tagging, MGSE and AM. Realized genome sharing is a proxy for covariance at the causal variants, and in our analytical model it can be separated from effects that depend on pedigree relatedness. This does not fully separate the mechanisms behind the resemblance of relatives, but it brings us closer.

A key finding is that, in a biobank, the mix of pedigree classes at a given genetic relatedness changes across the relatedness spectrum. The shift between *x* = 0.02 and 0.05 is particularly important: below it, pairs are effectively unrelated; above it, detectable relatives begin to dominate, especially in samples made up mostly of distant relatives, such as the UK Biobank. All three mechanisms produce non-linearity through this shifting mixture, and all are expected to produce a sharp rise in covariance in this range. Coarse bins of genetic relatedness average over it and so obscure the signatures of the different mechanisms; a finer view across the whole range is needed to see them.

Our pedigree classes are a simplification of real data: we consider only pairs in the same generation who descend from full or half siblings, and so do not separate relationships of the same degree, such as half siblings, avuncular pairs and grandparent–grandchild pairs. These differ in the variance of their realized relatedness (Hill and Weir, 2011), and plausibly in how much environment they share. However, the findings on the shifting mixture depend only on how the within-class distributions overlap and on the composition of the sample. They should therefore hold more generally.

Our theory and simulations show that when the markers used to build the GRM are filtered by allele frequency, as is common practice, and causal variants are rare, much of the heritability can be missed in distant relatives, consistent with previous results (Yang et al., 2010; Kemper et al., 2021), and our contribution is to show how this effect accrues as we move from close relatives, where tagging occurs via IBD segments from recent shared ancestors, to distant relatives, where it is mostly due to BLD. We also show that this situation can be understood as a misspecification of genetic architecture, and even if all causal variants are observed, assuming a wrong relationship between allele frequency and effect size can bias the slope of phenotypic covariance on the GRM. Whole-genome sequence data should largely remove the bias from missing rare variants, consistent with the results of Wainschtein et al. (2026). A misspecified relationship between allele frequency and effect size, however, leaves a bias even when every causal variant is genotyped, because it comes from the model of genetic architecture rather than from missing variants.

We also model MGSEs, shared environments that persist across generations. MGSEs depend only on the pedigree, not on realized genetic relatedness, so they shift the within-class intercepts. Pooled across classes, these shifts appear as rises in covariance wherever the mixture moves toward pairs more closely related through the pedigree. We show results for a parental-mean model, but these properties hold whatever the form of the environmental covariance *C*_*k*_, as long as it depends only on pedigree class (Supplementary Section S4.2).

It is interesting to consider the implications of these patterns for heritability estimation. Many approaches seek to model family shared environments explicitly, or to absorb them into additional variance components in order to avoid biasing heritability estimates. Some assign separate environmental components to particular relationships, such as nuclear families, couples and siblings (Xia et al., 2016), or model shared familial environment within a structural equation model (Muñoz et al., 2016). Others model close and distant relatives using different relatedness matrices (Zaitlen et al., 2013), or use only the variation in relatedness that remains after conditioning on parental genotypes (Young et al., 2018). Each of these makes assumptions about which relatives share an environment and how that sharing decays with pedigree distance. Our findings suggest that an ideal approach would control for pedigree relatedness directly, allowing each class its own intercept, without specifying how the environment decays. This would extend the logic of IBD regression within full sibships (Visscher et al., 2006), which uses only the variation in realized relatedness among siblings, to every pedigree class. Under random mating, the within-class slopes would then be free of MGSE. However, pedigree class can typically be recovered from genotype data only up to about the third degree, so this is likely to be challenging in most datasets. Even so, allowing for different intercepts across regions of relatedness, and examining how the slope changes across them, may reveal key features of a trait’s architecture and which regions are likely to lead to biases.

On its own, AM changes the within-class slopes and intercepts, and so biases slope-based estimates of heritability. Combined with the other mechanisms, it amplifies and couples them. The bias it induces grows with both *h*^2^ and *b*^2^, so it matters most for highly heritable traits with a strong shared environment. In close relatives, AM mostly raises the intercept: it bundles trait-increasing alleles together, and close relatives inherit these bundles irrespective of their realized IBD sharing. In consequence, AM lowers the slope in close relatives, because it inflates the phenotypic variance but not the IBD part of the slope. This leaves the sibling IBD slope at 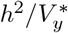, below *h*^2^, agreeing with (Kemper et al., 2021, Supplementary Note 3). In contrast, AM raises the slope in distant relatives, toward the unrelated limit 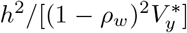, above *h*^2^, consistently with Border et al. (2022).

A thread running through all three mechanisms is the split of genetic relatedness into an IBD part and a BLD part. The two carry different information: IBD tags every causal variant on a shared segment, whatever its frequency, whereas BLD tags causal variants only as well as population LD allows and, under assortment, also carries the long-range dependence between causal loci. The share of the variance in relatedness that is due to IBD therefore sets how each mechanism acts within a class, and its decline across the relatedness spectrum moves the within-class slopes between the sibling and the unrelated limits. However, averaging over the shifting mixture of relatives can push the slope above the bounds.

Finer binning of the UK Biobank relatedness spectrum reveals heterogeneity across traits that coarser analyses obscure. Kemper et al. (2021) summarized pairs with relatedness above 0.05 in seven bins, whereas we use bins of width 0.005 throughout; to our knowledge, this is the first analysis to resolve the covariance of relatives this finely above 0.05. On a preliminary basis, we find coarsely that anthropometric traits tend to show a more consistent, linear relationship over moderate to high relatedness than dietary, lifestyle and exposure traits, and the most striking deviations from linear patterns are in exposure to pollution, and dietary traits, where apparent shifts in intercept resemble the MGSE prediction (although such shifts are not specific to MGSE alone).

Finer bins resolve more structure, but without pedigree information the mechanisms are not fully separable. In population data, one direction would be to estimate *π*_*k*_(*x*) and then fit alternative models of trait architecture. Datasets with recorded pedigrees would allow more direct tests: for example, contrasting pairs with the same genetic relatedness but different pedigree classes would isolate the environmental intercept, and so distinguish MGSE from imperfect tagging. Relatedness disequilibrium regression (Young et al., 2018) offers another route: by conditioning on parental genotypes, it uses only the variation in relatedness due to Mendelian segregation, which an environment shared through the pedigree cannot mimic, so it is robust to MGSE of any form. Its limitation is that it requires genotyped parents. One could also fit the model to real data. Under assumptions about *π*_*k*_(*x*), and with *ν*_*k*_, *υ*_*k*_ and the closed forms for *β*_*k*_, *α*_*k*_ and *Y*_*k*_, estimating *h*^2^, *b*^2^, *ρ*_*y*_ and *υ*_*∞*_ for each trait is tractable, and would allow the mechanisms to be compared formally. Until then, the readings of individual traits in our empirical analysis are descriptive. We leave this for future work.

It will be important to extend these analyses beyond the UK Biobank, to other cohorts and ancestries, and to datasets that span a broader range of environments. There is no reason to expect environmental exposures and assortative mating patterns to be shared in the same way across datasets, so contrasting covariance profiles across cohorts may be informative from an epidemiological perspective. Tagging effects may also differ substantially between ancestries, since they depend on LD and on the allele frequency spectrum. One might also speculate that, if a trait is affected by MGSEs, conditioning on known environmental factors affecting it, where possible, would reduce staircase-like increases in the phenotypic covariance profiles.

These patterns may also matter for phenotypic prediction. If phenotypic covariance is not linear in genetic relatedness, predictors that assume it is, such as GREML-based prediction or polygenic scores trained in a sample with a given mixture of relatives, may perform differently depending on how closely the target individuals are related to the training sample. Modeling a non-linear relationship between genetic relatedness and phenotypic covariance directly might recover some of this information, for example in a BLUP-style estimator, without requiring a detailed model for the shared environment or assortment.

In conclusion, we study a central but poorly understood relationship in quantitative genetics: the increase in phenotypic covariance across the genetic relatedness spectrum. As this relationship is shaped by every process acting on a trait in a population, including assortative mating patterns, and by the composition of the cohort in which it is measured, it is a rich summary of trait architecture, which we hope may help to elucidate these features. Observing it across the full range of relatedness is a new opportunity afforded by large-scale biobank data. The empirical patterns we observe suggest that there is much to learn from these covariance profiles, and from how they vary across traits, cohorts and environments.

## Acknowledgments

We thank Carl Veller and Jeremy Berg for helpful discussion. This work was supported by the National Institutes of Health (grant R21HG014303 to JN). We also thank the participants of All of Us and the UK Biobank, which was accessed under application number 88057.

## Code and data availability

Code for the simulations, the theoretical predictions and every figure in this paper is available at https://github.com/mkiravn/glitch-paper-code.

## S1 Phenotypic model

We model a phenotype with a genetic and an environmental component. We use standardized path coefficients so that we can more easily use path analysis:

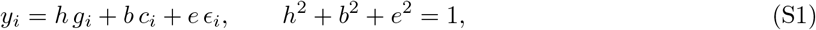

where *g*_*i*_ is the breeding value, *c*_*i*_ a familial environmental component, and *ϵ*_*i*_ independent noise, each with unit variance under random mating. We write *w* := *h g* + *b c* for the transmissible part of the phenotype, so that *y* = *w* + *e ϵ*. Unstarred symbols denote values under random mating and starred symbols equilibrium values under assortment: *V*_*g*_ and 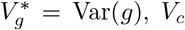 and 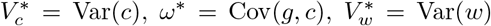and 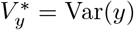 We set *V*_*g*_ = *V*_*c*_ = 1, so that *h*^2^, *b*^2^ and *e*^2^ are shares of the phenotypic variance under random mating, and keep *V*_*g*_ explicit only where it distinguishes the genic from the equilibrium variance (§S5). As in the data, we work with phenotypes standardized at equilibrium, 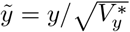; under random mating 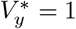and 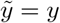.

The genetic architecture consists of *M*_*q*_ causal loci, each with effect size *u*_*l*_ on the standardized genotype, such that

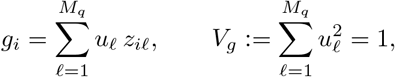

where *z*_*il*_ is the standardized genotype and *V*_*g*_ the genic variance.

We focus on a specific form of environmental transmission which we call “parental-mean”, where the offspring inherit the midparent environment plus noise:

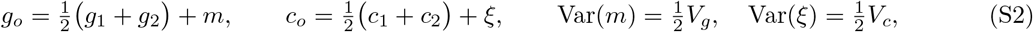

with the Mendelian term *m* and its environmental analog *ξ* independent of each other and of the parents (Rice et al., 1978; Cloninger et al., 1979; Cavalli-Sforza and Feldman, 1981).

We also have relationship classes, which denote pairs by their relationship through the pedigree. We use an index for these, *k*, that starts at *k* = 2 for first-degree relatives, and encodes the expected relatedness in the absence of inbreeding for a pair of class *k, ∑*_*k*_ = 2^−(*k*−1)^, so that *k* = 2 for full sibs and parent–offspring, *k* = 3 for half sibs, avuncular and grandparent–grandchild pairs, and *k* = 4 for first cousins — although the classes we analyse are all pairs in the same generation who descend from full or half siblings (collateral relatives). Under random mating lineal and collateral pairs of the same class behave alike; under assortment they do not (§S5), and the results for assortative mating below are for collateral pairs. A subscript *k* on a moment, as in Cov_*k*_(·,·) and Var_*k*_(·), denotes the moment taken across pairs of class *k*, and we write 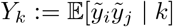 and *C*_*k*_ := E[*c*_*i*_*c*_*j*_ | *k*].

We will be interested in the regression of the phenotypic cross-product on genomic relatedness *x* per class *k* (Haseman and Elston, 1972; Elston et al., 2000; Kemper et al., 2021), which is

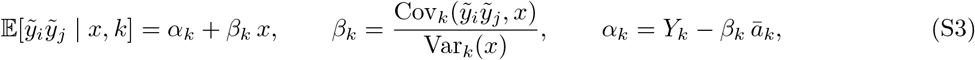

using E[*x k*] = *∑*_*k*_. Since the regression passes through the class means, *α*_*k*_ + *β*_*k*_*x* = *Y*_*k*_ + *β*_*k*_(*x* − *ā*_*k*_). Pairs of different classes overlap in *x*, so the expected cross-product at a given *x* averages these lines over the classes present, with weights *π*_*k*_(*x*) = Pr(*k* | *x*) (Eq. (1) of the main text), and its local slope is Eq. (26). For each mechanism below we derive *β*_*k*_, *α*_*k*_ and *Y*_*k*_, from which both follow.

## S2 Decomposing relatedness into IBD and background LD

Many of the results will depend on relatedness and on whether it is due to identity by descent (IBD) or to identity by state outside IBD segments, which we call background LD (BLD). The variance of the BLD part arises from stochastic variation in coalescence across the genome, rather than from segments shared through the recent pedigree. Throughout, M is the set of *M* markers used to compute relatedness and Q the set of *M*_*q*_ causal loci; in each simulated scenario both are chosen by allele frequency. For each pair of individuals, we can calculate genomic relatedness as the cross-product of standardized genotypes over the markers:

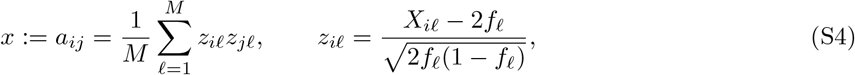

where *X*_*il*_ ∈ {0, 1, 2} is the diploid genotype value of individual *i* at locus *l* and *f*_*l*_ its allele frequency. The standard genomic relatedness matrix (GRM) is the matrix **A** composed of the entries *a*_*ij*_ (Yang et al., 2010).

Suppose we knew the segregation at meiosis at every locus (call it *σ*, which indicates which allele was transmitted). We can decompose this relatedness into a part that depends on *σ*, and a part that does not:

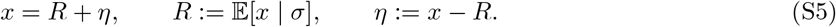

*R* is the genomic relatedness implied by the transmissions, without knowing the allele frequencies or genotypes at each locus (cf. Powell et al., 2010). Writing *S*_*l*_ ∈ {0, 1, 2} for the number of alleles the pair shares IBD at locus *l*, E[*z*_*il*_*z*_*jl*_ | *σ*] = *S*_*l*_*/*2, so

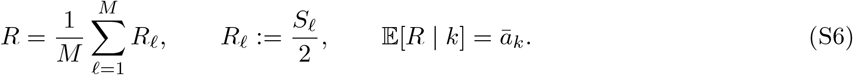

*η* is the residual part, which depends on the allele values. Its variance depends on the effective number of loci in the marker panel, which we will call *M*_*e*_. For a pair sharing nothing IBD, the genotypes of *i* and *j* are independent, so each covariance below factorizes exactly into 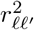, with 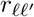; the correlation between the standardized genotypes at *l* and *l*^*′*^:

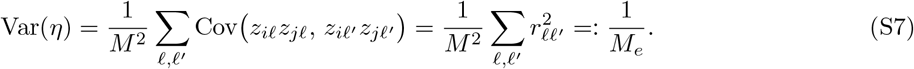

This quantity is in turn related to the average linkage disequilibrium (LD) between the loci 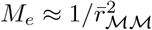, with 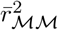 the average *r*^2^ over pairs of markers) (Goddard et al., 2011; Goddard, 2009). Since E[*η*] = E[E [*η* | *σ*]] = 0 and E[*Rη*] = E[*R* E[*η* | *σ*]] = 0, we have Cov(*R, η*) = 0.

The variance of *x*, and its covariance with the phenotypic cross-product, therefore split into IBD and BLD parts. We use this decomposition to model the regression of the cross-product on relatedness under both imperfect tagging (§S3) and assortative mating (§S5).

Conditioning on the relationship degree *k*: The IBD part clearly depends on *k*, since *k* determines the distribution of *σ*. However, *η* has mean zero given *σ* whatever the class, and to a good approximation its variance is the unrelated value (S7) for every *k*:

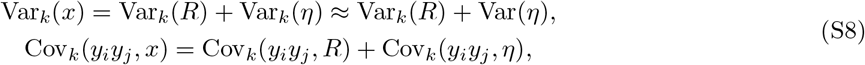

where the equalities are exact, because *x* = *R* + *η* and Cov_*k*_(*R, η*) = 0 in every class. Only Var_*k*_(*η*) ≈ Var(*η*) is an approximation.

Another quantity of interest will be the proportion of variance in relatedness that is due to IBD, rather tha BLD, for relatives of degree *k*:

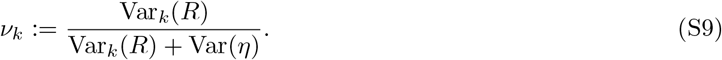

There is greater variance in IBD proportion shared between close relatives, compared to distant ones. This variance was studied by Hill and Weir (2011); Veller et al. (2020), and depends on relationship degree and the recombination map. At the extreme of complete unrelatedness, the variance is driven completely by BLD, and is relatively small.

## S3 Imperfect tagging of causal variants

### S3.1 Heritability from marker-based relatedness

In practice causal SNPs are unobserved, so the true causal covariance **G** is approximated by the genetic covariance at the observed markers, **A**. The resulting slope of the genetic–phenotypic covariance relationship is

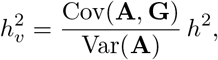

the heritability captured by the observed variants, so the Haseman–Elston (HE) estimate is biased downward by the factor Cov(**A, G**)*/* Var(**A**). Yang et al. (2015, Supplementary Note, Eqs. 6–7) showed that among unrelated individuals this factor depends on population-level LD:

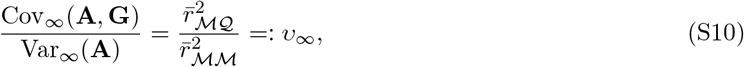

where 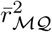 is the average LD *r*^2^ between marker and causal variants and 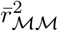 that among marker variants. Here we use ∞ to subscript “unrelateds”. Wray (2005) derived that the maximum possible LD between two variants with minor allele frequencies 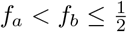 is 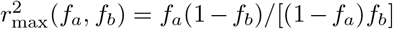, and VanLiere and Rosenberg (2008) showed this to be the global maximum over haplotype configurations. Markers at higher frequency than causal variants therefore give 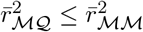, and hence bias heritability downwards.

Equation (S10) holds among unrelated pairs. The question we approach here is how this slope varies across relatedness. Empirically, phenotypic covariance rises faster than marker-based relatedness among distant relatives, which Kemper et al. (2021) attributed to imperfect tagging of causal variants by the markers.

### S3.2 Two-locus decomposition

Conditioning (S10) on the pedigree class gives *υ*_*k*_. Writing *Z*_*l*_ := *z*_*il*_*z*_*jl*_ for the same-locus cross-product of a pair, each matrix entry is an average of these, 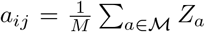 and 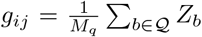, so the matrix ratio becomes a ratio of double sums over locus pairs:

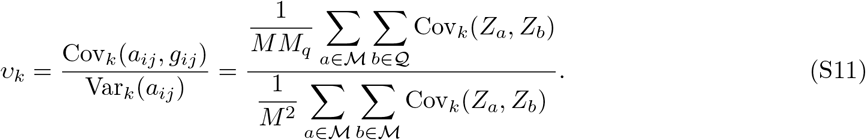

The numerator and denominator differ only in which locus pairs are averaged over — M × Q above and M × M below.

With *S*_*l*_ the number of alleles shared IBD at locus *l* (S6), write *S* = (*S*_*a*_, *S*_*b*_) for the joint state at the two loci. *σ* determines *S*_*l*_ at every locus, and E[*Z*_*l*_ | *σ*] = *S*_*l*_*/*2 (S6). The IBD part of the relatedness in (S5) is *R* (S6), the average IBD state over the markers. By the law of total covariance,

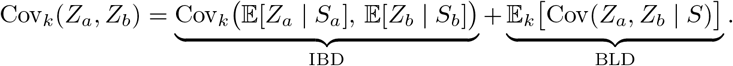

Substituting E[*Z*_*l*_ | *S*_*l*_] = *S*_*l*_*/*2, the first term is 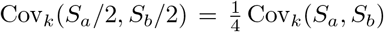. For classes with no IBD2 states — everything except full sibs in an outbred pedigree — we can write *ϕ*_*k*_ for the one-locus IBD probability and *τ*_*k*_(*c*_*ab*_) for the correlation of the two IBD indicators at recombination fraction *c*_*ab*_, with *τ*_*k*_(0) = 1 and 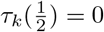 (Hill and Weir, 2011),

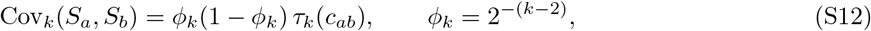

since with no IBD2 state *S*_*l*_ is a Bernoulli indicator of probability *ϕ*_*k*_, whose variance is *ϕ*_*k*_(1−*ϕ*_*k*_), and *τ*_*k*_ is by definition the correlation between the indicators at the two loci. Together these give 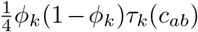 for the IBD term. The second term depends on the allele frequencies and on the LD between the two loci, and is obtained state by state. Write 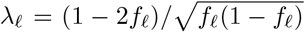 for the third moment of a standardized Bernoulli allele indicator *x*, which satisfies *x*^2^ = 1 + *λ*_*l*_*x*. Alleles on the same founder haplotype carry the population LD, and alleles on different founder haplotypes are independent. Conditioning on the joint state, and on which founder haplotype each allele comes from, then gives Table S1 exactly for biallelic loci, without any normal approximation. The IBD state can differ between the loci only if a crossover between them has broken the shared haplotype in one carrier. That carrier’s allele at the other locus then comes from another founder haplotype and is independent of the shared allele.

**Table S1:** Two-locus covariances conditional on the joint IBD state, assuming no consanguinity, so that *S*_*l*_ ∈ {0, 1, 2} counts cross-individual IBD allele pairs, for tightly linked loci. Where *S*_*a*_ ≠ *S*_*b*_, one carrier’s shared haplotype has recombined between the loci. (2, 0) and (0, 2) need two crossovers and take 0 or 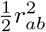 depending on which haplotypes recombined. Every entry is a combination of 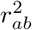and *r*_*ab*_*λ*_*a*_*λ*_*b*_.

| $S_a$ | $S_b$ | $\mathbb{E}[Z_a Z_b \mid S_a, S_b]$ | $\text{Cov}(Z_a, Z_b \mid S_a, S_b)$ |
| --- | --- | --- | --- |
| 0 | 0 | $r_{ab}^2$ | $r_{ab}^2$ |
| 1 | 0 | $\frac{1}{2}r_{ab}^2$ | $\frac{1}{2}r_{ab}^2$ |
| 0 | 1 | $\frac{1}{2}r_{ab}^2$ | $\frac{1}{2}r_{ab}^2$ |
| 2 | 0 | 0 or $\frac{1}{2}r_{ab}^2$ | 0 or $\frac{1}{2}r_{ab}^2$ |
| 0 | 2 | 0 or $\frac{1}{2}r_{ab}^2$ | 0 or $\frac{1}{2}r_{ab}^2$ |
| 1 | 1 | $\frac{1}{4} + \frac{3}{4}r_{ab}^2 + \frac{1}{4}r_{ab}\lambda_a\lambda_b$ | $\frac{3}{4}r_{ab}^2 + \frac{1}{4}r_{ab}\lambda_a\lambda_b$ |
| 2 | 1 | $\frac{1}{2} + \frac{1}{2}r_{ab}^2 + \frac{1}{4}r_{ab}\lambda_a\lambda_b$ | $\frac{1}{2}r_{ab}^2 + \frac{1}{4}r_{ab}\lambda_a\lambda_b$ |
| 1 | 2 | $\frac{1}{2} + \frac{1}{2}r_{ab}^2 + \frac{1}{4}r_{ab}\lambda_a\lambda_b$ | $\frac{1}{2}r_{ab}^2 + \frac{1}{4}r_{ab}\lambda_a\lambda_b$ |
| 2 | 2 | $1 + r_{ab}^2 + \frac{1}{2}r_{ab}\lambda_a\lambda_b$ | $r_{ab}^2 + \frac{1}{2}r_{ab}\lambda_a\lambda_b$ |

Every entry of the table is a combination of 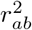 and *r*_*ab*_*λ*_*a*_*λ*_*b*_, so averaging each column over Pr(*S*_*a*_, *S*_*b*_ | *k*) produces exactly two coefficients; call them *γ*_*k*_ and *ψ*_*k*_. The *r*_*ab*_*λ*_*a*_*λ*_*b*_ term appears only in the rows with *S*_*a*_ ≥ 1 *and S*_*b*_ ≥ 1, so *ψ*_*k*_ comes entirely from pairs sharing IBD at both loci: it vanishes among unrelated pairs and grows as relatives become closer. The (0, 0) row is the unrelated case and returns 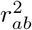 exactly, which is where Var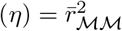 comes from. Collecting,

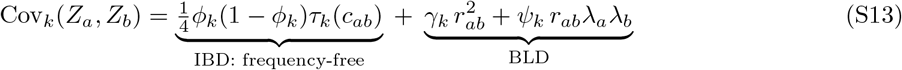

with *γ*_*k*_ and *ψ*_*k*_ the state-averaged coefficients just described. Both depend on *k* and, through the probabilities of the joint states, on the recombination fraction between the loci, but not on the allele frequencies. The frequencies enter the BLD term through *λ*_*a*_*λ*_*b*_, and through *r*_*ab*_ itself, whose attainable range they bound (§S3.1).

The two pieces of (S13) are the two parts of (S5). Averaging over ordered pairs of markers gives Var_*k*_(*x*), and the terms separate as

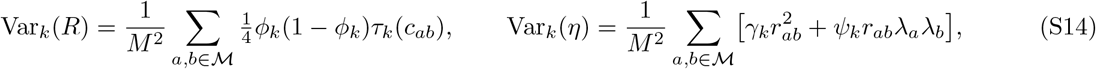

reproducing Var_*k*_(*x*) = Var_*k*_(*R*) + Var_*k*_(*η*) of (S8). The frequency-free term is the IBD part; the frequency-dependent term is the BLD part.

A useful approximation for estimating these quantities genome-wide is *γ*_*k*_ = 1, *ψ*_*k*_ = 0, which gives

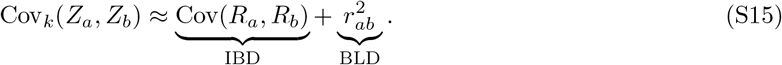

The additional terms are dropped because they contribute little to the averages but are dominated by rare variants, through *λ*_*a*_*λ*_*b*_, which grows without bound as *f* → 0, so their locus-pair estimates are extremely heavy-tailed and destabilize the ratio. Setting *γ*_*k*_ = 1 is justified by Table S1: the (0, 0) row has coefficient exactly 1, and Pr(*S*_*a*_ = *S*_*b*_ = 0 | *k*) → 1 as the IBD term becomes small, so *γ*_*k*_ → 1 in precisely the regime where the BLD term is what matters. Under (S15), Var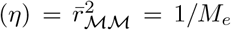 with 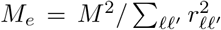the effective number of independent markers (Goddard et al., 2011; Goddard, 2009). If, in addition, the causal loci are spread over the genetic map like the markers, so that the IBD covariance between marker and causal loci equals that among markers (Cov_*k*_(*R, R*_*Q*_) = Var_*k*_(*R*); §S7.3), the class-*k* attenuation becomes a weighted average,

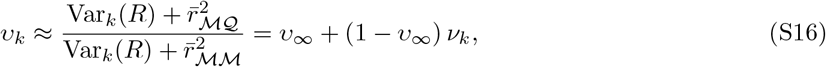

with *ν*_*k*_ the IBD share of the variance in relatedness (S9). Within families the IBD term dominates and *υ*_*k*_ → 1: identity by descent is a property of the segment, not of the markers used to detect it, which is why family designs are less sensitive to marker ascertainment than population studies. Among unrelated pairs *υ*_*k*_ → *υ*_*∞*_, which is consistent with (S10). The average over pairs of markers in (S14) is an integral over the genetic map. With markers at density *µ*(*t*) at map position *t* (in Morgans), normalized so that ∑_chr_∫*µ*(*t*) *dt* = 1,

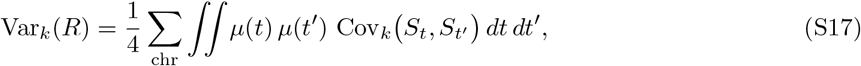

where 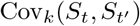 depends only on the recombination fraction 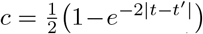 (Haldane’s map function; Haldane, 1919): *it is ϕ* (1 − *ϕ*_*k*_)*τ*_*k*_(*c*) for classes without IBD2 states (S12) and 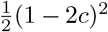 for full sibs. Loci on different chromosomes are unlinked and contribute nothing. With markers spread uniformly over the map, *µ* is constant and (S17) gives the values of Hill and Weir (2011); we evaluate it at the simulated marker positions, which are evenly spaced in base pairs. Figure S1 compares (S17) with the simulations, together with the IBD share *ν*_*k*_ it implies.

**Figure S1:**
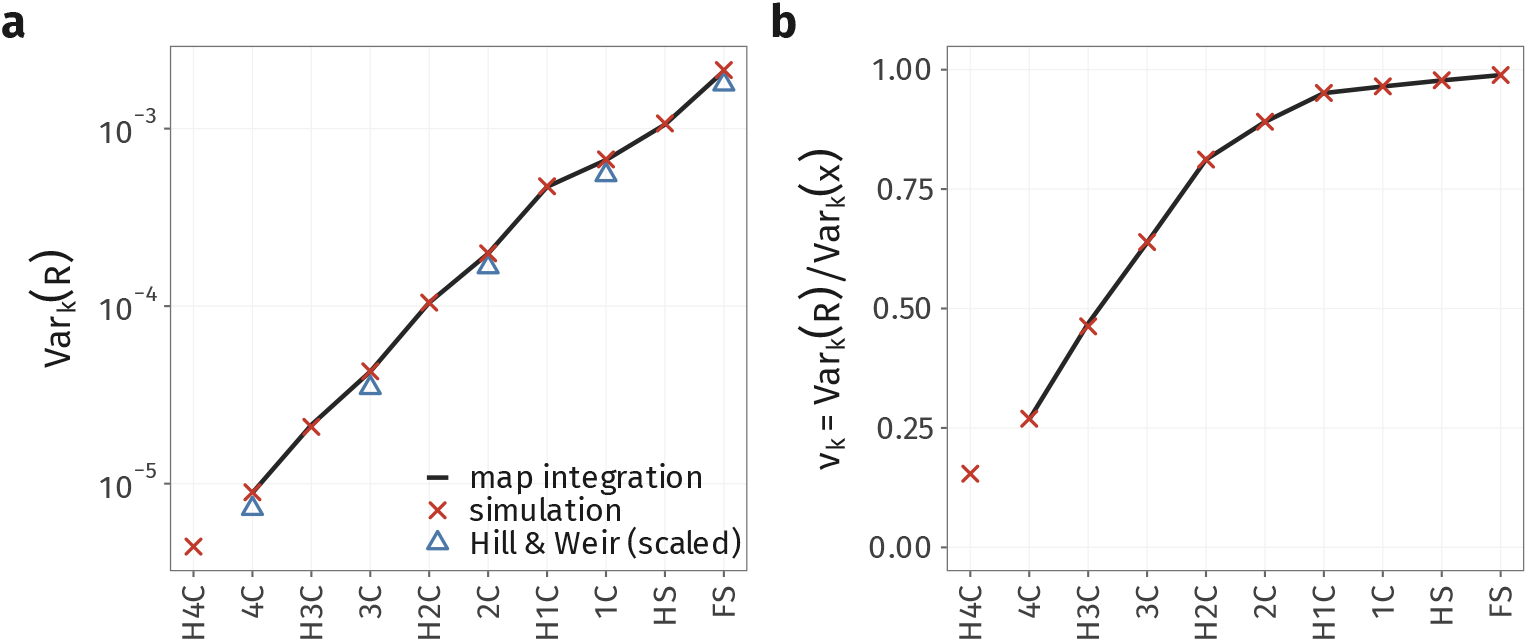
The variance of IBD relatedness within each pedigree class, from the genetic map and from simulation. (a) Var_*k*_(*R*) by pedigree class: the map integral (S17) over the simulated marker positions with the Haldane map function (line, available to *k* = 10, where the two-locus IBD correlations stop); simulation, 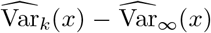 (crosses, 50 replicates; 95% intervals, ±1.96 jackknife SE); and the values tabulated by Hill and Weir (2011) (triangles). Hill and Weir assume 22 autosomes in five length classes (2 × 2.75, 4 × 2.10, 6 ×1.75, 8 × 1.25 and 2 × 0.75 M; 35.9 M in total), Haldane’s map function, and loci spread uniformly along the genetic map; we scale their values to the simulated genome (29.1 M) by the ratio of uniform-map integrals for the two genomes (1.17–1.20). The remaining difference, 16–19%, arises because the simulated markers are evenly spaced in base pairs rather than on the genetic map. (b) The IBD share *ν*_*k*_ = Var_*k*_(*R*)*/* Var_*k*_(*x*), from simulation 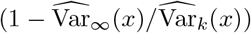 and from the map integral, with Var(*η*) taken from unrelated pairs.

### S3.3 Frequency-dependent effect sizes

The derivation so far treats all causal loci symmetrically, with fixed effects. Under a random-effects model in which the effect-size variance depends on allele frequency, there may be imperfect tagging effects even when every causal variant is genotyped.

Let 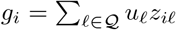 with E[*u*_*l*_] = 0 and per-locus effect-size variance 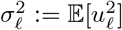, allowed to depend on allele frequency. (This is a variance indexed by a locus, and is not to be confused with the segregation *σ*, which has no index.) Averaging over the effects for given genotypes,

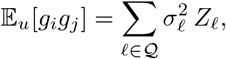

so the object the trait actually defines is not the **G** of (S11) but a *weighted* causal matrix

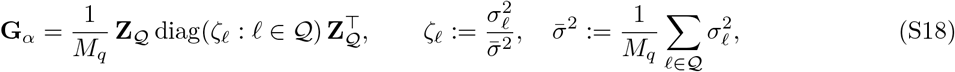

where *ζ*_*l*_ is the effect-size variance of locus *l* relative to the mean over causal loci, so that **G**_*α*_ = **G** when every locus has the same variance. The class-specific attenuation is then (S11) with **G**_*α*_ in place of **G**,

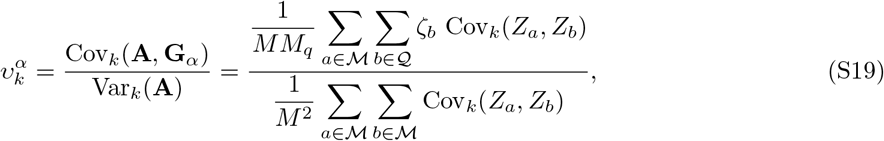

which is (S11) with each marker–causal pair reweighted by the relative contribution *ζ*_*b*_ of its causal locus to the additive genetic variance. Everything else — the two-locus decomposition (S13), the split into IBD and BLD parts — goes through unchanged, because the weights touch only which pairs are averaged, not the covariance being averaged.

A common specification is the *α*-model (Speed et al., 2012, 2017), which on the standardized scale used here is

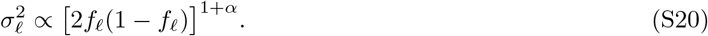

Rare variants receive relatively greater weight when *α <* −1, which amplifies the difference between marker-based and causal covariance at low relatedness. Note that *α* = −1 is the GCTA assumption, under which 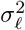 is constant across loci and (S19) reduces to *υ*_*k*_. Figure S2 shows the within-class slopes that result when every variant is both marker and causal: for *α >* −1 the slope in distant relatives exceeds *h*^2^, for *α <* − 1 it falls below, and in close relatives it returns to about *h*^2^ whatever the value of *α*.

### S3.4 Slope, intercept and phenotypic covariance

Here there is no shared environment and mating is random, so *y* = *hg* + *eϵ*. Tagging attenuates the slope by *υ*_*k*_ (S11) and leaves the intercept unchanged: in the absence of other forces, the within-class line passes through the origin:

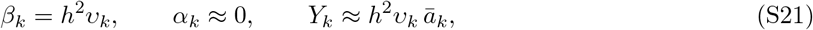

and the pooled covariance is *h*^2^*x ∑*_*k*_ *π*_*k*_(*x*) *υ*_*k*_. Under frequency-dependent effect sizes, *υ*_*k*_ is replaced by 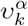 (S19).

## S4 Multi-generational shared environments

Some quantitative traits may also be shaped by non-genetic effects that run in families, which are often referred to as “shared environments”. We consider in particular factors that do not depend on genetics, such as culturally transmitted factors, geographic and environmental exposures, and socioeconomic factors. We do not consider genetic nurture or dynastic effects, in which an individual’s environment depends on its parents’ phenotypes rather than on their environments. We dub these multi-generational shared environments (MGSEs), to indicate that they may span longer genealogical distances than traditional sibling or family effects. Throughout this section mating is random; we return to assortment in §S5.

**Figure S2:**
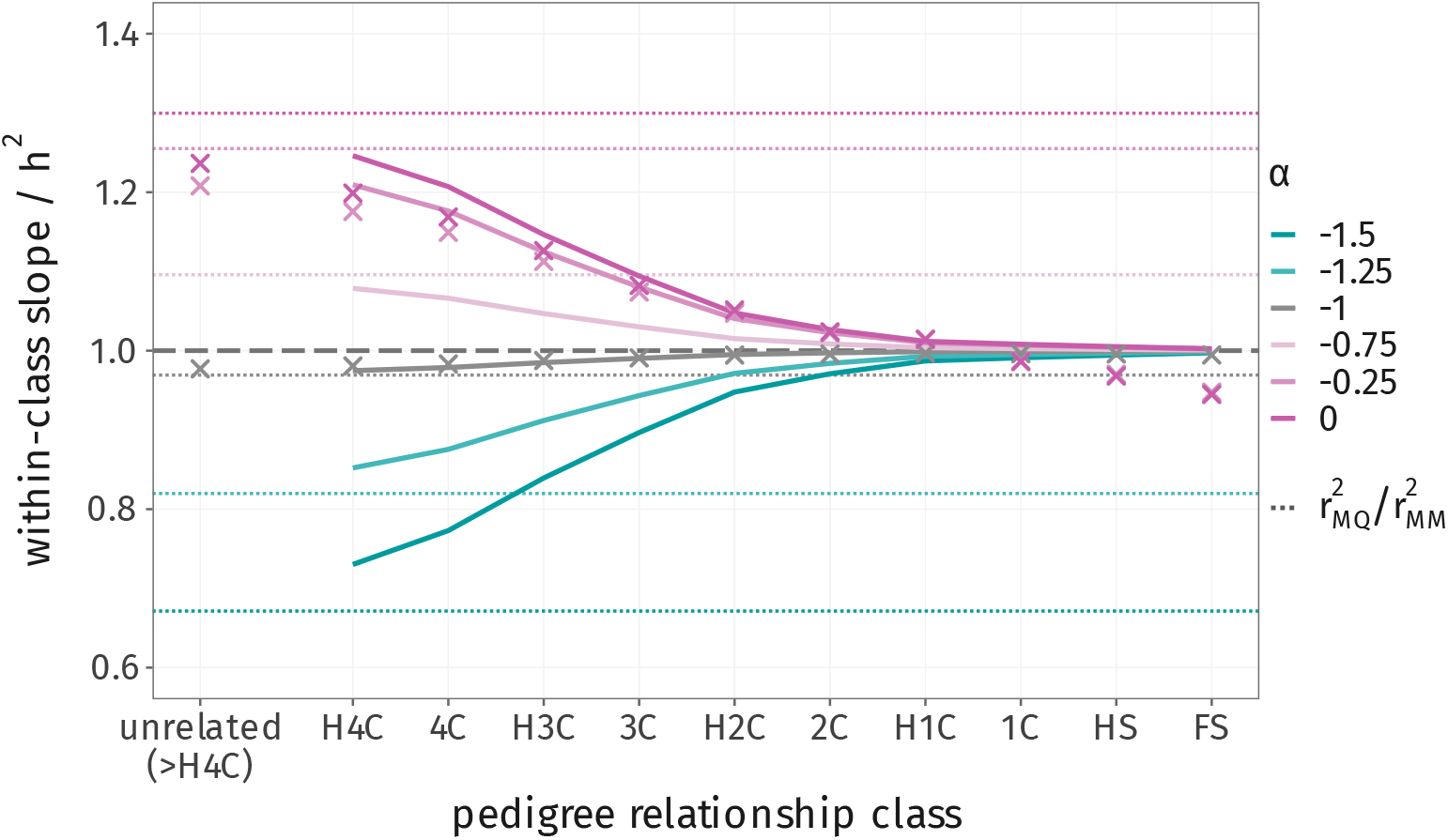
Frequency-dependent effect sizes bias the within-class slope even when every variant is genotyped. Within-class slope of phenotypic covariance on genetic relatedness, relative to *h*^2^, by pedigree class, when every variant is both marker and causal and the per-allele effect-size variance scales as [2*f*_*l*_(1 − *f*_*l*_)]^*α*^ (S20); *α* = −1 (gray) is the architecture the standard GRM assumes. Lines, theory (S19); crosses, simulation, for the values of *α* simulated; dotted, the limit among unrelated pairs, 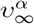; dashed, 1.

We model MGSEs by adding a C component to the phenotype, *c* in (S1), which enters with path coefficient *b*, similarly to the C of the ACE models used in twin studies (Figure S4 with *ρ*_*y*_ = 0). Given the pedigree *P*, and under random mating, the environments are independent of every genotype and of the segregation *σ*, and have mean zero.

The covariance of *c* between two individuals then depends only on their pedigree relatedness, not on their realized relatedness,

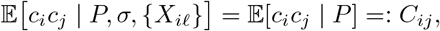

and we write *ā*_*ij*_ := E[*x* | *P*] for the pedigree relatedness of the pair (twice their kinship coefficient), so that E[*ā*_*ij*_ | *k*] = *ā*_*k*_. If every pair of class *k* has the same *C*_*ij*_, which we write *C*_*k*_ as in §S1, the expected covariance of the class is *Y*_*k*_ = *h*^2^*ā*_*k*_ + *b*^2^*C*_*k*_, since under random mating E[*g*_*i*_*g*_*j*_ | *k*] = *ā*_*k*_, and E[*g*_*i*_*c*_*j*_] = 0 because *c* is independent of the genotypes given the pedigree.

For the slope, *x* is a function of the genotypes, so the environments are independent of *x* given the pedigree. In a class whose pairs all share the same pedigree relationship, E[*c*_*i*_*c*_*j*_ | *x, k*] = *C*_*k*_, and E[*g*_*i*_*c*_*j*_ | *x, k*] = 0 because E[*c*_*j*_ | *P*] = 0, so only the genetic part of *y*_*i*_*y*_*j*_ varies with *x*. With the markers equal to the causal loci (M = Q), E[*g*_*i*_*g*_*j*_ | *x, k*] = *x*, and the shared environment enters the intercept only:

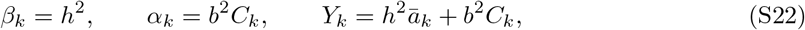

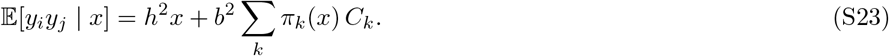

The local slope departs from *h*^2^ only where the composition of classes changes with *x*. With imperfect tagging, *β*_*k*_ and the genetic part of *α*_*k*_ are as in (S21), and *b*^2^*C*_*k*_ is added to the intercept.

### S4.1 The parental-mean model

We give a specific example of a generative model for the environment that produces MGSEs, and the covariances it generates. This is the parental-mean model of §S1, a special case of the cultural transmission model of Rice et al. (1978), with each parent’s environment transmitted with coefficient 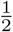. There are three reasons why this model is of particular interest. First, it results in an equilibrium variance. Second, its covariances decay with pedigree distance exactly as genetic covariances do (*C*_*k*_ = *ā*_*k*_), so that, averaged within a class, they are indistinguishable from those of an additive model with heritability *h*^2^ + *b*^2^, even without any genetic effects. In this sense it is a particularly confounding form of MGSE. Third, it considerably simplifies the results for assortative mating in §S5.

Here each offspring receives the midparent environment plus independent noise of variance 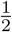 (S2), and its contribution to the phenotype is parametrized by *b*^2^ (S1). Under random mating the two parents are uncorrelated, so Var 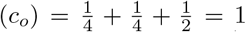, and the variance is at equilibrium in every generation. Each parent–offspring step carries a coefficient 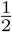, as for the breeding value, so

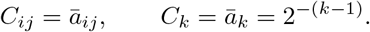

This results in a series of geometrically decaying *C*_*k*_, and lineal and collateral pairs of the same class (parent– offspring and full sibs; half sibs, avuncular and grandparent–grandchild pairs) have the same *C*_*k*_. Substituting into (S22), *β*_*k*_ = *h*^2^ and *α*_*k*_ = *b*^2^*ā*_*k*_, and the pooled covariance is (S23) with *C*_*k*_ = *ā*_*k*_.

### S4.2 Other environmental structures

(S23) holds for any *C*_*k*_, and under random mating the class composition *π*_*k*_(*x*) does not depend on the environment, so the pooled covariance for another structure follows from the same simulated pairs. As an extra example, Figure S3 compares the parental-mean model with a family of ancestral-household environments. Each household carries an independent effect, and an individual’s environment averages, with equal weight for each generation *d* = 0, …, *D*, the households in which its ancestors *d* generations back grew up (*d* = 0 is the individual’s own household), scaled to unit variance. Two relatives of class *k* first share a household at depth *x*_*k*_ = ⌊ (*k* − 1)*/*2⌋ (full sibs at depth 0; half sibs and first cousins at depth 1; half first and second cousins at depth 2; and so on), and from there on share a fraction 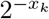 of their ancestors’ households, so that

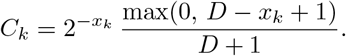

*D* = 0 is a household effect shared only by full sibs. With *D* = 0 the pooled covariance follows *h*^2^*x* up to the full-sibling range; each further generation extends the shared environment to more distant classes.

**Figure S3:**
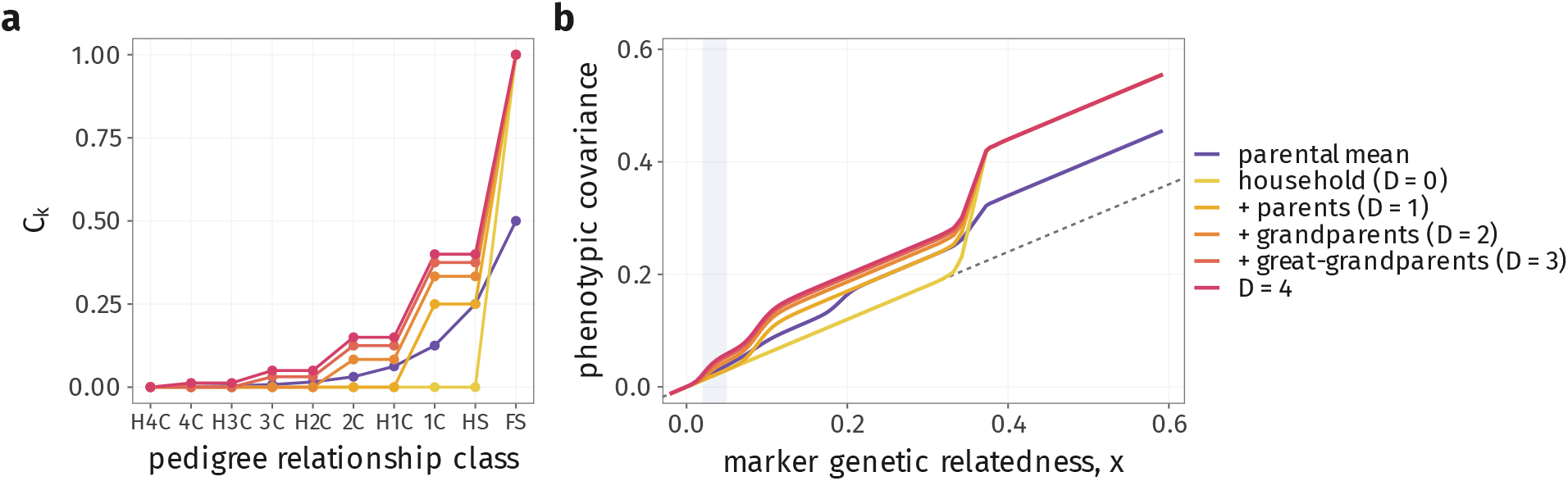
The shape of the pooled covariance depends on how far the shared environment extends. Random mating, *h*^2^ = 0.6, *b*^2^ = 0.2. (a) *C*_*k*_ by class under the parental-mean model (*ā*_*k*_) and under ancestral-household environments of depth *D* = 0 (household) to *D* = 4 (§S4.2). (b) Pooled mean standardized cross-product under S1; dashed, *h*^2^*x*. Theory, (S23), with *π*_*k*_(*x*) from the simulated pairs. Gray band as in Figure 2.

## S5 Assortative mating

Now we look at the relationship between phenotypic covariance and relatedness under phenotypic assortment parametrized by *ρ*_*y*_, the phenotypic correlation between mates. Assortment changes two things: each component’s variance is inflated above its random-mating value, and *g* and *c* acquire a covariance *ω*^∗^ that is zero under random mating. Throughout this section, *c* is transmitted by the parental-mean model (§S4.1). The results for *b* = 0 do not depend on it; those involving *c* do, namely 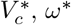 and *ρ*_*c*_, and through them 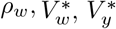 and *Y*_*k*_ when *b >* 0. In the main text, results specific to this model, such as 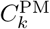, carry a superscript PM.

### S5.1 Path covariances

Figure S4 shows the paths connecting two mates and two of their children. Within an individual, the phenotype loads on *g, c* and *ϵ* through *h, b* and *e* (S1), and *g* and *c* covary at *ω*^∗^. Mates are connected only through their phenotypes, with correlation *ρ*_*y*_. Each component passes to the offspring with coefficient 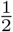 from each parent, plus the independent terms *m* and *ξ* (S2); for *c*, this is the parental-mean model.

**Figure S4:**
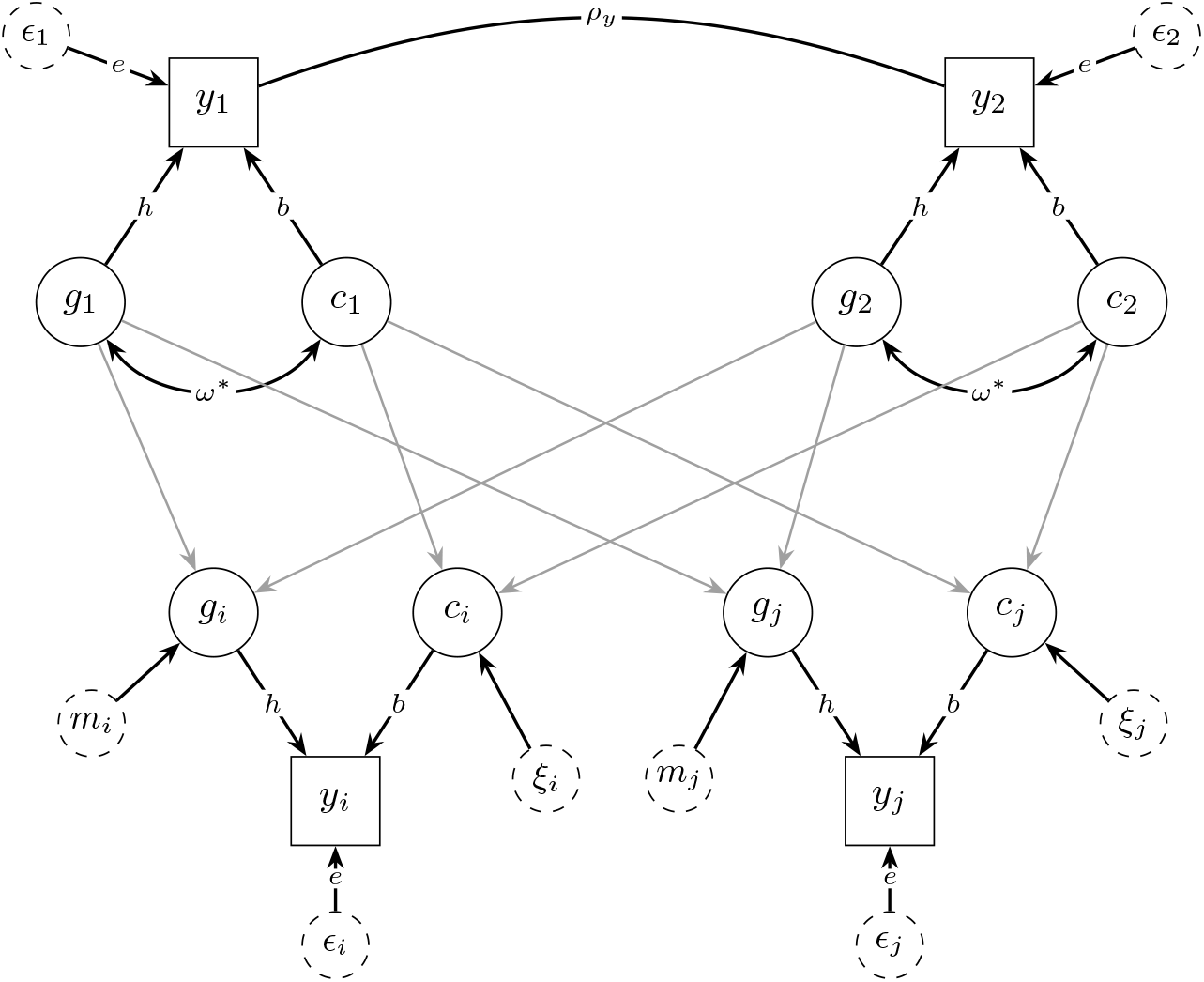
Path diagram for two mates and two of their children, *i* and *j*. Squares are observed phenotypes, circles latent components, dashed circles independent noise. Single-headed arrows are causal paths; the gray arrows from parents to children have coefficient 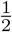, and the unlabeled arrows from *m* and *ξ* have coefficient 1. Double-headed arrows are covariances between exogenous variables (the parents’ *g* and *c*, of variances 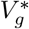 and 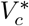). The undirected line is a *copath* (Cloninger, 1980). *Mates are chosen on their phenotypes, so this is not an ordinary correlation path: by Wright’s tracing rules (Wright, 1934) a two-headed arrow between y*_1_ and *y*_2_ would leave *g*_1_ and *g*_2_ uncorrelated. Instead, a chain may cross the copath only from one mate’s phenotype to the other’s, and it contributes 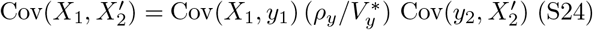 (S24). Chains may pass from one copath to another through a shared phenotype, as for two mates of the same person. *m* and *ξ* have variances 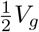 and 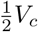(S2). Under random mating (*ρ*_*y*_ = 0, so that *ω*^∗^ = 0), this is the parental-mean model of §S4.1.

Tracing the paths gives every covariance we need; Table S2 collects them. Within an individual, the covariances of the phenotype with its genetic and shared-environment components are 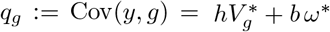 and 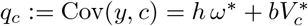. Between mates, which are connected only through *y*_1_ and *y*_2_, the covariances of the components are derived in §S5.2.

**Table S2:** Path covariances. *h, b* and *e* are standardized under random mating only; at equilibrium the standardized path from *g* to *y* is 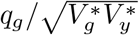.

| Quantity | Equilibrium value | Random mating |
| --- | --- | --- |
| *Within an individual* |  |  |
| $\text{Var}(g)$ | $V_g^*$ | $V_g = 1$ |
| $\text{Var}(c)$ | $V_c^*$ | $V_c = 1$ |
| $\text{Cov}(g, c)$ | $\omega^*$ | 0 |
| $\text{Var}(w)$ | $V_w^* = h^2 V_g^* + b^2 V_c^* + 2hb\omega^*$ | $h^2 + b^2$ |
| $\text{Var}(y)$ | $V_y^* = V_w^* + e^2$ | 1 |
| $\text{Cov}(y, g)$ | $q_g = hV_g^* + b\omega^*$ | $h$ |
| $\text{Cov}(y, c)$ | $q_c = h\omega^* + bV_c^*$ | $b$ |
| $\text{Cov}(y, w)$ | $V_w^*$ | $h^2 + b^2$ |
| *Between mates* |  |  |
| $\text{Cov}(y_1, y_2)$ | $\rho_y V_y^*$ | 0 |
| $\text{Cov}(g_1, g_2)$ | $\rho_g V_g^*$ (S25) | 0 |
| $\text{Cov}(c_1, c_2)$ | $\rho_c V_c^*$ (S25) | 0 |
| $\text{Cov}(g_1, c_2)$ | $\rho_{gc} \sqrt{V_g^* V_c^*} = \omega^*$ (S26) | 0 |
| $\text{Cov}(w_1, w_2)$ | $\rho_w V_w^*$ (S29) | 0 |
| *Parent to offspring* |  |  |
| $\text{Cov}(g_1, g_o)$ | $\frac{1}{2} V_g^* (1 + \rho_g)$ | $\frac{1}{2}$ |
| $\text{Cov}(c_1, c_o)$ | $\frac{1}{2} V_c^* (1 + \rho_c)$ | $\frac{1}{2}$ |

### S5.2 Mate correlations of the trait components

Mates are matched on the phenotype only. Therefore, given the phenotypes, each mate’s components are independent of the other’s, and any covariance between them must pass through *y*_1_ and *y*_2_ (Wright, 1921). For *X* measured on mate 1 and *X*^*′*^ on mate 2, regressing each on its own mate’s phenotype and using 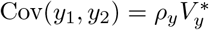 gives, at equilibrium,

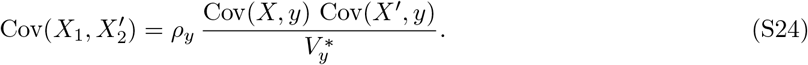

Joint normality is not needed: linearity of the regressions of each component on the phenotype, and of one mate’s phenotype on the other’s, is sufficient (Nagylaki, 1982). Substituting *q*_*g*_ and *q*_*c*_ into (S24) gives the three mate covariances of the components:

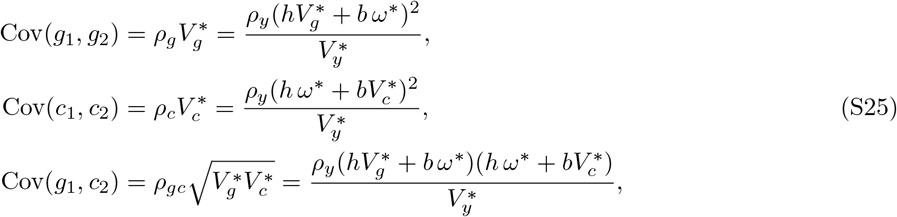

where *ρ*_*g*_, *ρ*_*c*_ and *ρ*_*gc*_ are the mate correlations. Note that written like this, the path coefficients are not standardized; in order to be, they need to be divided by the square root of the product of the respective equilibrium variances, 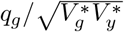 This generalizes equation (5) of Rice et al. (1978), which has a single transmissible factor, to two components and their cross-covariance. For *b* = 0 it gives 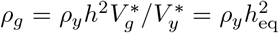, the mate correlation of breeding values under the equilibrium heritability (Border et al., 2022, Supplementary Note 1; Veller and Coop, 2024, S1 Text, Eq. S.45).

### S5.3 Gene–environment covariance under the parental-mean model

Assortment introduces a gene–environment covariance. Under the parental-mean model of environmental transmission, we can write *ω*_*t*+1_ = Cov(*g*_*o*_, *c*_*o*_) and expand:

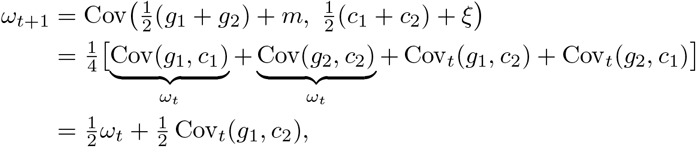

where Cov_*t*_ denotes a covariance between the mates of generation *t*, the noise terms dropping out because *m* and *ξ* are independent of everything else. Two of the four terms are within-parent and two between. Setting *ω*_*t*+1_ = *ω*_*t*_ = *ω*^∗^ gives the fixed point,

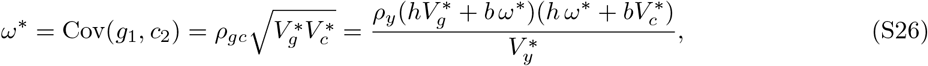

so the within-person correlation 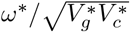 equals the mate correlation *ρ*_*gc*_. We solve for the equilibrium variances in the next subsection (for genic–cultural covariance under assortment, cf. Cloninger et al., 1979).

### S5.4 Variance inflation

The genic variance is defined as 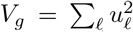. At meiosis, Mendelian sampling contributes some of this variance to the next generation: 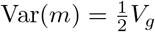. Taking the variance of offspring under assortment, and with the equilibrium parent covariance 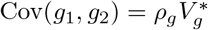 (S25),

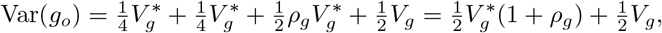

and setting 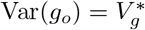 — the equilibrium condition — gives 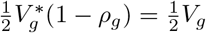, so

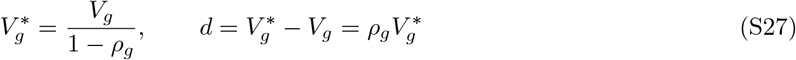

(Fisher, 1919; Wright, 1921; Crow and Felsenstein, 1968; Nagylaki, 1982). Under the parental-mean model, with 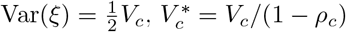 analogously.

Since 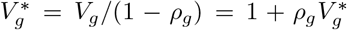 and likewise for *c* under the parental-mean model, while *ω*^∗^ = Cov(*g*_1_, *c*_2_) by (S26), substituting the mate covariances (S25) gives

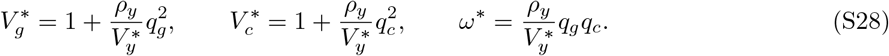

It is also useful to write the variance of the whole transmissible component of the phenotype, *w* = *hg* +*bc*, composed of the genetic, shared-environment, and gene–environment components, so that 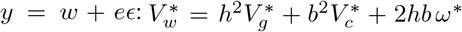 (Table S2), and 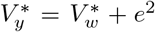. Under panmixia 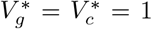 and *ω*^∗^ = 0, giving 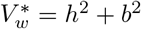.

Since 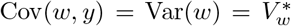, applying (S24) to *w* gives the mate correlation of this transmissible part:

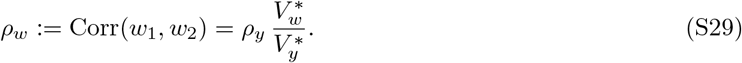

The transmissible part *w* is passed on as a parental mean with independent noise *hm* + *bξ*, of variance 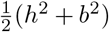 (cf. the pseudopolygenic case of the unitary model of Rice et al., 1978), so the same argument as (S27) gives

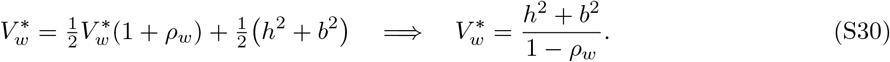

Substituting 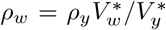 and 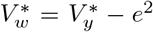 into (S30), multiplying through by 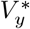 and using *h*^2^ + *b*^2^ + *e*^2^ = 1 gives the quadratic 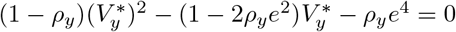, whose positive root is

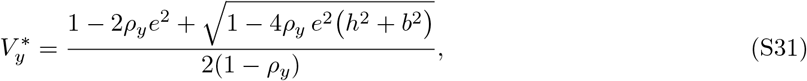

equal to 1 at *ρ*_*y*_ = 0, with 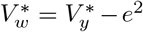. Substituting these into (S28) gives closed forms. First, substituting (S28) into 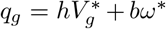 gives 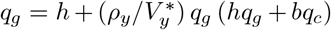, and 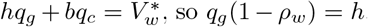. The same holds for *q*_*c*_:

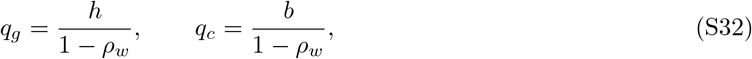

and putting these back into (S28),

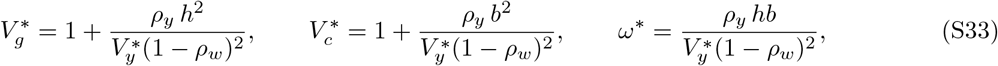

after which *ρ*_*g*_, *ρ*_*c*_ and *ρ*_*gc*_ follow from (S25).

Figure S5a follows the variances and the gene–environment covariance across the simulated generations: they reach the equilibrium values of (S33) within about ten generations for *ρ*_*y*_ ≤ 0.4, while for *ρ*_*y*_ = 0.6 they are still rising at generation 20. Figure S5b shows the equilibrium mate correlations of the components (§S5.2).

### S5.5 Covariance between relatives

#### S5.5.1 Descendants of full siblings

We will work through expressions for the descendants of full sibs first (the sibs *i* and *j* of Figure S4). The two sibs draw from the same parental pair, and their Mendelian terms *m*_*i*_, *m*_*j*_ are independent of each other and of the parents:

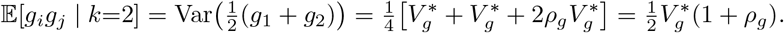

**Figure S5:**
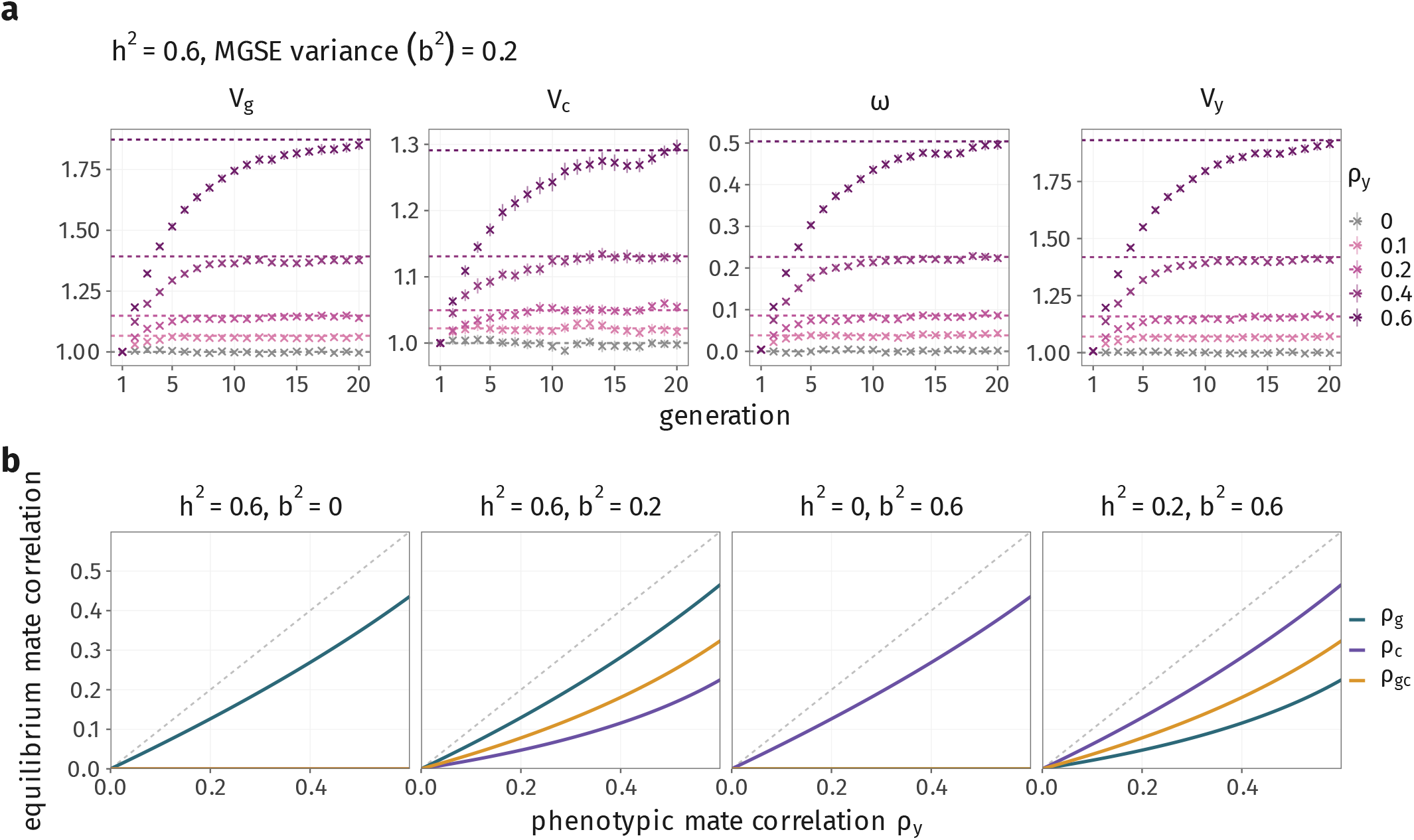
Assortative mating reaches equilibrium within the simulated generations, except at *ρ*_*y*_ = 0.6. (a) Genetic variance Var(*g*), shared-environment variance Var(*c*), gene–environment covariance Cov(*g, c*) and phenotypic variance Var(*y*), all on the random-mating scale, in each of the 20 simulated generations, for *h*^2^ = 0.6, *b*^2^ = 0.2 and *ρ*_*y*_ from 0 to 0.6 (crosses, mean over 50 replicates, 95% intervals); dashed, the equilibrium of (S33) and (S31). (b) Equilibrium mate correlations of the genetic (*ρ*_*g*_) and shared-environment (*ρ*_*c*_) components, and between one mate’s genetic and the other’s shared-environment component (*ρ*_*gc*_), against *ρ*_*y*_, for four architectures; dashed, *y* = *ρ*_*y*_.

The same calculation on *c* gives 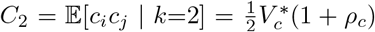, and for the cross-component term, using Cov(*g, c*) = *ω*^∗^ (S26), 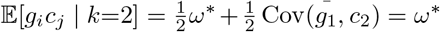. Parent–offspring pairs give the same three values. These are covariances of the transmissible components only. The phenotypic cross-product of a parent and offspring also picks up the parent’s own environment, which assortment correlates with the mate’s genotype, so 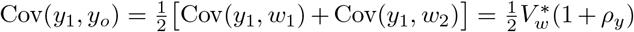, against 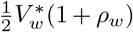 for full sibs, as in the lineal results of Yengo and Visscher (2018). The classes we consider are all collateral, so this does not enter our results.

A descent step replaces *j* by an offspring *o*_*j*_ of *j* and *j*’s mate *p*_*j*_, so for components *X, X*^*′*^ ∈ {*g, c*}

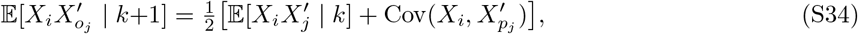

and *p*_*j*_ reaches *i* only through *y*_*j*_. Both terms are therefore built from within-individual covariances and the between-mate covariances. Writing

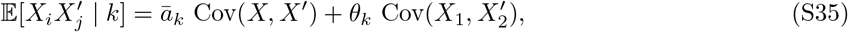

the in-law term follows from (S24). Since *ϵ*_*j*_ is independent of *i*, Cov(*X*_*i*_, *y*_*j*_) = Cov(*X*_*i*_, *w*_*j*_), and applying (S35) with Cov(*X, w*) = Cov(*X, y*) and Cov(*X*_1_, *w*_2_) = *ρ*_*w*_ Cov(*X, y*) gives

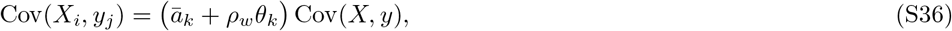

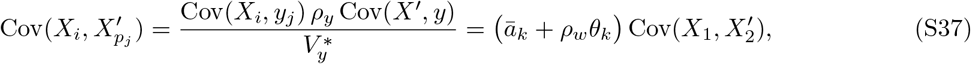

the second by (S24).

Substituting (S35) and (S37) into the descent step (S34), the within-individual and between-mate terms separate, leaving the recursion 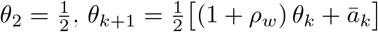, whose solution is

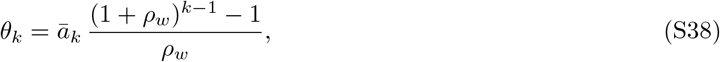

which at *ρ*_*w*_ = 0 is read as its limit, *θ*_*k*_ = *ā*_*k*_(*k* − 1), so that

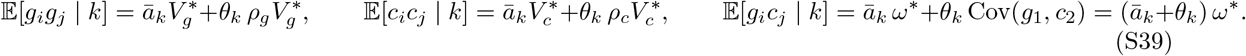

Applying the descent step on both sides of the pedigree gives the first cousins *o*_*i*_ and *o*_*j*_ (*k* = 4), and further steps give more distant cousins. Taking *X* = *X*^*′*^ = *w*, with 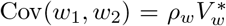, the expected phenotypic cross-product is

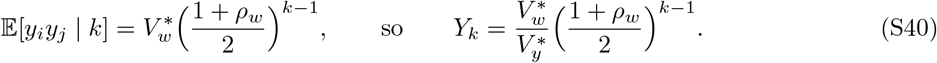

*For b* = 0, *ρ* = *ρ* and the first entry of (S39) reduces to the classical 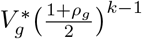 (Fisher, 1919; Nagylaki, 1978; Gimelfarb, 1981; Yengo and Visscher, 2018; Sunde et al., 2024).

#### S5.5.2 Descendants of half siblings

Half sibs *i* and *j* share a parent, 1, and their other parents, 2 and 3, are both mates of 1, so these two mates are themselves correlated at 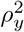. Expanding as for full sibs,

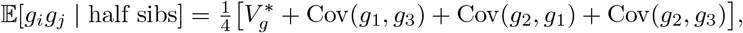

where 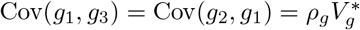 are mate covariances and, since 2 and 3 are connected only through *y*_1_,

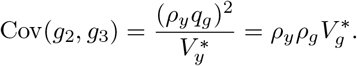

This gives

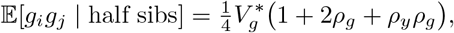

as in Nagylaki (1978) and Yengo and Visscher (2018), against 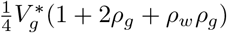for an avuncular *k* = 3 pair from (S39). The two differ by 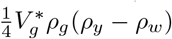, where 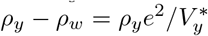 (S29). The same calculation for the whole transmissible component *w* gives 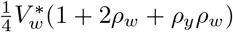 for half sibs, which exceeds the *k* = 3 value 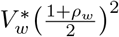 by the fraction

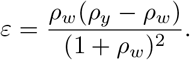

Each descent step (S34) multiplies the cross-product of *w* by 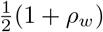, so *ε* is also the relative error in the class means of the descendants of half sibs. It vanishes when *e*^2^ = 0 (so that *ρ*_*w*_ = *ρ*_*y*_) and as *h*^2^ + *b*^2^ → 0 (so that *ρ*_*w*_ → 0), and is largest at *ρ*_*w*_ = *ρ*_*y*_*/*(2 + *ρ*_*y*_), so that 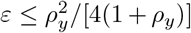 whatever *h*^2^ and *b*^2^: 0.8% at *ρ*_*y*_ = 0.2, 2.9% at *ρ*_*y*_ = 0.4 and 5.6% at *ρ*_*y*_ = 0.6. For *h*^2^ = 0.6 and *b*^2^ = 0 it is 2.2% at *ρ*_*y*_ = 0.4 and 3.5% at *ρ*_*y*_ = 0.6. Under the parental-mean model *ε* depends on *h*^2^ and *b*^2^ only through *h*^2^ + *b*^2^, and it is largest for traits with a modest transmissible share, at *h*^2^ + *b*^2^ ≈ 0.3 for *ρ*_*y*_ = 0.6 (4.9% at *h*^2^ + *b*^2^ = 0.2). Since the discrepancy is small over the range of *ρ*_*y*_ considered here, we treat half sibs as an ordinary *k* = 3 class throughout, which lets us treat them and their descendants with the same equation as the descendants of full sibs. This is an approximation, and it affects only the class means *Y*_*k*_ of half sibs and their descendants.

### S5.6 Slope and intercept of regression on relatedness

By definition, the regression slope of the standardized cross-product on relatedness is 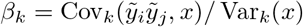 (S3). With Cov(*R, η*) = 0, it is a simple weighted average of two slopes:

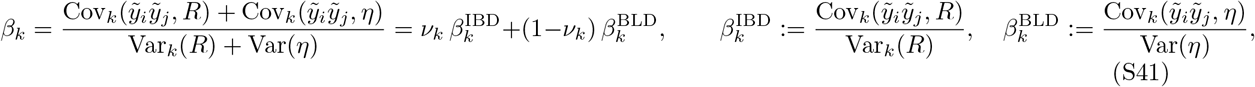

with *ν*_*k*_ as in (S9). We derive the covariances below for the unstandardized phenotype *y* and divide by 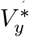 (S31). We show below that neither slope depends on the class (for siblings, and approximately for the BLD slope), and write them *β*^IBD^ and *β*^BLD^, as in the main text:

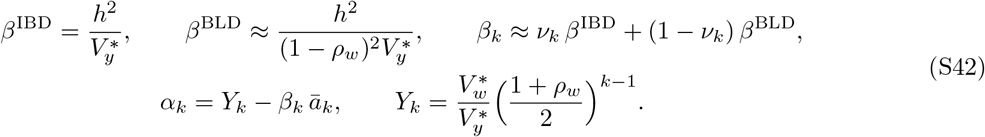

The IBD slope is exact for siblings and understates the slope for cousins (§S5.6.1); the BLD slope is derived in §S5.6.2, and the class mean *Y*_*k*_ in (S40). Without assortment, *ρ*_*w*_ = 0 and 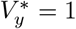, and every slope reduces to *h*^2^.

#### S5.6.1 The IBD portion of the slope

##### Genetic component

In order to derive the slope of the genetic component on IBD, we first split the genetic trait component into same- and cross-locus sums,

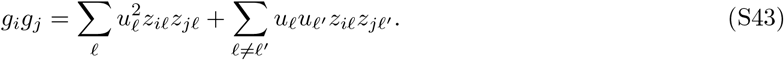

The first has E[*z*_*il*_*z*_*jl*_ | *σ*] = *R*_*l*_, the realized IBD at locus *l* (*R*_*l*_ = *S*_*l*_*/*2 in (S6)). For the second part, we can write 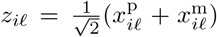, where 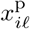and 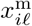 are the standardized paternal and maternal alleles, 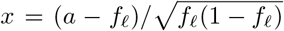for an allele indicator *a* ∈ {0, 1}, so that 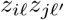 is 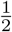 times a sum of two same-parent and two cross-parent allele products:

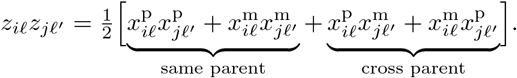

We can separate the LD into cis and trans components, representing the covariance between alleles on the *same* haplotype, and the covariance between alleles on an individual’s different haplotypes. Let 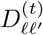 be *cis*-LD in generation *t*, and 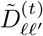 be *trans*-LD. Looking at a pair of alleles in a full-sib pair, let *χ* = 1 if they came from a parent’s same haplotype and *χ* = 0 if they came from two different ones in the same parent, with *χ*^p^ and *χ*^m^ for the paternal and maternal terms. Then 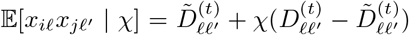. A cross-parent term is the covariance between the two parents’ transmitted alleles, which is trans-LD and does not depend on the segregation.

If we collect these terms,

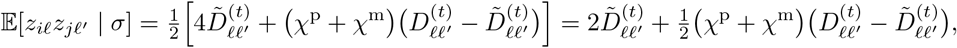

the 2 being 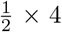 terms each contributing 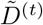. This holds in any generation. A transmitted gamete carries the parent’s cis-LD when no crossover separates *l* and *l*^*′*^ and its trans-LD otherwise, so 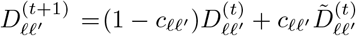, and at the equilibrium under assortment, for any 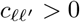,

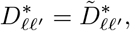

(Nagylaki, 1982; Veller and Coop, 2024), which is why the equilibrium correlations between relatives do not depend on the linkage map (Nagylaki, 1978). The *χ* term therefore vanishes and 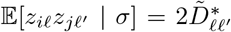, a constant. For more distant relatives the constant takes a different value, but the argument is the same: at every meiosis on the path the choice of haplotype is multiplied by 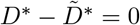. This removes every way in which *σ* enters for siblings. For more distant relatives there is a second way, through the in-laws who marry into the pedigree; we return to it at the end of this section.

Recalling that *σ* denotes the segregation at every meiosis in the pedigree (§S2), and taking the conditional mean of (S43),

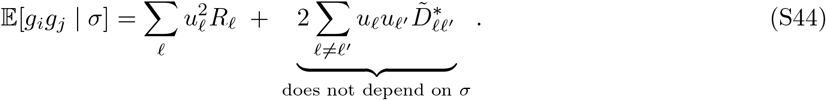

This means that the *conditional mean given σ* of the cross-locus sum is constant. Since the IBD, *R*, is a function of *σ*, the tower property gives, for any *W*,

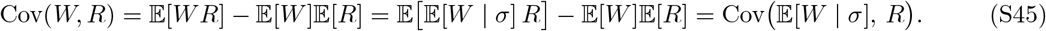

Intuitively, two individuals may share many trait-increasing alleles across different loci by chance, without having inherited a longer stretch of genome in common (i.e., they happen to share many alleles identical by state, in tracts that are not IBD); that similarity contributes to *g*_*i*_*g*_*j*_, but it does not depend on the particular meioses that took place, so it does not covary with *R* and cannot appear in the IBD portion of the slope.

Applying (S45) to *W* = *g*_*i*_*g*_*j*_ and substituting (S44), the constant term drops out of the covariance and

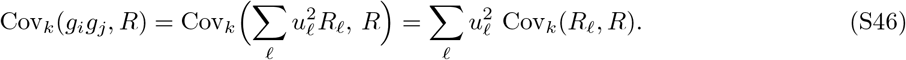

Since 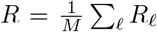, Var_*k*_(*R*) is the *unweighted* average of Cov_*k*_(*R*_*l*_, *R*) while (S46) is its *u*^2^-weighted average. If causal loci are spread over the map like markers, the two agree and

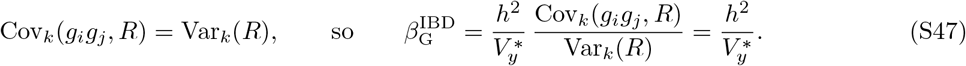

However, this is an assumption; some ways it might fail are

- if causal variants are clustered on part of the map, so that their Cov_*k*_(*R*_*l*_, *R*) is systematically above or below the marker average;
- if effect size covaries with local recombination rate, since Cov_*k*_(*R*_*l*_, *R*) is larger in regions of low recombination;
- under a frequency-dependent effect-size model (§S3.3), where the *u*^2^ weights track allele frequency and hence, through ascertainment, the marker distribution. We return to this later in §S6.

##### Environmental components

The shared environment is transmitted passively (§S1): *c*_*i*_ and *c*_*j*_ are functions of ancestral *c*’s and of the *ξ*’s alone, none of which depend on which haplotypes were transmitted. For full sibs under the parental-mean model, write 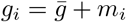 and 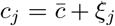, with 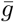 and 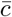 the midparent values. Every term of *c*_*i*_*c*_*j*_, and of 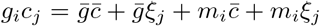, has a conditional mean given *σ* that does not depend on *σ*: 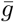 and 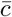are fixed before the meioses, and *m*_*i*_ and *ξ*_*j*_ have mean zero given the segregation. For more distant relatives the same holds at each meiosis along the line of descent, but not for an in-law, whose *c* depends on *σ* through the phenotype of the line member it mated with; we treat in-laws in §S5.6.1. By (S45),

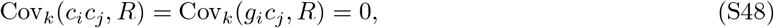

so 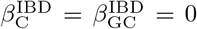: the shared environment and the gene–environment term enter the class mean, and hence the intercept, but not the IBD portion of the slope.

##### The IBD limit for siblings

Expanding the phenotypic cross-product,

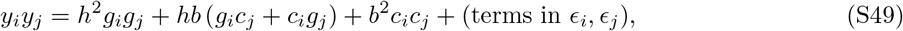

where the *ϵ* terms are independent of *σ*. If we regress on IBD alone, the gene–environment and environmental terms drop by (S48), leaving only the genetic one:

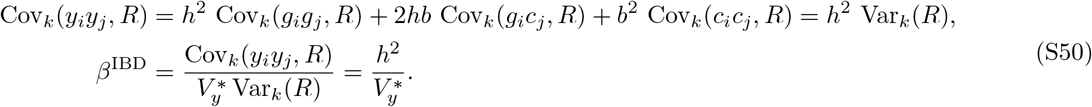

For siblings this holds for any strength of assortment. It is the random-mating genetic variance over the equilibrium phenotypic variance, and lies below *h*^2^ because 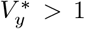. For 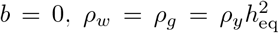with 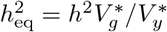, and since 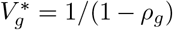 (S27), 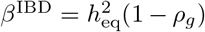 the expectation of regression on IBD between full sibs in Kemper et al. (2021, Supplementary Note 3) (see also Visscher et al., 2006).

##### Cousins

This holds for full sibs and half sibs. For cousins it understates the true slope, because of the in-laws, the spouses who married into the two lines of descent. Each was chosen on the phenotype of the line member it married, and so covaries with the segments that line member shares IBD with the other line. For first cousins *o*_*i*_ and *o*_*j*_, the children of the sibs *i* and *j* and their mates *p*_*i*_ and *p*_*j*_, *p*_*i*_ reaches the other line only through the copath to *y*_*i*_. By (S24), Cov(*w*_*p*_*i, W* | *σ*) = *ρ*_*w*_ Cov(*y*_*i*_, *W* | *σ*) for any *W* on that line, and *o*_*i*_ carries half of *p*_*i*_’s *w*. Applying this at both in-laws, and (S44) and (S48) to the relatives on the two lines,

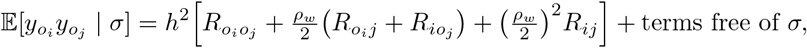

where *R*_*xy*_ is the IBD of *x* and *y*. For instance, the chain from 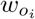 to *y*_*j*_, across the copath to 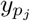 and down through 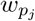 to 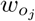 gives the in-law term 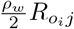. A segment that *o*_*i*_ shares with *j* reaches *o*_*j*_ with probability 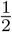, and one the sibs share reaches both cousins with probability 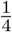 so 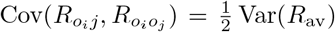 and 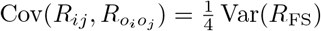 By (S45),

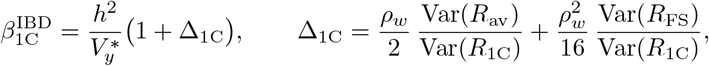

where av, 1C and FS denote avuncular, first-cousin and full-sib pairs. This is exact under the model, for any *b*: because each in-law was chosen on *y*, the weight *ρ*_*w*_*/*2 passes partly through its *g* (a fraction *h*^2^*/*(*h*^2^ + *b*^2^)) and partly through its *c* (a fraction *b*^2^*/*(*h*^2^ + *b*^2^); Eq. (S32)), and both are included in Δ_1C_. In cousins the shared-environment and gene–environment terms therefore also covary with IBD, and (S48) holds for siblings only. The variance ratios follow from the map integral (S17); simulated pedigrees on our recombination map give about 1.37 and 3.15, so that 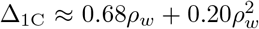. Avuncular pairs, which have one in-law on the path, are understated in the same way, while lineal pairs such as grandparent and grandchild are not.

The same holds for more distant cousins, whose paths carry more in-laws, but we do not derive it here. The slopes we predict for cousins, which use 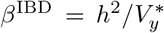, are therefore too low, and their intercepts *α*_*k*_ = *Y*_*k*_ − *β*_*k*_*ā*_*k*_ too high. Figure S7 shows the size of this error in simulation. At *ρ*_*y*_ = 0.4 (*h*^2^ = 0.6, *b*^2^ = 0.2), the simulated slopes of half first to half fourth cousins exceed the prediction by 0.45 *h*^2^ on average (95% interval 0.36–0.54 *h*^2^; inverse-variance weighted over the seven classes), about 40% of the predicted slope. The gap is 0.11 *h*^2^ (0.03–0.18) at *ρ*_*y*_ = 0.2 and indistinguishable from zero under random mating. For first cousins, adding Δ_1C_ raises the predicted slope from 0.74 *h*^2^ to 0.91 *h*^2^, against 0.93 *h*^2^ in simulation. Even so, the pooled predictions fit the simulations well: the class means are exact, and each within-class line passes through its class mean (S42), so a slope error only tilts the line about that point. Working out the slopes of more distant cousins is left for further work.

#### S5.6.2 The BLD portion of the slope

Unlike the IBD results, the results of this subsection are approximate. They treat the standardized genotypes as jointly normal with the trait components, take 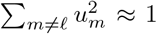, assume the markers are the causal loci, and finally drop the across-pair term 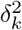. Among unrelated pairs, the simulated slopes lie 0.20 *h*^2^ above the resulting BLD limit at *ρ*_*y*_ = 0.4, a limit that assumes the markers are the causal loci (*M*= *Q*; Figure S7). Under random mating they lie 0.13 *h*^2^ below *h*^2^. This reflects mild imperfect tagging: the marker GRM keeps SNPs above 1% minor allele frequency, whereas 24% of the causal sites are rarer, so *υ*_*∞*_ ≈ 0.9 in these simulations.

In general, we can use Isserlis’ identity (Isserlis, 1918) to write the covariance of cross-products approximately. For zero-mean, jointly normal variables *W*_1_, …, *W*_4_ it gives

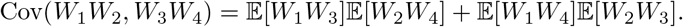

Taking 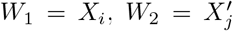for components *X, X*^*′*^ ∈ {*g, c*} and *W*_3_ = *z*_*il*_, *W*_4_ = *z*_*jl*_, and treating the standardized genotypes as jointly normal with the components,

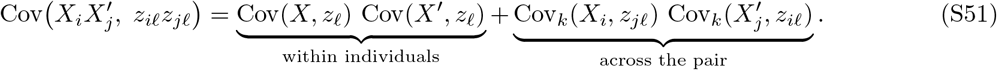

Since *i* reaches locus *l* of *j* only through *j*’s breeding value, every across-pair factor goes through *g*,

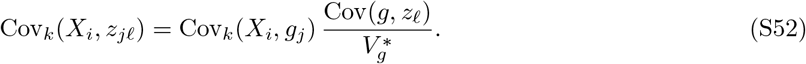

The second factor is a within-individual term, so all the dependence on *k* lies in the first factor. We treat the within-individual terms component by component, and collect the across-pair terms in the BLD limit.

##### Genetic component

In order to derive the slope of the genetic component on BLD, we first need the covariance between an individual’s breeding value and its own genotype at *l*. Expanding *g* = ∑_*m*_ *u*_*m*_*z*_*m*_,

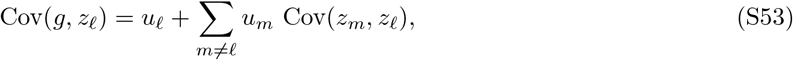

so it also picks up every other causal locus that *l* is in disequilibrium with. Under assortment the LD between causal loci is proportional to their effects, 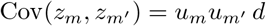, which reproduces the between-locus variance 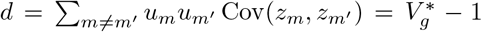 of (S27) (Nagylaki, 1982); the same structure arises under selection (Bulmer, 1971). With 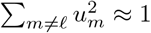

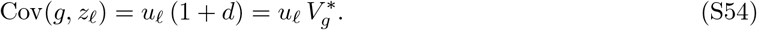

Taking *X* = *X*^*′*^ = *g* in (S51), the within-individual term at locus *l* is Cov(*g, z*_*l*_)^2^. However, this describes a single locus; the regression we are interested in uses the whole GRM. Since 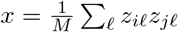 is a sum, we can write:

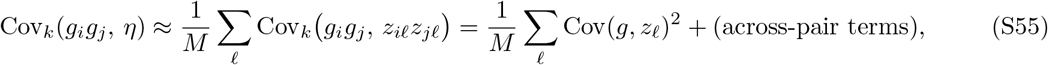

where the middle expression is strictly the covariance with *x*; the part of it carried by *R* matters only where *ν*_*k*_ is close to one (close relatives), and there the BLD portion has little weight in (S41) because the variance in IBD is much larger than that of BLD.

The denominator is Var(*η*) = 1*/M*_*e*_ (S7), which carries all the LD between markers. Background LD between the causal loci enters the numerator (S55) in the same way, through Cov(*g, z*_*l*_) (S53), so when the markers are the causal loci and the effect-size model is appropriate (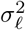 constant, §S3.3), it cancels from the ratio, as for Haseman–Elston regression under random mating, and we need only the LD induced by assortment. Summing (S54) with 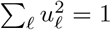

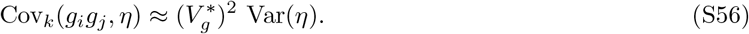

We discuss what happens when the causal loci are not well tagged by the markers in §S6. This inflation of the BLD slope is the source of the upward bias of marker-based heritability estimators under assortative mating described by Border et al. (2022).

##### Environmental components

Since *c* covaries with *z*_*l*_ only through *g*,

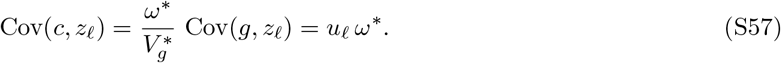

Taking *X* = *X*^*′*^ = *c*, and *X* = *g, X*^*′*^ = *c*, in (S51) and summing over loci as for the genetic component,

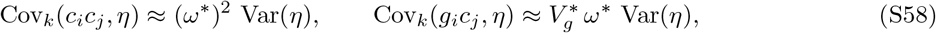

and the same for *c*_*i*_*g*_*j*_. Unlike their IBD counterparts (S48), both vary with BLD: through *ω*^∗^, the shared environment tags the genotypes that *η* reads. Both vanish without assortment, when *ω*^∗^ = 0.

##### The BLD limit

If we regress on the BLD part alone, expanding the phenotypic cross-product as in (S49), all three terms survive by (S56) and (S58):

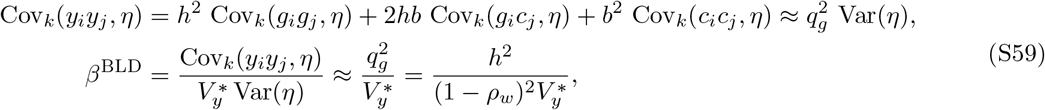

the last step by 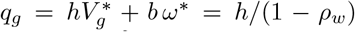 (S32). This exceeds *h*^2^ under positive assortment: by (S30), 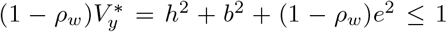, so. 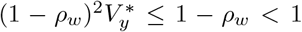 For *b* = 0, *ρ*_*w*_ = *ρ*_*g*_ and 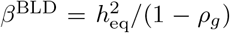, the expected HE regression estimate among unrelated individuals under assortment derived by Border et al. (2022, Eq. 2, for exchangeable loci), 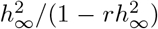 in their notation, where *r* is the phenotypic mate correlation and the subscript ∞ denotes equilibrium (so their 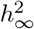 is our 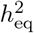). The two limits for siblings and for unrelated pairs therefore lie on opposite sides of *h*^2^, and differ by the factor 1*/*(1 − *ρ*_*w*_)^2^.

##### Across-pair terms

It remains to justify dropping the across-pair terms. Write *q*_*l*_ = Cov(*y*_*i*_, *z*_*il*_) for the covariance between an individual’s phenotype and its own genotype at *l*. By (S54) and (S57),

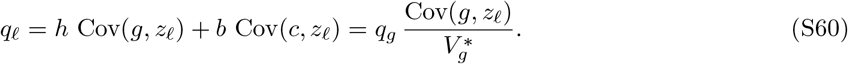

Weighting the across-pair terms of (S51) as in (S49), by bilinearity they combine into E[*y*_*i*_*z*_*jl*_] E[*y*_*j*_*z*_*il*_], and by (S52),

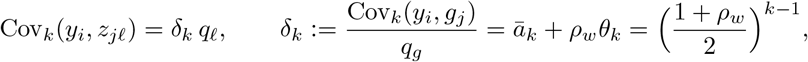

the last step by (S36) with *X* = *g* and (S38). The full BLD slope is therefore

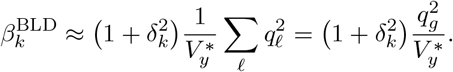

We drop 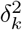 because it enters (S41) multiplied by (1 − *ν*_*k*_), which is smallest where *δ*_*k*_ is largest, so its contribution to *β*_*k*_ is small across the range of classes considered here. The term is present even without assortment, when *δ*_*k*_ = *ā*_*k*_: for half siblings it raises the BLD part of the slope by about 6%, but that part carries little weight there. Without it, like the sibling IBD slope, the BLD slope does not depend on *k*.

#### S5.6.3 Total slope

Substituting *β*^IBD^ (S50) and *β*^BLD^ (S59) into (S41) gives the total slope on GRM relatedness in (S42). Table S3 (top) gives it component by component. The weighted average (S41) is exact up to Var_*k*_(*η*) ≈ Var(*η*) (S8); otherwise the approximation enters only through *β*^BLD^. For cousins the IBD part understates the slope (§S5.6.1). Without assortment *ρ*_*w*_ = 0 and 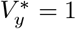, so *β*^IBD^ = *β*^BLD^, *ν*_*k*_ drops out, and *β*_*k*_ = *h*^2^ for every *k*.

#### S5.6.4 The intercepts

We derive the intercepts using the slope and the class covariance. Since the regression passes through the class means, *α*_*k*_ = *Y*_*k*_ − *β*_*k*_*ā*_*k*_ (S3), with *Y*_*k*_ by (S40). Component by component, the class covariances follow from (S39) with the weights of (S49); Table S3 (bottom) lists them with the intercepts. The three class covariances sum to *Y*_*k*_, since 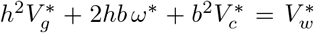 and 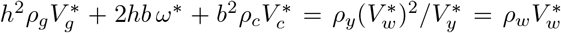by (S25) and (S26), so the intercepts sum to *α*_*k*_. Without assortment *α*_*k*,G_ = *α*_*k*,GC_ = 0 and *α*_*k*_ = *α*_*k*,C_ = *b*^2^*C*_*k*_, which is *b*^2^*ā*_*k*_ under the parental-mean model: the shared environment enters the intercept only. The relation *α*_*k*_ = *Y*_*k*_ −*β*_*k*_*ā*_*k*_ and the class covariances are exact, so the intercepts are approximate only through the slopes, and, for half sibs and their descendants, through *Y*_*k*_. Table S3 collects the results.

**Table S3:** Slopes (top) and intercepts (bottom) of the within-class regression of the standardized cross-product 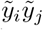on *x* (*V*_*g*_ = *V*_*c*_ = 1). *θ*_*k*_ is given by (S38), *ν*_*k*_ by (S9), and the genetic BLD slope uses 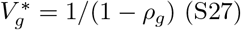. The BLD entries drop the 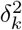term and assume the markers are the causal loci. The IBD entries are exact for siblings and approximate for cousins (§S5.6.1; Figure S7). The class covariances are exact under the model. The BLD entries are approximate (§S5.6.2), and so are the totals and intercepts that use them. Without assortment 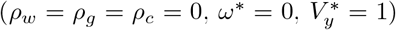 the total slope reduces to *h*^2^, the GC and C slopes to zero, and every intercept to *b*^2^*C*_*k*_ (*b*^2^*ā*_*k*_ under the parental-mean model).

| Component | $\beta^{\text{IBD}}$ | $\beta^{\text{BLD}}$ | $\beta_k = \nu_k \beta^{\text{IBD}} + (1 - \nu_k) \beta^{\text{BLD}}$ |
| --- | --- | --- | --- |
| Genetic (G) | $\frac{h^2}{V_y^*}$ | $\frac{h^2}{(1 - \rho_g)^2 V_y^*}$ | $\frac{h^2}{V_y^*} \left[ \nu_k + \frac{1 - \nu_k}{(1 - \rho_g)^2} \right]$ |
| Gene–environment (GC) | 0 | $\frac{2hb V_g^* \omega^*}{V_y^*}$ | $(1 - \nu_k) \frac{2hb V_g^* \omega^*}{V_y^*}$ |
| Shared environment (C) | 0 | $\frac{b^2 (\omega^*)^2}{V_y^*}$ | $(1 - \nu_k) \frac{b^2 (\omega^*)^2}{V_y^*}$ |
| Total | $\frac{h^2}{V_y^*}$ | $\frac{h^2}{(1 - \rho_w)^2 V_y^*}$ | $\frac{h^2}{V_y^*} \left[ \nu_k + \frac{1 - \nu_k}{(1 - \rho_w)^2} \right]$ |

Table S3: Slopes (top) and intercepts (bottom) of the within-class regression of the standardized cross-product $\tilde{y}_i \tilde{y}_j$ on $x$ ( $V_g = V_c = 1$ ).
| Component | Class covariance | $\alpha_k$ |
| --- | --- | --- |
| Genetic (G) | $h^2 (\bar{a}_k + \theta_k \rho_g) V_g^* / V_y^*$ | $h^2 (\bar{a}_k + \theta_k \rho_g) V_g^* / V_y^* - \beta_{k,G} \bar{a}_k$ |
| Gene–environment (GC) | $2hb (\bar{a}_k + \theta_k) \omega^* / V_y^*$ | $2hb (\bar{a}_k + \theta_k) \omega^* / V_y^* - \beta_{k,GC} \bar{a}_k$ |
| Shared environment (C) | $b^2 (\bar{a}_k + \theta_k \rho_c) V_c^* / V_y^* = b^2 C_k / V_y^*$ | $b^2 C_k / V_y^* - \beta_{k,C} \bar{a}_k$ |
| Total | $Y_k = \frac{V_w^*}{V_y^*} \left( \frac{1 + \rho_w}{2} \right)^{k-1}$ | $Y_k - \beta_k \bar{a}_k$ |

#### S5.6.5 Slope, intercept and phenotypic covariance

With *β*_*k*_, *α*_*k*_ and *Y*_*k*_ as in (S42), the pooled covariance follows from Eq. (1) of the main text, and reduces without assortment to that of the parental-mean model (§S4.1). For cousins the IBD part of *β*_*k*_ understates the slope (§S5.6.1).

Figure S6 compares these class means, slopes and intercepts, component by component, with the simulations, and Figure S7 shows the difference between the two in each class. The class means agree in every class. The slopes agree for siblings, where they sit near the IBD limit. In cousins, at *ρ*_*y*_ = 0.4, the simulated slopes exceed the theory by 0.45 *h*^2^ on average (§S5.6.1), as expected from the in-law terms that the IBD part omits (§S5.6.1). Among unrelated pairs they exceed the BLD limit by 0.20 *h*^2^ (§S5.6.2). The intercepts of cousins lie below the theory, partly because of this slope error. The rest is an offset that is also present among unrelated pairs and grows with *ρ*_*y*_, from −0.0006 under random mating to −0.006 at *ρ*_*y*_ = 0.4. This is consistent with the centring of the simulated phenotypes on the mean over all pairs (§S8.1), whereas the theory takes unrelated pairs as the zero.

In summary, assortative mating affects both the slopes and the intercepts of the within-class lines:

- In close relatives, most of the inflation goes into the intercept. Assortment packages trait-increasing alleles together, and close relatives inherit these packages whatever their realized IBD sharing, so they add a roughly constant amount to the covariance.
- Among unrelated pairs, the inflation comes from small increases in positive LD between trait-increasing variants across the population, and so goes into the slope.
- With a parental-mean MGSE, the genetic and environmental components become coupled through *ω*^∗^: at fixed *h*^2^, increasing *b*^2^ raises the genetic mate correlation *ρ*_*g*_, and vice versa.
- The shared environment acquires a BLD slope, 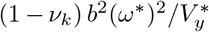 (Table S3), which it does not have under random mating. Assortment and MGSE therefore act together to distort the relationship more than either alone.

## S6 Assortative mating plus imperfect tagging

When both imperfect tagging and assortative mating are occurring, we can again make use of the IBD/BLD split to see how they interact. The IBD term does not depend on which markers are used. The BLD term instead is attenuated by *υ*_*∞*_, approximately, as in (S16). Therefore we can write

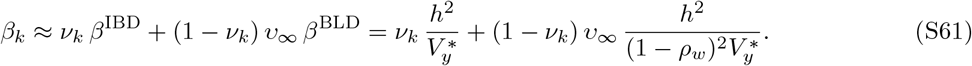

Let Γ be the ratio of the covariance the BLD part contributes per unit of its variance to that of the IBD part,

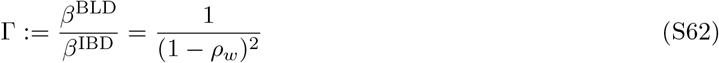

by (S42). For *b* = 0, *ρ*_*w*_ = *ρ*_*g*_ and it is 1*/*(1 − *ρ*_*g*_)^2^, the ratio of the two limits in §S5.6.2; with a parental-mean MGSE component *ρ*_*w*_ *> ρ*_*g*_ and it is larger, because *ω*^∗^ adds to *q*_*g*_ without adding to the genic variance.

Imperfect tagging removes (1 − *ν*_*k*_)(1 *υ*_*∞*_) *β*^BLD^ from the slope, so the fraction of the slope with *M* = *Q* that survives is

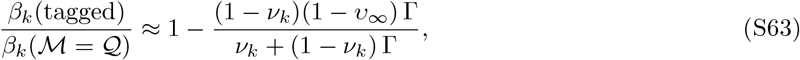

This holds for any *b* under the parental-mean model, since the *b*-dependence sits entirely inside Γ.

In the random-mating case Γ = 1 and (S63) reduces to *υ*_*k*_ = *ν*_*k*_ + (1 *ν*_*k*_)*υ*_*∞*_ (S16). The fraction removed increases with Γ, so as the strength of assortment increases, a larger fraction of the slope is lost to imperfect tagging. This is not an interaction between the two mechanisms within the BLD part, which they scale independently, but a consequence of assortment shifting the slope towards its BLD part, the part that tagging attenuates. The variance shares *ν*_*k*_ are unchanged.

**Figure S6:**
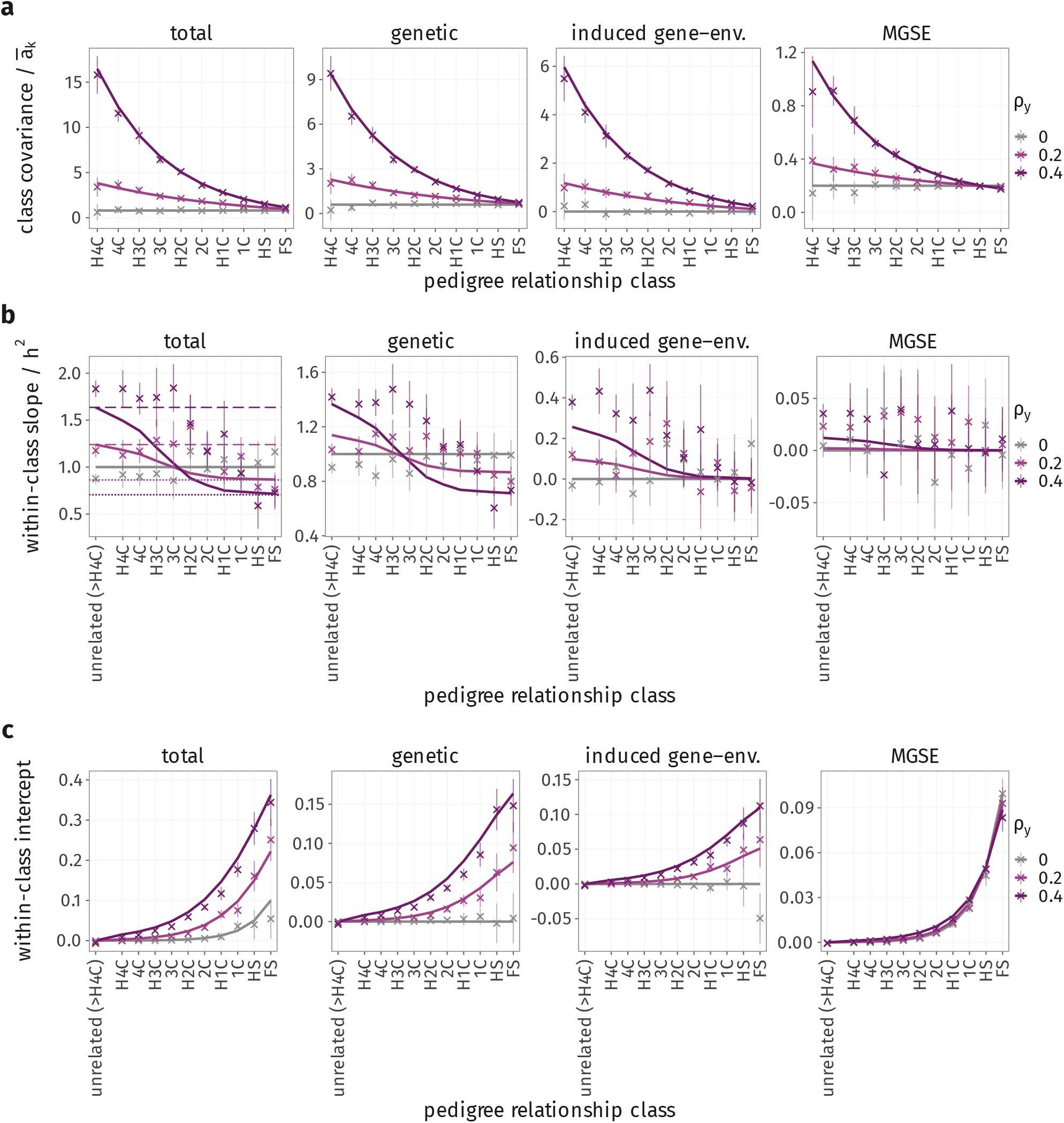
Class covariances, within-class slopes and intercepts under assortative mating, by component. *h*^2^ = 0.6, *b*^2^ = 0.2, *ρ*_*y*_ = 0, 0.2 and 0.4; total, genetic (G), induced gene–environment (GC) and MGSE (C). (a) Class covariance divided by*ā*_*k*_ (Table S3); simulated class means are taken relative to the mean among unrelated pairs. (b) Within-class slope of the cross-product on *x*, relative to *h*^2^, with each simulation’s measured *ν*_*k*_; in the total panel, dotted lines mark the sibling IBD limit 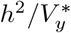 and dashed lines the BLD limit 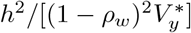]. (c) Within-class intercept. All on the standardized scale (divided by 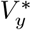). Lines theory; crosses simulation (50 replicates; 95% intervals, ± 1.96 jackknife SE). The theoretical slopes and intercepts of cousins (1C–H4C) are approximate (§S5.6.1; Figure S7).

**Figure S7:**
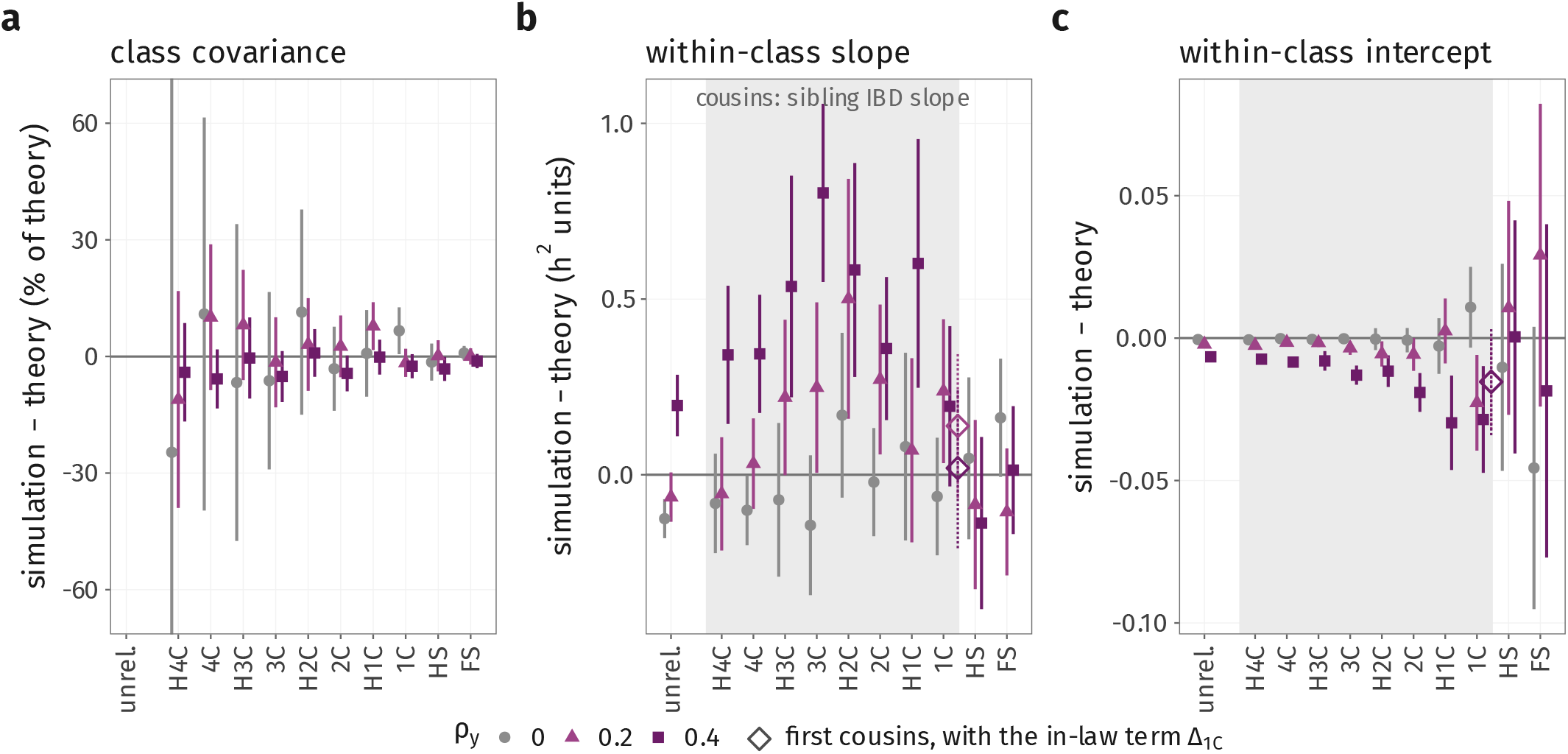
Simulation minus theory in each pedigree class, under assortative mating. The totals of Figure S6 (*h*^2^ = 0.6, *b*^2^ = 0.2, parental-mean MGSE, *ρ*_*y*_ = 0, 0.2 and 0.4). (a) Class covariance, as a percentage of the theory; unrelated pairs have no class covariance on this scale. (b) Within-class slope, in units of *h*^2^. (c) Within-class intercept, on the standardized scale. Points, simulation minus theory; bars, the 95% interval of the simulation (± 1.96 jackknife SE over 50 replicates; the theory has no sampling error). Shaded, the cousin classes (1C–H4C), whose theoretical slope uses the sibling IBD slope 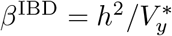and so omits the in-law terms (§S5.6.1). Open diamonds (offset to the right of 1C), first cousins with the in-law term Δ_1C_ added to the theory. Intervals in (a) are clipped.

### Frequency-dependent effect sizes

Under a frequency-dependent model such as (S20), the IBD and BLD parts change differently. The IBD part does not involve the markers’ allele frequencies: by (S46) it is the *u*^2^-weighted average of Cov_*k*_(*R*_*l*_, *R*) over causal loci, against the unweighted average over markers in Var_*k*_(*R*), and Cov_*k*_(*R*_*l*_, *R*) depends on where *l* lies on the map, not on its frequency. So (S47) still holds unless allele frequency, and with it *u*^2^ or marker density, varies systematically along the map. The BLD part is attenuated by 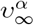, the value of (S19) among unrelated pairs, in place of *υ*_*∞*_. The assortment factor does not change: by (S54) and (S60), *q*_*l*_ = *u*_*l*_ *q*_*g*_, so assortment scales every causal locus by the same factor whatever its frequency. Hence (S61) and (S63) hold with 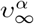 for *υ*_*∞*_. With *α <* −1, rare causal variants get more weight, and if the markers are more common than the causal variants they tag them less well (§S3.1), so 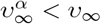 and the attenuation is larger.

### S6.1 Slope, intercept and phenotypic covariance

As with imperfect tagging alone (§S3.4), tagging changes the slope (S61) and leaves the intercept as it is when M = Q, *α*_*k*_ of (S42). The slope falls by (1 − *ν*_*k*_)(1 − *υ*_*∞*_) *β*^BLD^, so the line at *x* = *ā*_*k*_ falls by this amount times *ā*_*k*_, which is small in every class: in close relatives *ν*_*k*_ ≈ 1, and in distant relatives *ā*_*k*_ is small. Under frequency-dependent effect sizes, *υ*_*∞*_ is replaced by 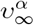.

## S7 Estimating *ν*_*k*_, *M*_*e*_ **and** *υ*_*k*_

The quantities *υ*_*k*_ and *ν*_*k*_ can both be expressed as ratios of second moments of two genomic relationship matrices within a relationship class, Var_*k*_(*a*_*ij*_) and Cov_*k*_(*a*_*ij*_, *g*_*ij*_). They can be estimated in two ways. The first averages over pairs of *individuals*. It needs genotypes at the markers and at the causal variants, and the pedigree class of every pair. This is available from our forward simulation data (§S8), and we use it as a reference. The second averages over pairs of *loci*, as in (S11). It needs only a genetic map and an LD reference panel, and it is what one would use to predict *υ*_*k*_ for real data. Figure S8 compares the two.

As throughout, M is the marker set and Q the causal set (§S2).

### S7.1 Variance in relatedness and IBD

Let **Z** be the *N* × *M* matrix of standardized marker genotypes (S4) in a sample of *N* individuals. The GRM is **A** = **ZZ**^⊤^*/M*, and the sample LD matrix is 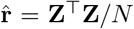. For a symmetric matrix, tr(**U**^2^) is the sum of its squared entries, and tr(**UT**) = tr(**TU**). Hence

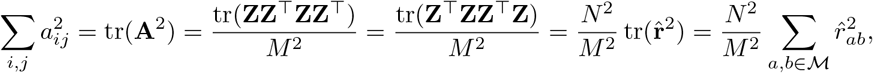

and dividing by *N* ^2^,

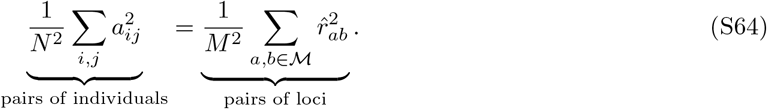

The same holds for **A** and **G**, with 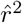 between markers and causal loci. The average LD in a sample is therefore the average squared relatedness of the individuals included in it.

This matters when LD is measured in a sample that contains relatives. Their large *a*_*ij*_ inflate the left-hand side, and so the LD as well, even between loci on different chromosomes. For two unlinked loci, in place of the 1*/N* expected among unrelated individuals,

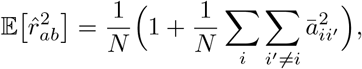

with 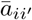 the expected relatedness of *i* and *i*^*′*^. This extra LD is the relatives’ IBD, which the model already counts in *R*. If it were used for Var(*η*) it would be counted twice. So the LD panel should consist of unrelated individuals.

Conversely, if the left-hand side of (S64) is restricted to unrelated pairs, it is the LD of unrelated individuals. The variance of the simulated GRM in class ∞ is therefore the same quantity as Var(*η*) computed from an unrelated panel drawn from the final generation. That is why we use the individual-pair estimates as the reference for the locus-pair ones. Here “unrelated” has a specific meaning: no common ancestor within the five generations used to assign classes (§S8.1). In real data every pair is related at some depth, and “unrelated” means unrelated relative to a threshold or a reference population.

### S7.2 Pairs of individuals

#### Classes

Pairs are assigned to classes *k* = 2, …, 11 as in §S8.1, and pairs with no common ancestor within five generations make up class ∞. Within each class every pair is used, and

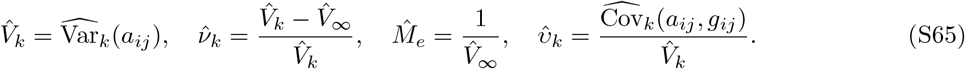

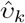 is the least-squares slope of *g*_*ij*_ on *a*_*ij*_ within class *k*, which is (S11) computed directly.

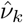 estimates [Var_*k*_(*R*) + Var_*k*_(*η*) − Var_*∞*_(*R*) − Var_*∞*_(*η*)]*/*[Var_*k*_(*R*) + Var_*k*_(*η*)]. This equals *ν*_*k*_ of (S9) under two conditions:

- There is no IBD in the unrelated class. This is an approximation, because “unrelatedness” is a relative rather than absolute term, and always defined with comparison to a baseline sample. However, this can be reconciled with the theory if population LD is measured in the unrelated panel assumed.
- Var(*η*) is the same in every class. This holds approximately: by (S14) and Table S1, Var_*k*_(*η*) departs from its unrelated value only through IBD-sharing states, so the relative error is *O*(*ϕ*_*k*_). It enters 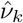 multiplied by 1 − *ν*_*k*_, which is near zero in close relatives, where *ϕ*_*k*_ is large. In distant relatives, where 1 − *ν*_*k*_ approaches one, *ϕ*_*k*_ is small.

### S7.3 Pairs of loci

The locus-pair estimator uses the approximation (S15), *γ*_*k*_ = 1 and *ψ*_*k*_ = 0: each pair of loci contributes its IBD covariance plus its *r*^2^. Averaging over pairs,

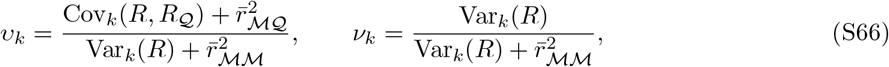

where 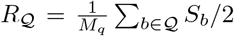 is the IBD part of *g*_ij_, and the averages 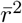 run over all pairs, including each locus paired with itself (§S7.4). This is (S16) without the assumption Cov_*k*_(*R, R*_*Q*_) = Var_*k*_(*R*). It gives 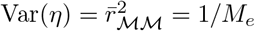 and 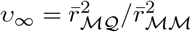, as in (S10).

#### IBD part

By (S12), the IBD covariance of two loci depends only on the recombination fraction between them, so Var_*k*_(*R*) (S14) and

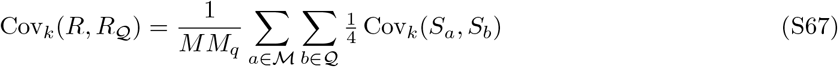

need only the map positions of the loci, not their genotypes. Cov_*k*_(*S*_*a*_, *S*_*b*_) is *ϕ*_*k*_(1 − *ϕ*_*k*_)*τ*_*k*_(*c*_*ab*_) for classes without IBD2 states, 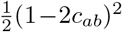 for full sibs, zero for parent–offspring pairs, and zero between chromosomes. Relationship types of the same degree differ in this variance (Hill and Weir, 2011), so a class that pools several relationship types uses the mean over its pairs. We sum over every pair of loci on each chromosome; a random subsample of loci could be used instead.

#### LD part

The LD terms are averages of *r*^2^,

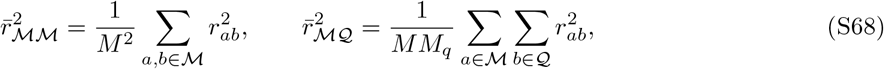

with *r*_*ab*_ the correlation of genotype dosages in the panel, as in the GRM. There are too many pairs to enumerate. We draw a random sample of markers, compute the *r*^2^ of each with every other locus on its chromosome, and scale the sum up to all markers. Pairs on different chromosomes, which are nearly all unlinked, are averaged over a smaller random subsample of loci. Sampling noise in 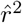 is removed with 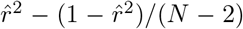 (Bulik-Sullivan et al., 2015), which is valid for a panel of *N* unrelated individuals (§S7.1).

### S7.4 Same-locus terms

The averages (S68) include each locus paired with itself, with 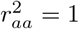. These pairs are added exactly rather than estimated with the others:

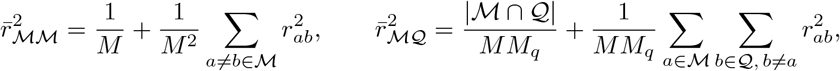

where |M ∩ Q| is the number of loci that are both markers and causal. For a pair with no IBD, each same-locus term *z*_*il*_*z*_*jl*_ has variance 1, which gives the 1*/M* in the first. The share of the term 1*/M* in Var(*η*) is *M*_*e*_*/M*. That is small, but not negligible at typical marker densities.

### S7.5 Frequency-dependent effect sizes

Under the random-effects model of §S3.3, causal locus *b* carries the relative effect-size variance *ζ*_*b*_ of (S18); under the *α*-model (S20), *ζ*_*b*_ ∝ [2*f*_*b*_(1 − *f*_*b*_)]^1+*α*^, and *ζ*_*b*_ = 1 for every locus when *α* = −1. The causal relatedness is then 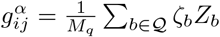, the entries of **G**_*α*_, and 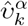 is (S65) with *g*^*α*^ in place of *g*.

In the locus-pair estimator, every sum over causal loci *b* carries the weight *ζ*_*b*_: in Cov_*k*_(*R, R*_*Q*_) (S67), in 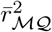 (S68), and in the same-locus term, which becomes ∑_*b*∈*M*∩*Q*_ *ζ*_*b*_/(*MM*_*q*_). Every partner of a sampled marker is included in the LD sums, so the weights enter exactly, and one sample of markers serves every *α*.

When every variant is causal, **G**_*α*_ is a weighted sum of matrices built separately for each allele-frequency bin B. With *M*_*B*_ loci in the bin, *g*_*B,ij*_ the relatedness at those loci, and *ζ*_*B*_ the weight at the bin’s mean frequency, 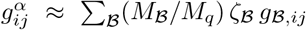. The class covariance is linear in *g*, so 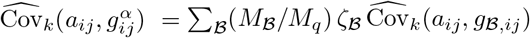, and 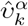 for every *α* follows from the per-bin covariances, exact up to the variation of 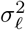 within a bin.

## S8 Supplementary Methods

### S8.1 Forward-in-time simulations with xftsim

We conducted forward-in-time simulations using the `xftsim` package (Border et al., 2026).

#### Genetic data and recombination

We generated genotypes for 5,000 founder individuals representing the GBR population of the 1000 Genomes Project (Phase 3; The 1000 Genomes Project Consortium, 2015) using `HAPGEN2` (Su et al., 2011). Recombination was simulated explicitly using the sex-averaged human Pyrho recombination map (Spence and Song, 2019).

#### Demography

Generations were discrete and non-overlapping. At each generation, females and males were paired at random (or assortatively: each sex’s phenotypes were standardized and mixed with independent Gaussian noise, 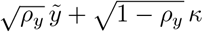, and females and males were paired by rank on this score, which gives a phenotypic correlation between mates of *ρ*_*y*_; the linear assortative mating regime of `xftsim`). When the two sexes were unequal in number, the smaller pool was resampled with replacement, so that some individuals mated more than once and the pedigree contains half-sibling relationships. Each pair produced a Poisson-distributed number of offspring, with the mean reset each generation to *N/n*_pairs_ so that the population stayed at the founder size *N*. We recorded all parent–offspring relationships, giving the full pedigree at the end of the simulation.

**Figure S8:**
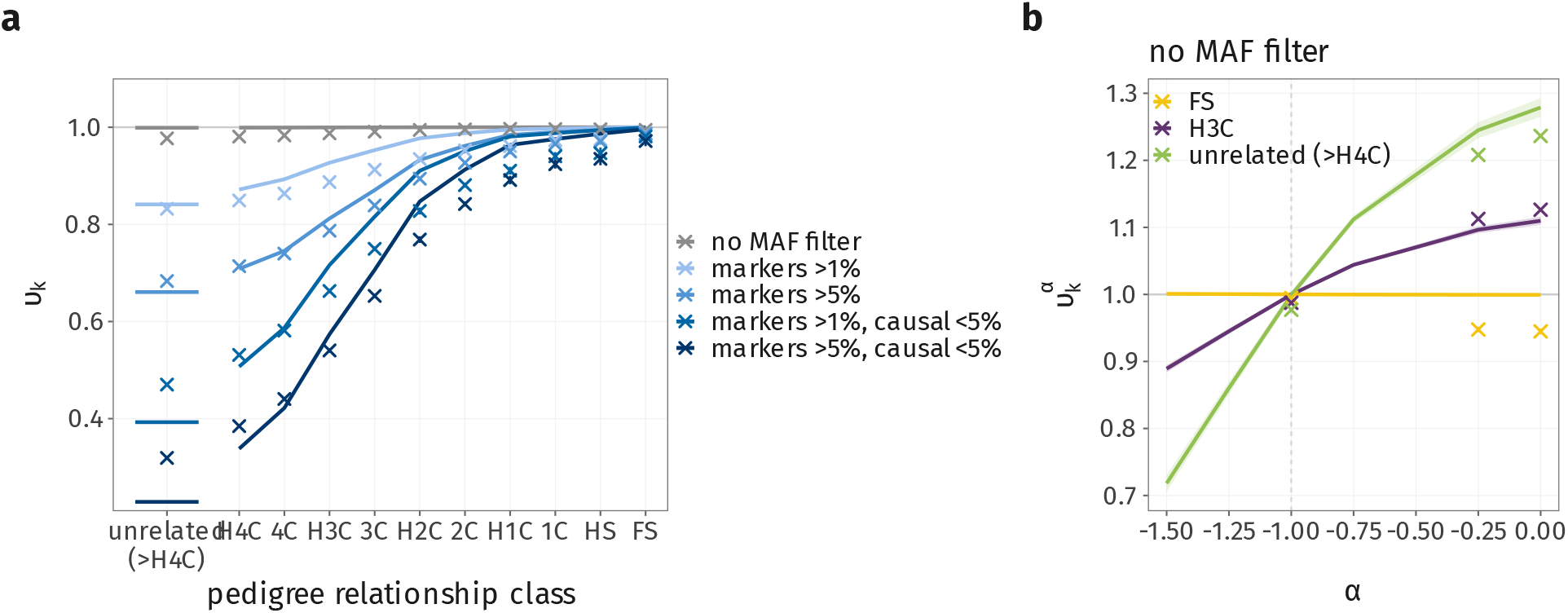
Tagging *υ*_*k*_ **estimated from pairs of individuals and from pairs of loci**. (a) *υ*_*k*_ by pedigree class for each marker/causal MAF scenario. Pairs of individuals: the within-class slope of the causal on the marker GRM (crosses, mean ± SD across replicates). Pairs of loci: the locus-pair estimator (S66) on the founders panel with the same marker and causal sets (lines, mean over 3 replicates; computed to half fourth cousins (H4C), *k* = 11, and for *k* =∞). (b) 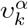 against *α* for full sibs, half third cousins and unrelated pairs, with every variant both marker and causal: pairs of loci (lines, ± SD band) and pairs of individuals where simulated (crosses).

#### Tracking pedigree relatedness

We computed the pedigree relatedness *ā*_*ij*_ (§S4), twice the kinship coefficient, among individuals in the final generation using the standard recursion, carried forward one generation at a time,

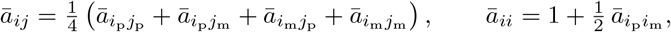

where *i*_p_ and *i*_m_ are the father and mother of *i*. We checked it against the *pedigree kinship* function of `sgkit` (https://sgkit-dev.github.io), and the two agreed exactly. We computed *ā*_*ij*_ both over the full pedigree and over the most recent five generations only; for the latter, individuals five generations back are treated as unrelated, non-inbred founders.

Over many generations, pairs of the same nominal degree differ slightly in their realized pedigree relationship, because of loops in the pedigree above their most recent common ancestors. To obtain well-defined classes, we assign classes using the five-generation relationship. All sampled individuals belong to the same generation, so a pair related through a single common ancestor, or a single ancestral couple, within this window has relationship exactly 2^−(*k*−1)^ for an integer *k* (§S1), which we take as its class. This gives ten classes, from full siblings (*k* = 2) and half siblings (*k* = 3) through first cousins (*k* = 4) to half fourth cousins (*k* = 11). Pairs related through more than one ancestor or couple have relationships that are not powers of two and are excluded: about 2% of all pairs, but 9% of pairs related within five generations. These pairs have higher relatedness than single-path pairs of the same degree, so the exclusion trims the right tails of the class distributions *f*_*k*_(*x*). Pairs with no common ancestor within five generations form class ∞ (§S7.2).

#### Genomic relatedness

At the end of each simulation, we computed the genomic relatedness between individuals from their genotypes at a specified set of SNPs, as the mean product of their standardized genotypes (S4).

#### Parental-mean environmental transmission

We simulated environmental transmission forward in time under the parental-mean model (S2). At the start of the simulation, each founder was assigned an environmental value *c*_*i*_ ~ N (0, 1). In each subsequent generation, each offspring inherited the mean of its parents’ values plus an independent Gaussian deviation,

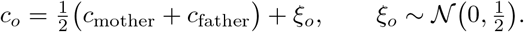

With this choice of Var(*ξ*), the variance of *c* stays constant across generations under random mating (§S4.1). Under random mating, and because the founders’ values are independent, E[*c*_*i*_*c*_*j*_ | *P*] is the pairs’ pedigree relatedness over all 20 generations, so pairs in class ∞ still share part of their environment (*C*_*∞*_ = 0.004). As in the data, the simulated phenotypes are centred, so their covariances are measured relative to the average pair (Section 2.4 of the main text).

#### Phenotype

Phenotypes were generated under the additive model with a shared-environment component (S1), *y*_*i*_ = *h g*_*i*_ + *b c*_*i*_ + *e ϵ*_*i*_. Here *g*_*i*_ = ∑_*l*∈*Q*_ *u*_*l*_*z*_*il*_ is the breeding value at *M*_*q*_ = 100,000 causal SNPs, with effects *u*_*l*_ ~ N (0, 1*/M*_*q*_), and *ϵ*_*i*_ ~ N (0, 1), so that the genetic, environmental and residual parts have variances *h*^2^, *b*^2^ and *e*^2^ = 1 − *h*^2^ − *b*^2^.

#### Simulation length and outputs

Simulations were run for 20 generations. After the last generation, all individuals were sampled, and for every pair we computed

1. the genomic relatedness at the marker SNPs, 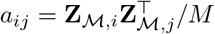, with **Z**_*M*_ the matrix of standardized genotypes at the *M* markers (S4);
2. the pedigree relatedness *ā*_*ij*_, from the paths connecting the pair through the pedigree; and
3. the covariance of the genetic component, both at the variant level, 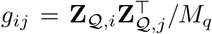, and for the realized breeding values, *g*_*i*_*g*_*j*_; of the MGSE component, *c*_*i*_*c*_*j*_; and of the phenotype, *y*_*i*_*y*_*j*_.

We ran 50 replicates of each scenario. Throughout, 95% intervals are ± 1.96 delete-one-replicate jackknife standard errors across replicates.

#### S8.1.1 Imperfect tagging in forward-in-time simulations

We also examined imperfect tagging in these simulations, with the classes *k* assigned from the pedigree as above. We computed the causal relatedness at a set of causal loci with allele frequency below *f*_0_, **G**^(*<f*0^), following Yang et al. (2010), and the genomic relatedness at marker loci with frequency above 1%, **A**^(*>*1%)^. The slope of the regression of 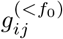 on 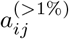 within each class *k* estimates *υ*_*k*_ (S11); we write it 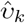. Figure 3 of the main text shows the results for *f*_0_ = 5%.

We also computed the expected 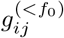for all pairs with 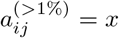 under the sampling schemes of the main text, which vary the proportion of pairs that are close relatives rather than unrelated. For a given scheme, the classes present at *x* have proportions *π*_*k*_(*x*) (§S1), and by (S21)

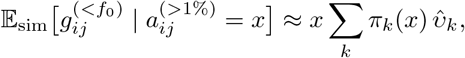

and the expected phenotypic covariance is *h*^2^ times this.

## S9 Empirical methods

### S9.1 Cohorts and sample selection

We analysed the UK Biobank (Bycroft et al., 2018), filtered by similarity to CEU / GBR. Specifically, we used two rounds of projection and filtering, adapted from Kemper et al. (2021). Projection used HapMap3 SNPs (The International HapMap 3 Consortium, 2010) shared with 1000 Genomes (MAF ≥ 1% in the reference), hard-called from the imputed data (INFO ≥ 0.3, minor allele count ≥ 5, missingness ≤ 5%, biallelic). In the first round, we computed two principal components from all 1000 Genomes samples, projected UK Biobank participants using the reference allele frequencies, and kept those within the 99% ellipse (Mahalanobis distance, 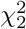) around the 1000 Genomes European (EUR) samples in PCs 1–2. We then re-filtered SNPs within this subset (MAF ≥ 1%, Hardy–Weinberg equilibrium (HWE) *p ≥* 10^−5^) and repeated the projection on principal components of the 1000 Genomes EUR samples. Self-reported ethnicity was not used, and participants who withdrew consent were excluded. In the second round, we kept individuals whose distance to the 1000 Genomes CEU and GBR samples gave *p* ≥ 10^−5^. This left *N* ≈ 430,000 participants.

### S9.2 Genomic relatedness and pair selection

In this preliminary analysis, we computed relatedness in two stages, to avoid computing it for all pairs of individuals. First, we computed a “low-pass” GRM on a random set of 50,000 SNPs and screened all pairs for *a*_*ij*_ *>* 0.03. Second, we added 100,000 random pairs, drawn among the individuals in the screened pairs, to represent the unrelated end of the spectrum. We then recomputed *a*_*ij*_ for every selected pair on the full SNP set (HapMap3 SNPs with MAF ≥ 1%, HWE *p ≥* 10^−6^ and missingness ≤ 5%, without LD pruning; approximately 10^6^) with the estimator of Yang et al. (2010), using the SNPs genotyped in both individuals. As the screen used a noisy estimate, many screened pairs fall below 0.03 on recomputation, so pairs with *a*_*ij*_ *<* 0.02 are a mixture of screened and random pairs; this produces the two modes of relatedness near 0.03.

We did not separate parent–offspring pairs from full siblings in this first analysis; the full-sibling range therefore includes parent–offspring pairs.

### S9.3 Phenotypes and covariates

We analyzed the traits shown in Figures S9–S16. These are continuous traits and biomarkers with more than 100,000 non-missing values, at the baseline assessment; values were used as recorded, without transformation. Each trait was residualized by ordinary least squares, adjusting for sex, age at recruitment, year of birth and genotyping batch. We then removed observations with residuals beyond 5 SD, to exclude implausible values, and standardized the remainder within sex. We did not include genomic principal components in this preliminary analysis.

### S9.4 Phenotypic covariance across relatedness

For each selected pair we computed the cross-product of the two individuals’ standardized residuals, 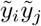, whose expectation is the phenotypic covariance of the pair. For display, we averaged cross-products in bins of relatedness of width 0.005 and plotted bins containing more than 50 pairs.

### S9.5 Regressions across relatedness ranges

We fitted ordinary least squares regressions of the pair-level cross-products on *a*_*ij*_, not of the bin means. The figures show a regression with an intercept within the full-sibling range (0.4 ≤ *a*_*ij*_ ≤ 0.6). We also fitted, but do not show, the same regression within the half-sibling range (0.2 ≤ *a*_*ij*_ ≤ 0.3) and a regression over 0.05 ≤ *a*_*ij*_ ≤ 0.7 with a single slope, a reference intercept, and offsets for 0.1 *< a*_*ij*_ ≤ 0.2, 0.2 *< a*_*ij*_ ≤ 0.4 and 0.4 *< a*_*ij*_ ≤ 0.6. The reference intercept applies to the pairs in neither band, 0.05 ≤ *a*_*ij*_ ≤ 0.1 and 0.6 *< a*_*ij*_ ≤ 0.7, so it pools distant relatives with the first-degree relatives whose relatedness exceeds 0.6. In each model the heritability estimate is the coefficient on *a*_*ij*_. We estimated uncertainty with 100 bootstrap resamples of pairs; because individuals appear in many pairs, these intervals are likely too narrow, and they are not shown.

### S9.6 SNP heritability reference

The dashed reference slope in Figure 8 and the supplementary figures is the SNP heritability estimated by LD score regression in unrelated individuals (Bulik-Sullivan et al., 2015), from the Neale Lab UK Biobank round 2 release (Neale Lab, 2018; Walters and Palmer, 2022): both sexes, 361,194 unrelated white-British participants, rank-inverse-normal transformed traits where available. It is a reference only: it comes from a different design and is not the expected slope under any of the models in the main text. Traits without a Neale Lab estimate have no dashed line.

### S9.7 All of Us analysis

We applied the same approach to the All of Us Research Program (The All of Us Research Program Investigators, 2019; The All of Us Research Program Genomics Investigators, 2024) (CDR v9, short-read WGS). The main differences from the UK Biobank pipeline are as follows.

#### Ancestry filtering

In the first round, we gated participants on the first five of the precomputed All of Us ancestry principal components (PCs). We kept those within the Mahalanobis radius that contains 99% of the 1000 Genomes European founders. In the second round, we refit a PCA within this set. We used HapMap3 variants (MAF *>* 1%, HWE *p >* 10^−6^, missingness *<* 5%), excluded long-range LD regions (Price et al., 2008) and pruned to *r*^2^ *<* 0.05. We then projected 1000 Genomes Europeans into this space. We kept participants within the 99.9% quantile of the projected CEU and GBR samples’ distances on PCs 1–5. This left 227,561 participants.

#### Genomic relatedness

We built the GRM from around 1 million biallelic variants with MAF *>* 1%, HWE *p >* 10^−6^ and missingness *<* 5%. These variants were taken from the allele count/allele frequency (ACAF) threshold subset, intersected with HapMap3, plus a random 20% of the non-HapMap3 ACAF variants. We binned all pairs, without subsampling. Parent–offspring and full-sibling pairs have the same expected relatedness. To separate them, we selected pairs with *a*_*ij*_ *>* 0.2 and classified them with KING (Manichaikul et al., 2010). Parent–offspring pairs are shown separately, and full-sibling estimates exclude them. Uncertainty comes from a 50-block delete-block jackknife over individuals.

#### Phenotypes and covariates

Traits come from physical measurements, electronic health record laboratory values and surveys, defined by Observational Medical Outcomes Partnership (OMOP) concept IDs. We used the most recent value per participant. We removed implausible values, then applied a rank-based inverse-normal transform. We residualized on age and the first 5 genomic PCs. These were computed from HapMap3 common variants pruned at *r*^2^ *<* 0.1 within the analysis sample. We removed residuals beyond 5 SD and standardized within sex.

#### Regressions

The unrelated slope is fitted to pairs with *a*_*ij*_ *<* 0.02. The half-sibling and full-sibling ranges match the UK Biobank analysis. The OneSlopeOffsets model uses only pairs with *a*_*ij*_ ≥ 0.09, and fits a single slope with a separate intercept in each of three bands, *a*_*ij*_ ∈ [0.09, 0.18), [0.18, 0.36) and [0.36, 0.7). As for the UK Biobank, the figures show only the full-sibling fit.

## S10 Empirical covariance profiles in the UK Biobank

Figures S9–S16 show the traits not in Figure 8, grouped by trait type.

## S11 Empirical covariance profiles in All of Us

Figures S17–S22 show the same profiles in All of Us for 36 traits, grouped by trait type. The encoding follows the UK Biobank figures, with three differences: each bin mean carries a ±1 standard error, parent–offspring pairs are binned separately, and the dashed slope is 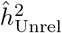 estimated in the same data rather than an external LD score regression estimate.

**Figure S9:**
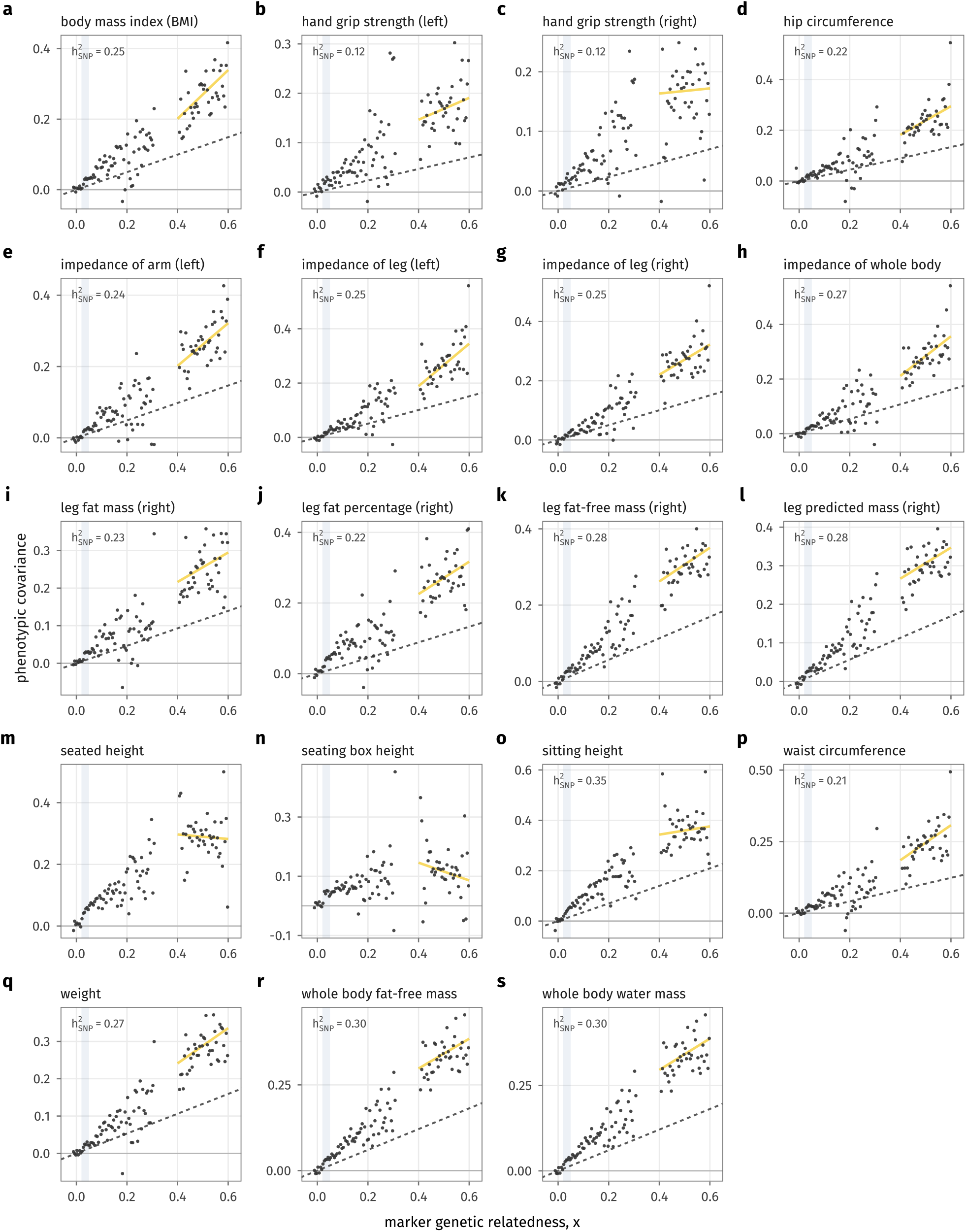
As Figure 8, for anthropometric traits. Panels without a dashed line have no Neale Lab estimate.

**Figure S10:**
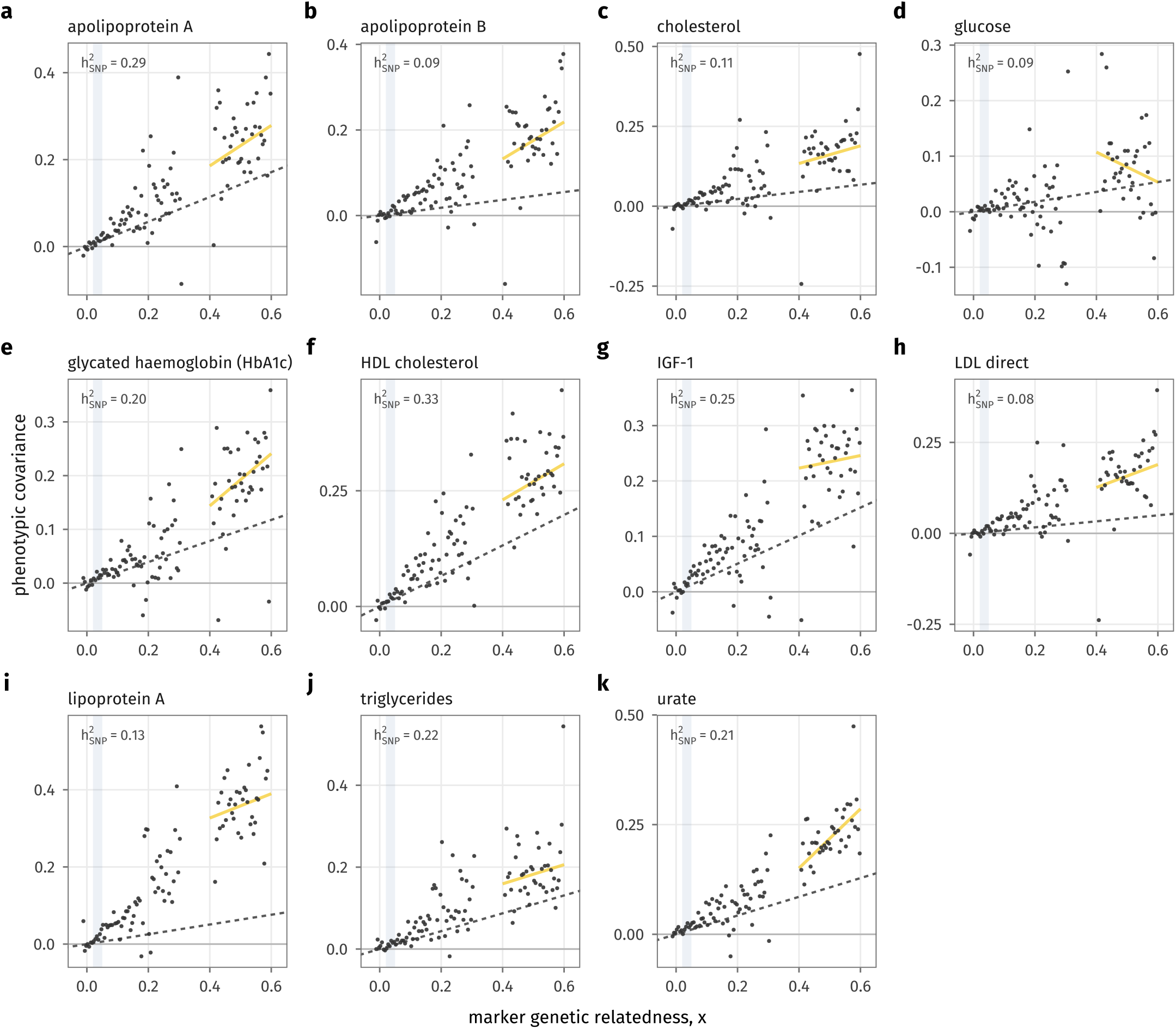
As Figure 8, for metabolic traits. Panels without a dashed line have no Neale Lab estimate.

**Figure S11:**
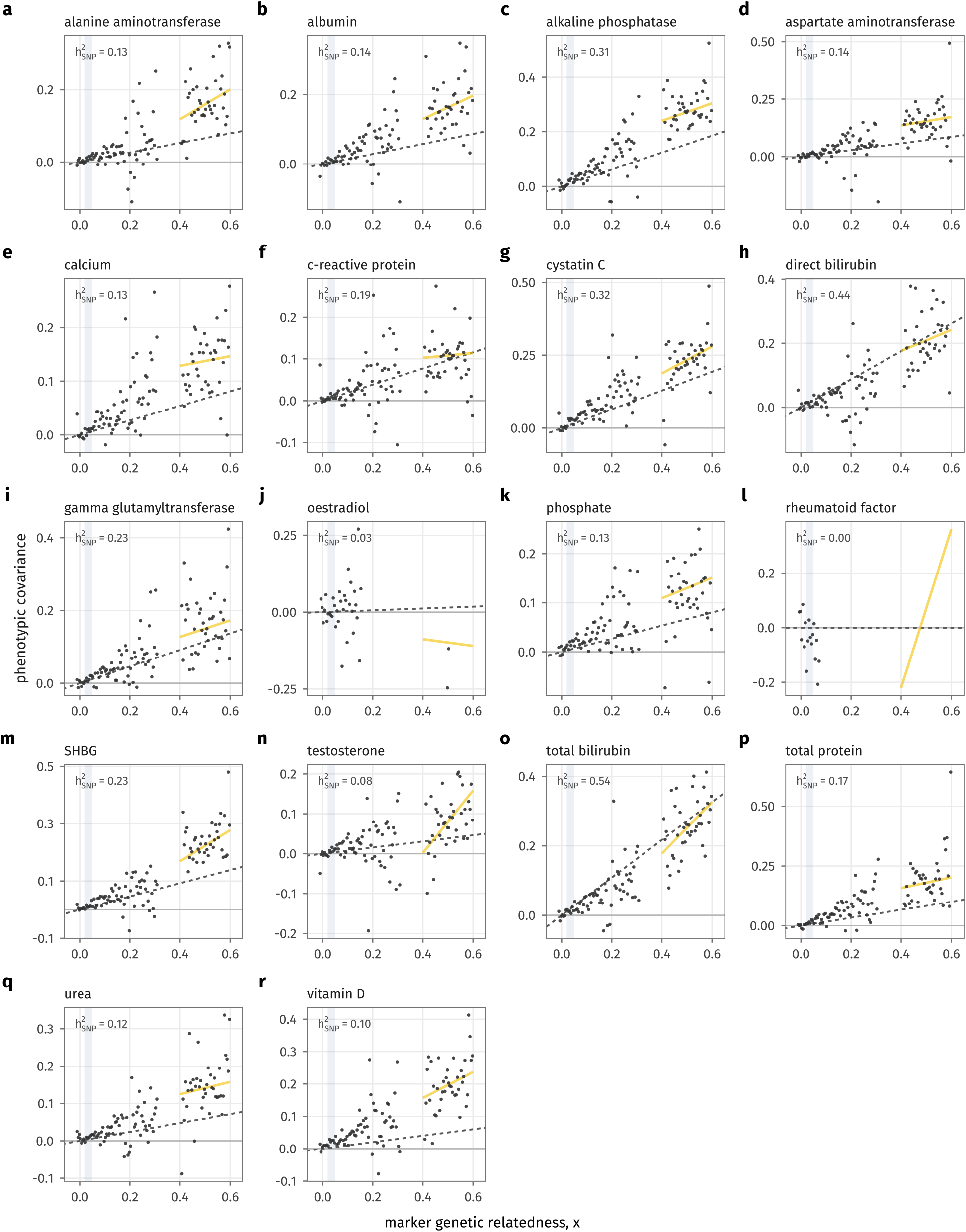
As Figure 8, for other blood biomarker traits. Panels without a dashed line have no Neale Lab estimate.

**Figure S12:**
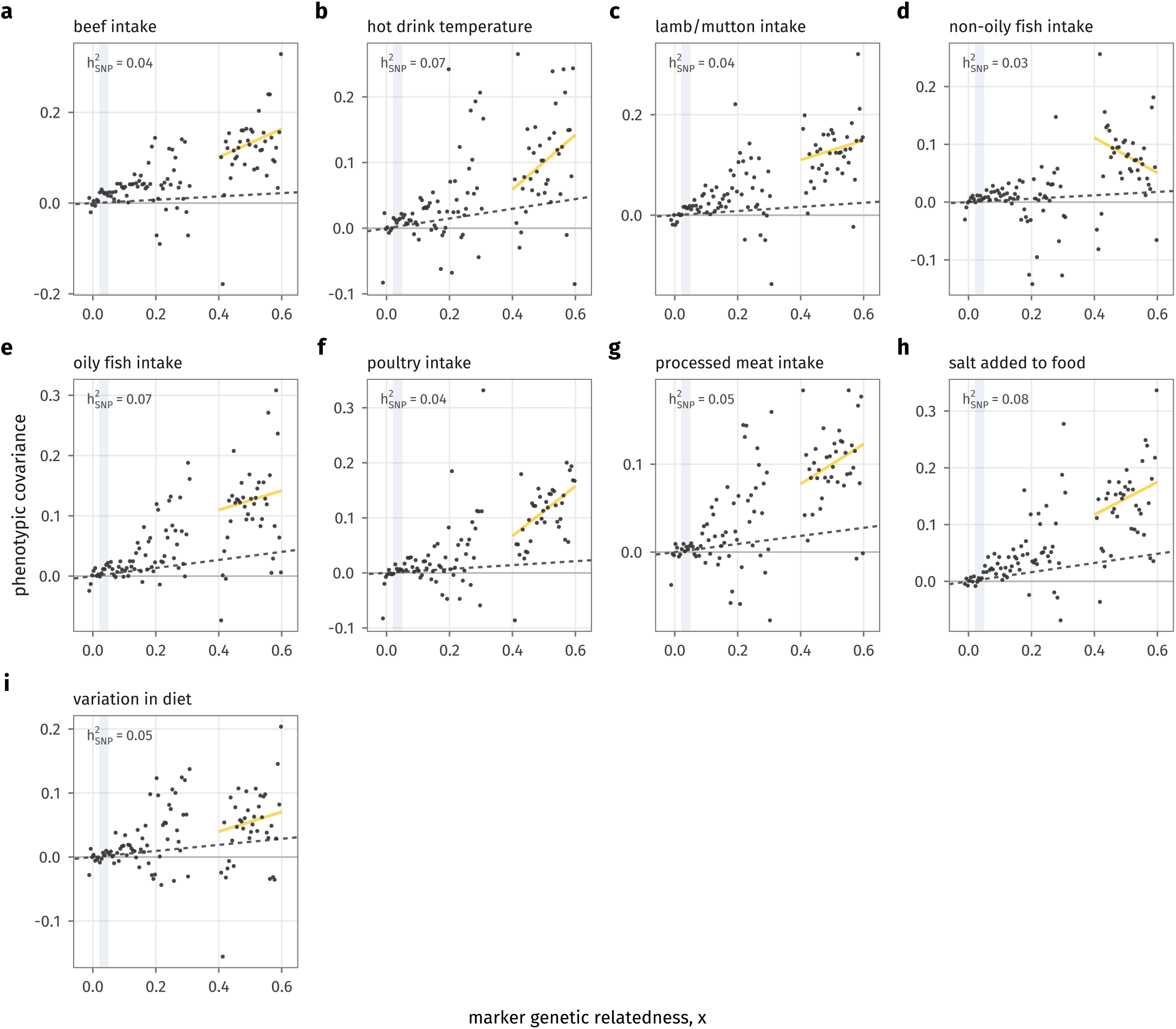
As Figure 8, for dietary traits. Panels without a dashed line have no Neale Lab estimate.

**Figure S13:**
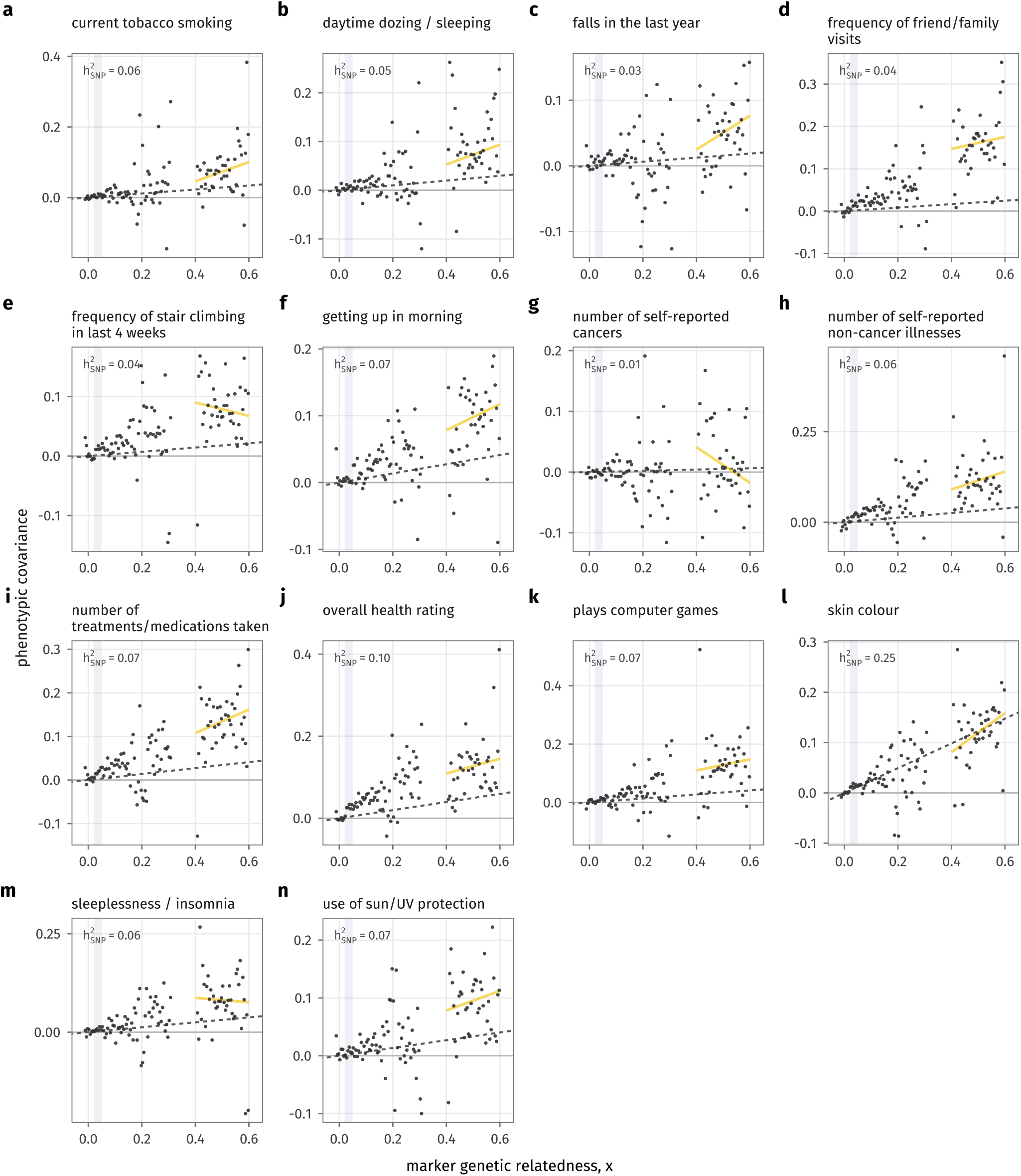
As Figure 8, for lifestyle and health traits.

**Figure S14:**
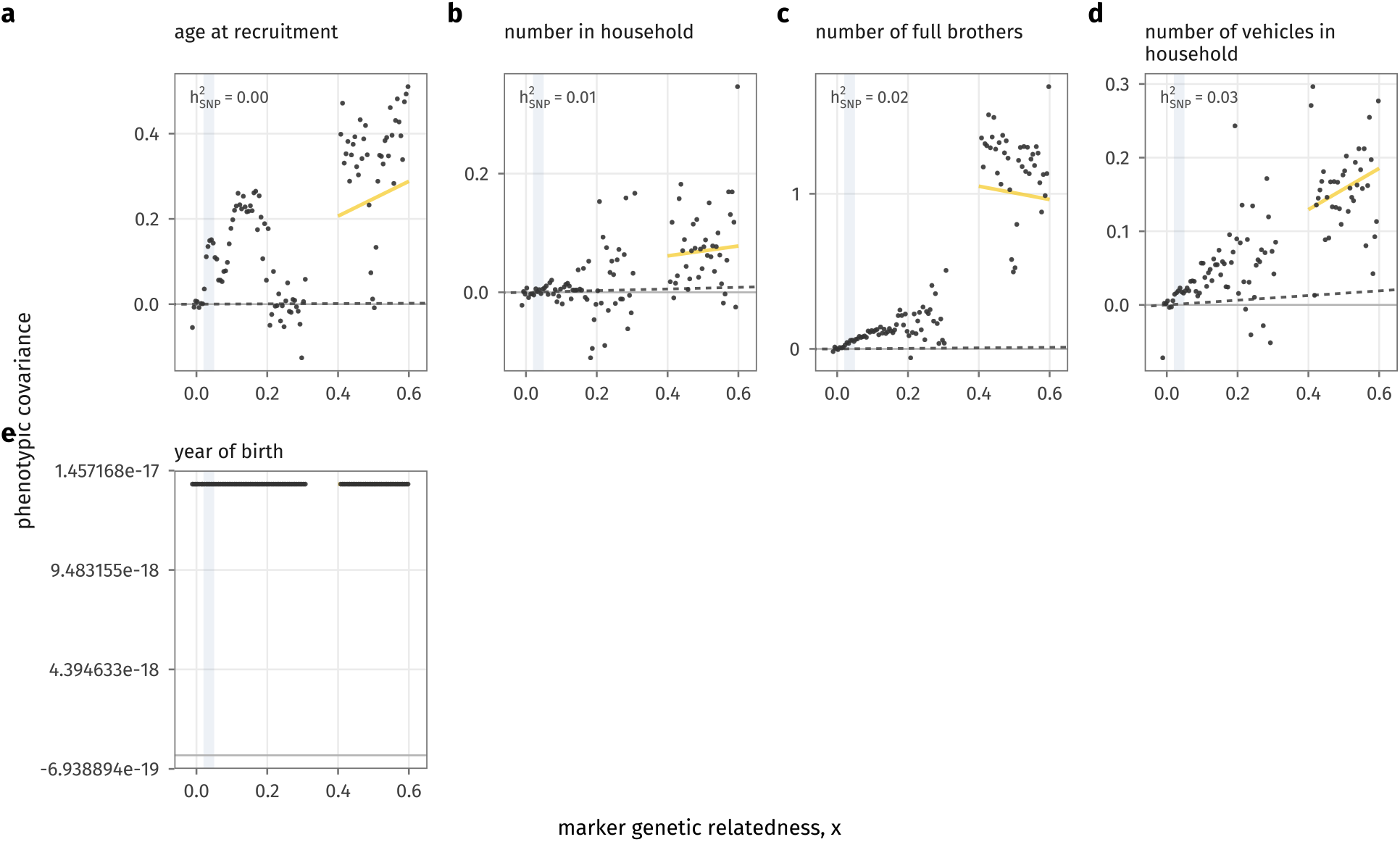
As Figure 8, for household and demographic traits.

**Figure S15:**
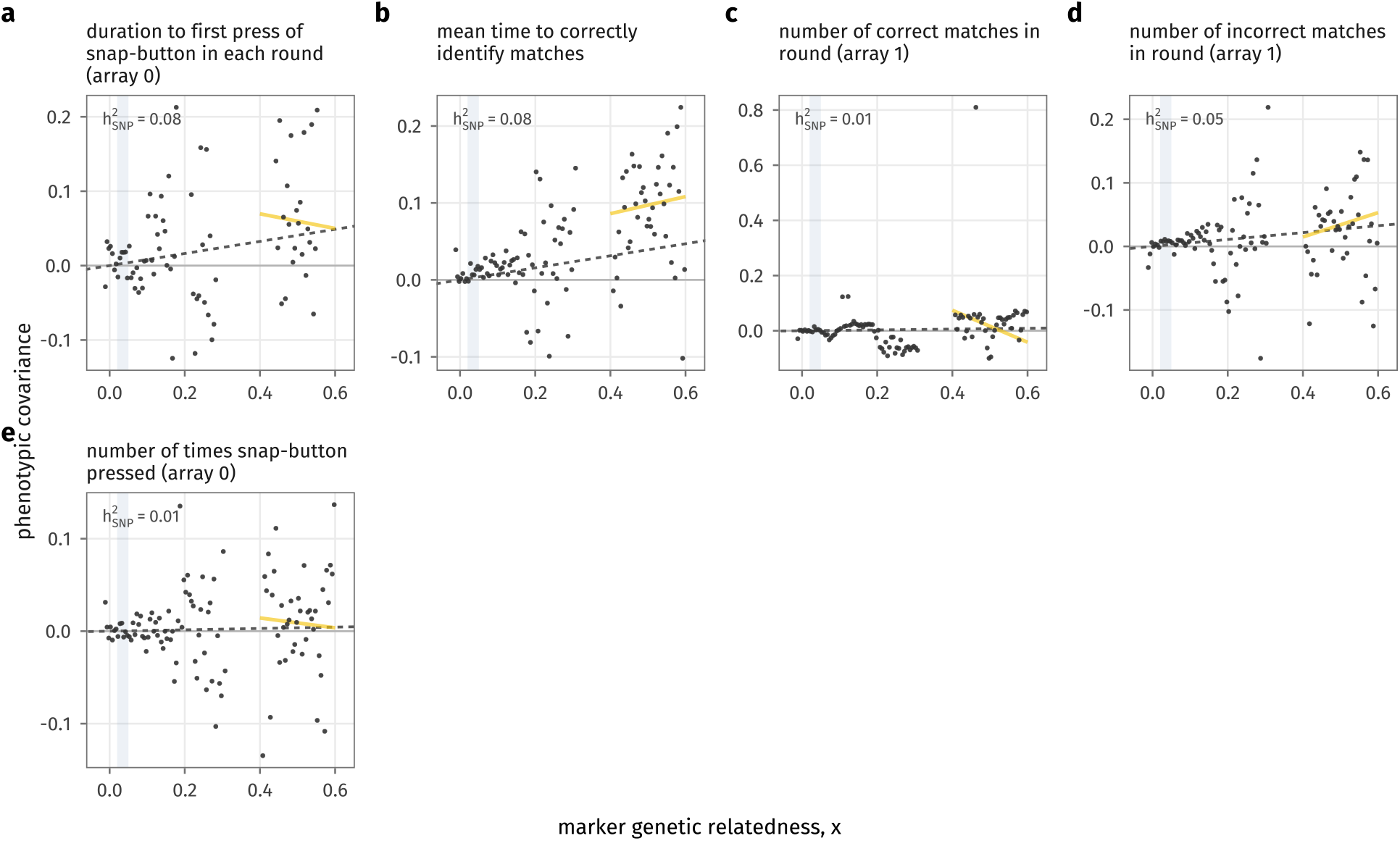
As Figure 8, for cognitive function traits.

**Figure S16:**
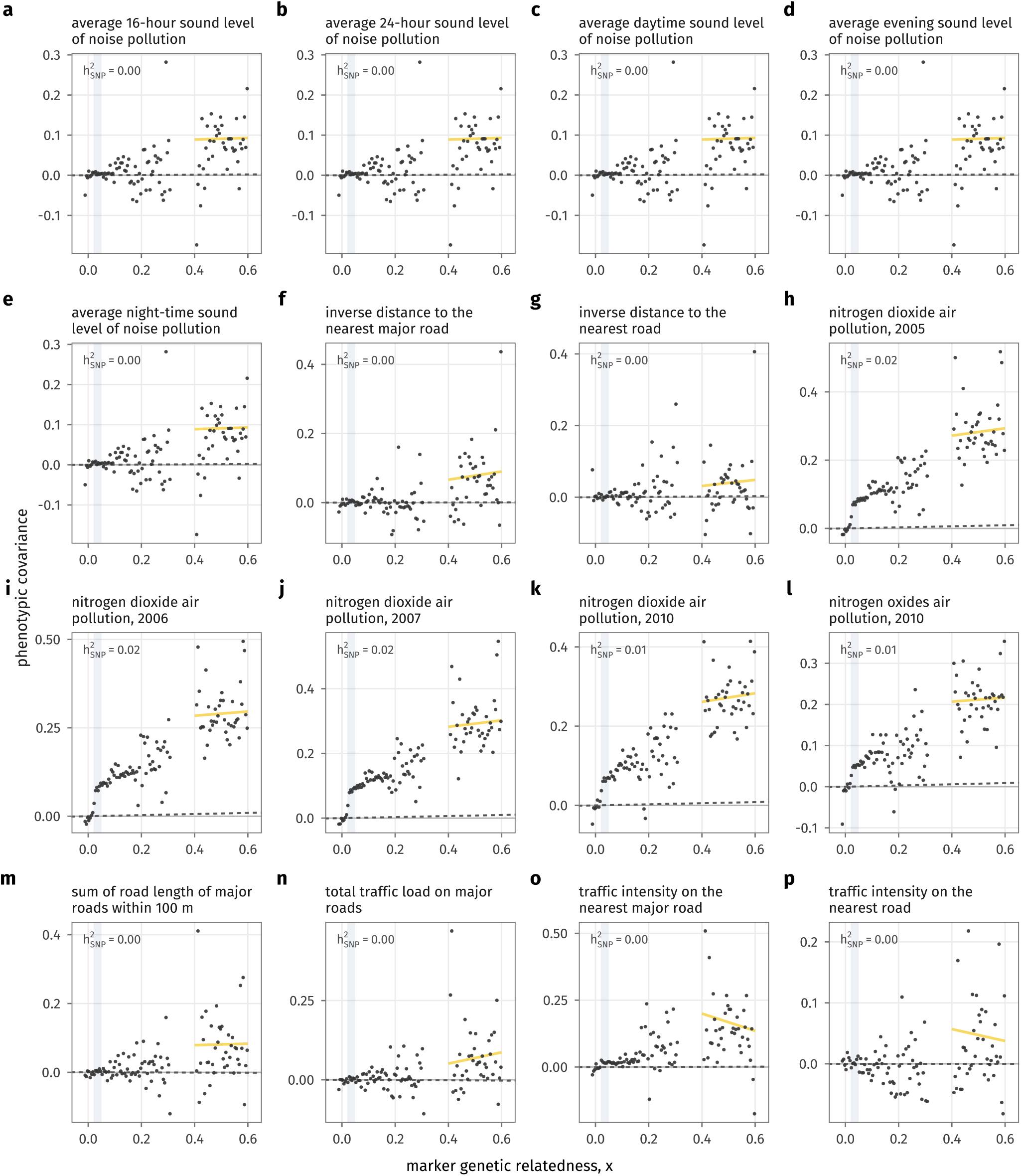
As Figure 8, for environmental exposure traits.

**Figure S17:**
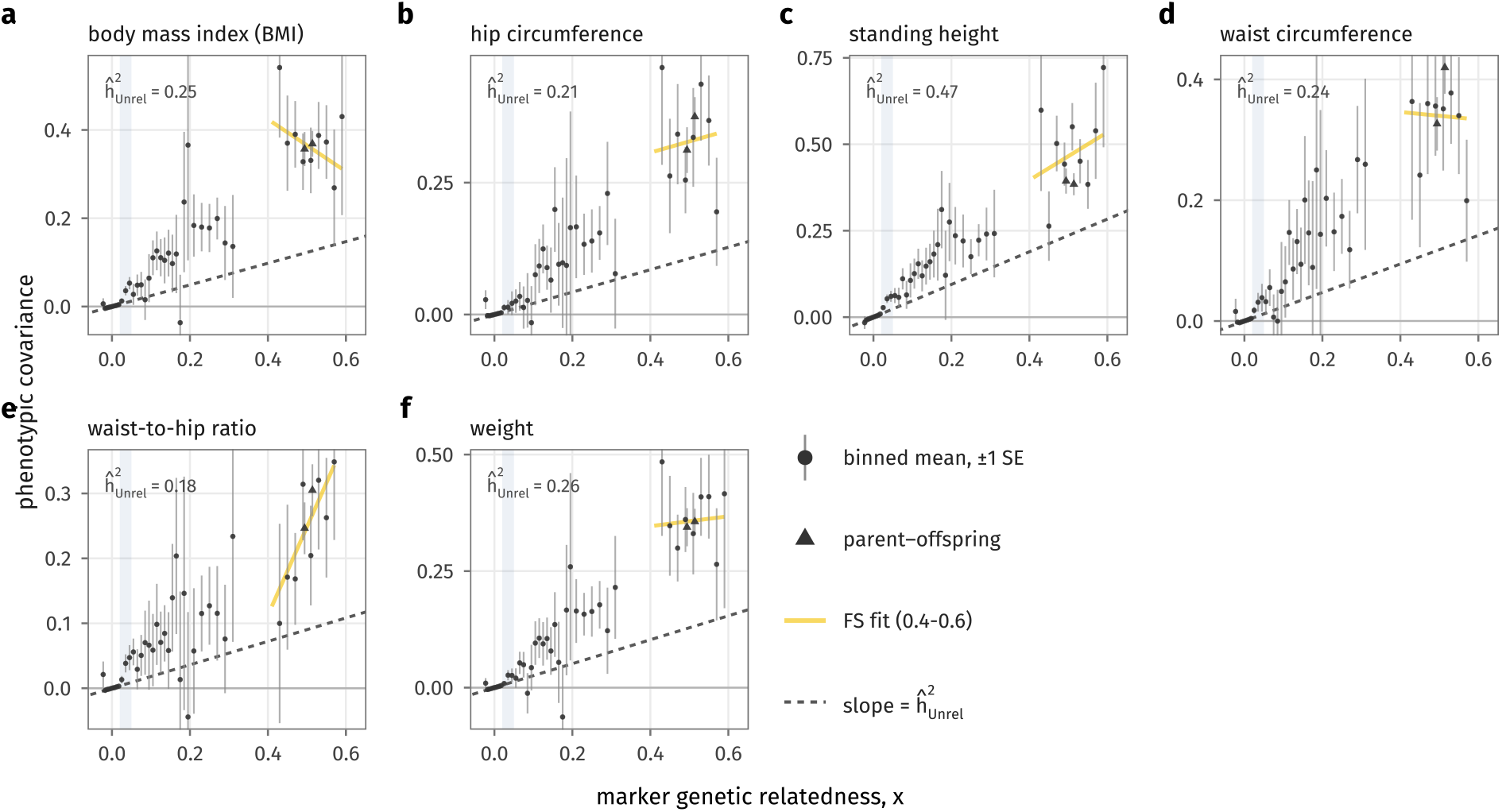
Phenotypic covariance across genetic relatedness in All of Us: anthropometric traits. Phenotypic covariance, estimated as the mean cross-product of standardized residuals, 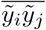, in bins of genetic relatedness *x* (width 0.005 below *x* = 0.02, 0.01 to *x* = 0.2 and 0.02 above; bins with at least 50 pairs), ± 1 standard error from a delete-block jackknife over individuals. Circles: all pairs except parent–offspring; triangles: parent–offspring pairs, binned separately and excluded from all fits. Gold line: regression with an intercept within the full-sibling range (0.4 ≤ *x* ≤ 0.6). Dashed lines: slope equal to 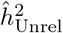, the regression through the origin on pairs with *x <* 0.02 (value in each panel). Shaded: *x* ∈ [0.02, 0.05]. Rank-inverse-normal transformed residuals adjusted for age, sex and genetic principal components.

**Figure S18:**
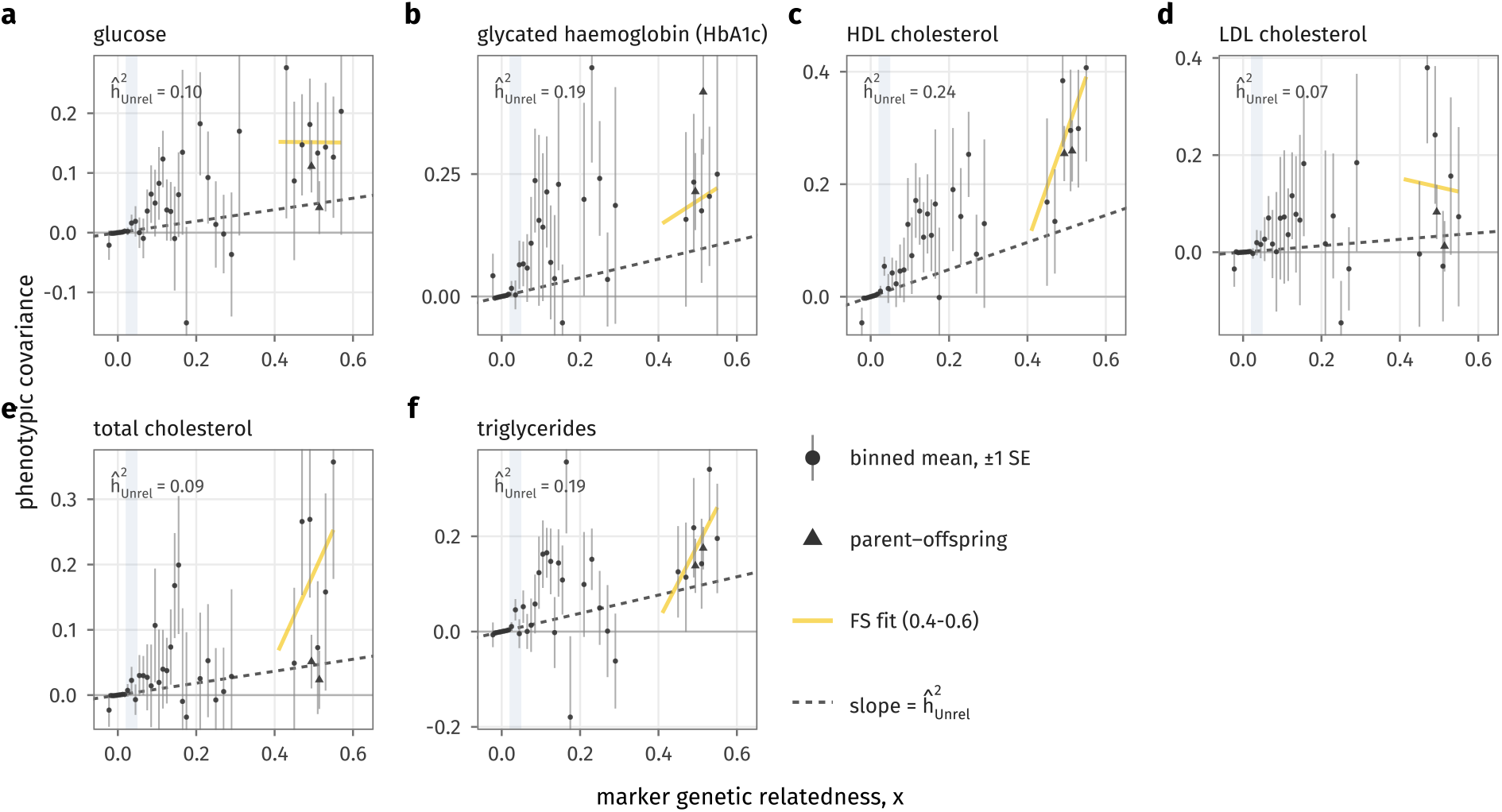
As Figure S17, for lipids and glucose.

**Figure S19:**
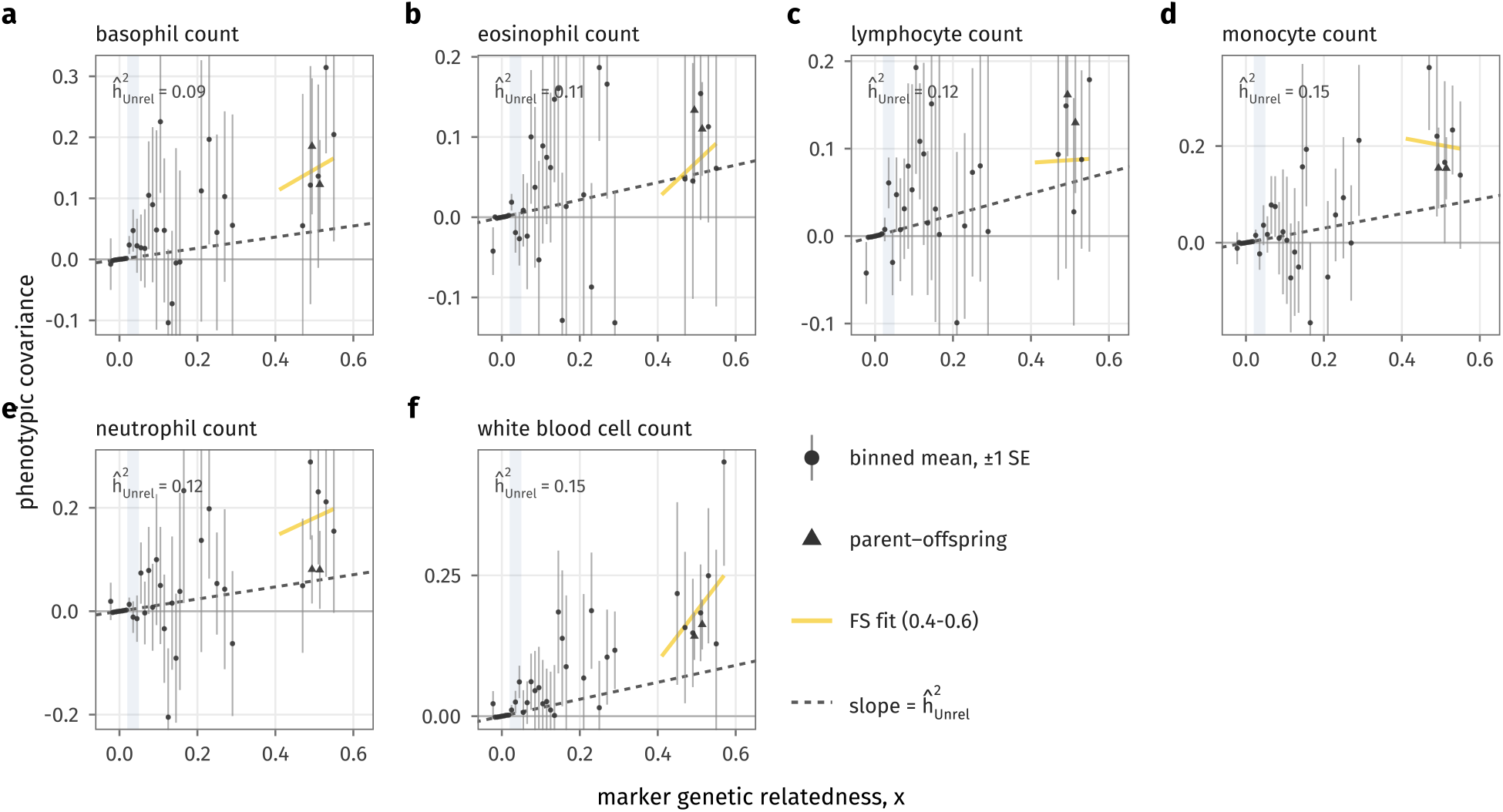
As Figure S17, for white blood cell traits.

**Figure S20:**
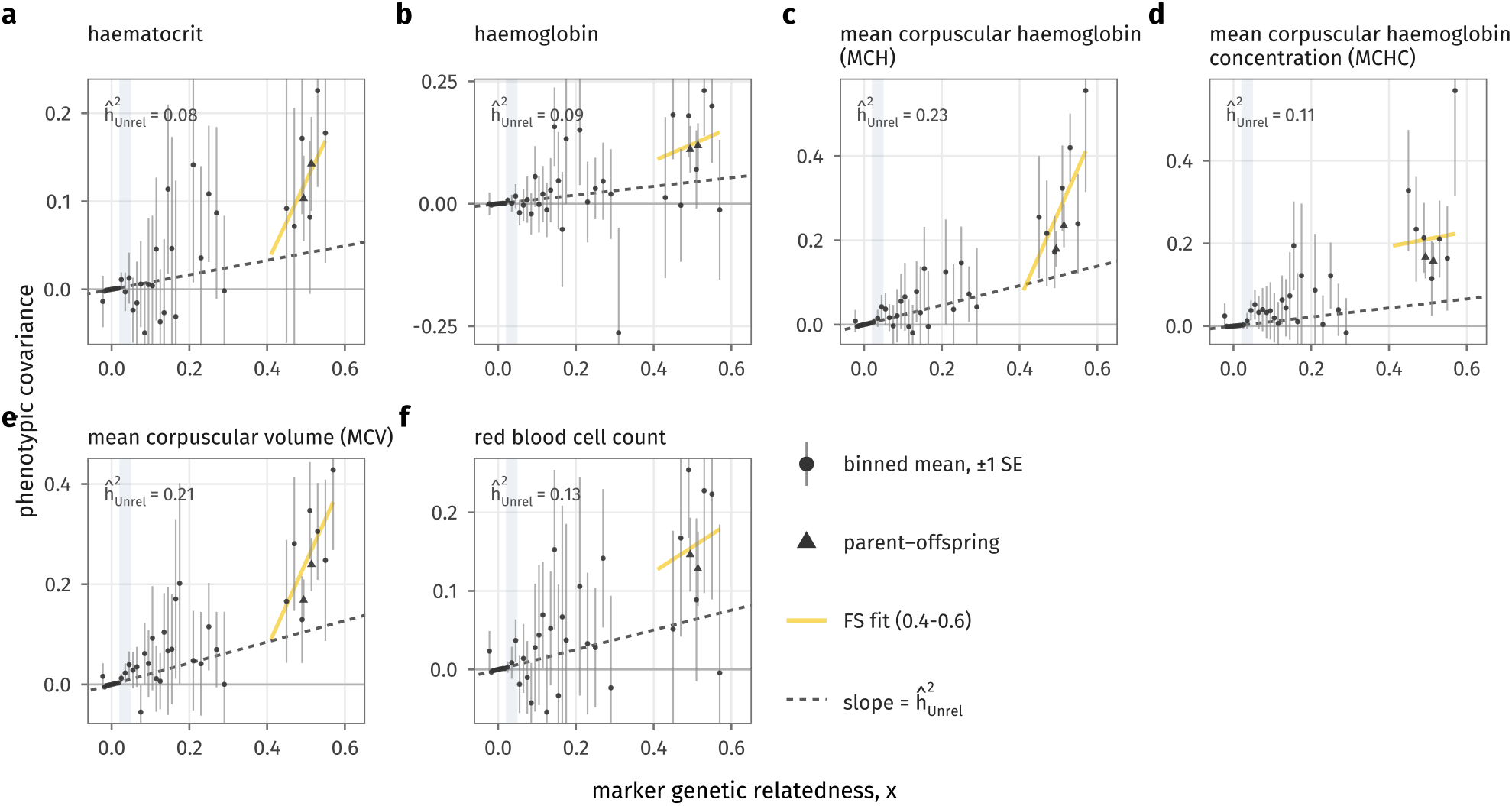
As Figure S17, for red blood cell traits.

**Figure S21:**
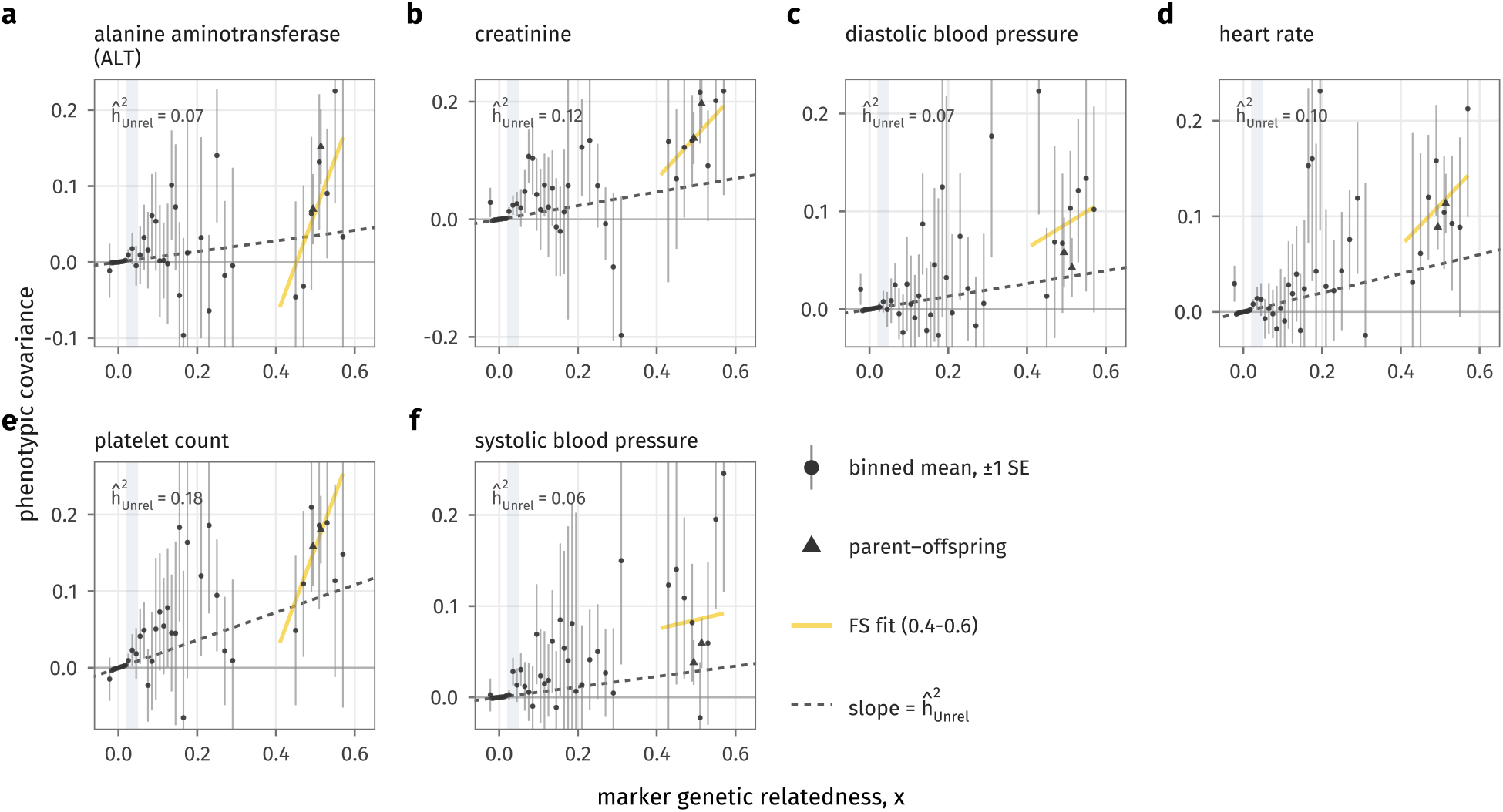
As Figure S17, for platelet, liver, kidney and cardiovascular traits.

**Figure S22:**
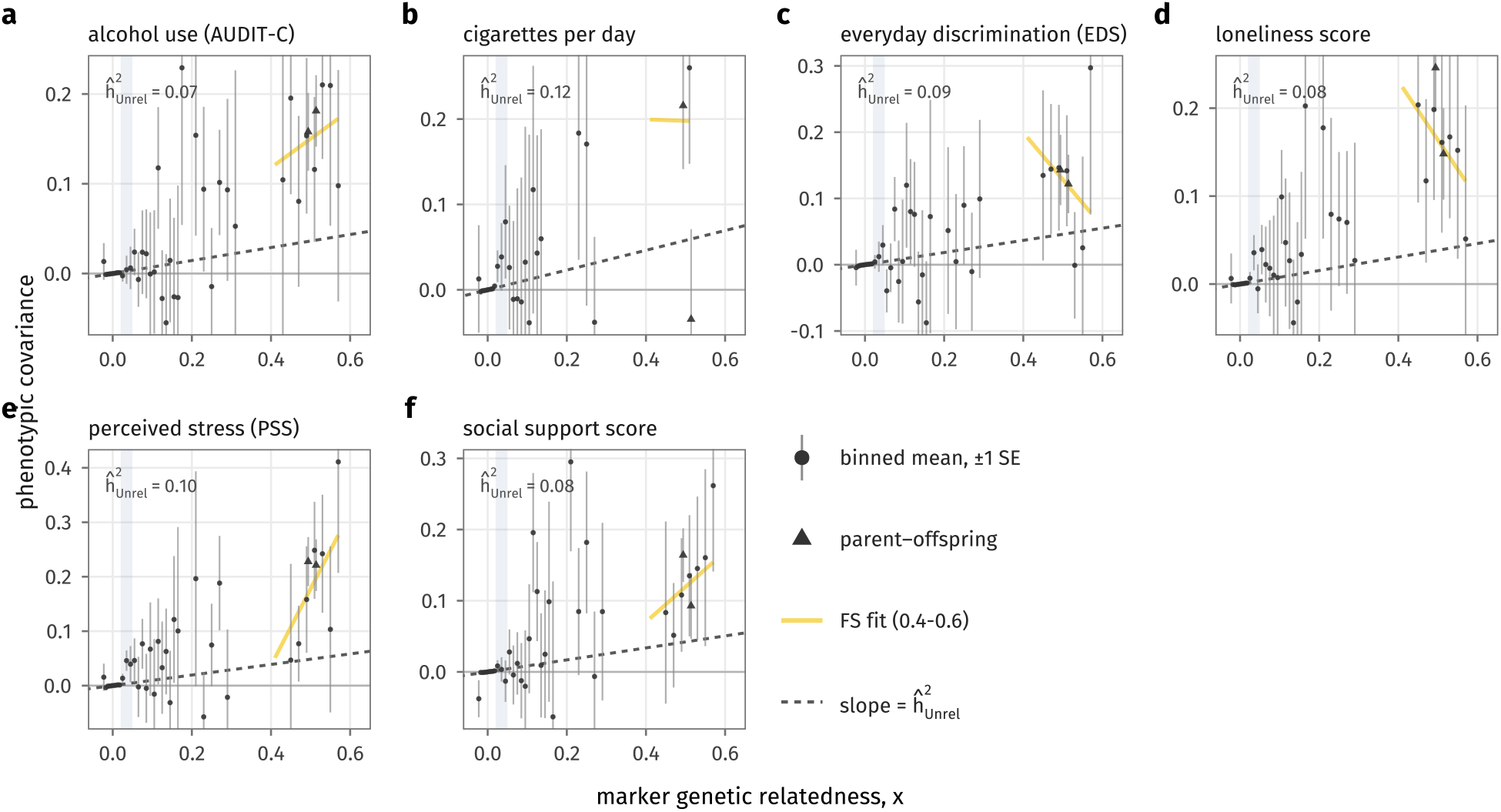
As Figure S17, for lifestyle and psychosocial traits.

